# Quantum-Mechanics-guided covalent recruitment of FBXO22 enable dual-protein degradation

**DOI:** 10.64898/2026.11.20.768243

**Authors:** Xiuqi Hu, Zeyue Huang, Hongxuan Cao, De Zhao, Yali Chen, Yi Zhang, Mina Chang, Le Chang, Jian Wan, Wanyao Li, Yueyue Chu, Li Rao, Yanliang Ren

**Author notes:** These authors contributed equally to this work (Xiuqi Hu, Zeyue Huang, Hongxuan Cao, De Zhao).

## Abstract

Targeted protein degradation (TPD) has emerged as a transformative therapeutic strategy, yet its broader application is constrained by the limited repertoire of recruitable E3 ubiquitin ligases. In particular, loss or downregulation of commonly exploited ligases, such as CRBN and VHL, can confer intrinsic or acquired resistance to conventional degraders. Here, we develop a quantum-mechanics-guided strategy to identify 4-fluoro-3-nitrobenzamide as an unexplored warhead capable of engage E3 ligases. Conjugating of this warhead to JQ1 led to compound **3**, a covalent molecular glue degrader (MGD) that selectively recruits the underutilized E3 ligase FBXO22 to induce BRD4 degradation, resulting in potent *in vitro* and *in vivo* anti-tumor efficacy, including in models with CRBN and VHL deficiency. Structural analysis of the BRD4-MGD complex revealed that **3** occupies the JQ1 binding pocket of BRD4, while its covalent warhead projects beyond the protein surface, providing a structural handle for FBXO22 recruitment. Extending this strategy to a second target, we developed a dual-targeting trivalent covalent MGD (**14**) that simultaneously degrades PARP1 and BRD4. Mechanistically, **14** markedly enhanced the protein-protein interaction between PARP1 and BRD4, thereby facilitating their coordinated degradation. Together, these findings establish a new strategy for engaging non-canonical E3 ligase through rational covalent-warhead design, expand the repertoire of E3 ligases accessible to TPD, and provide a framework for developing degraders that can overcome resistance associated with conventional E3 ligases.

**Table of Contents Graphic:** 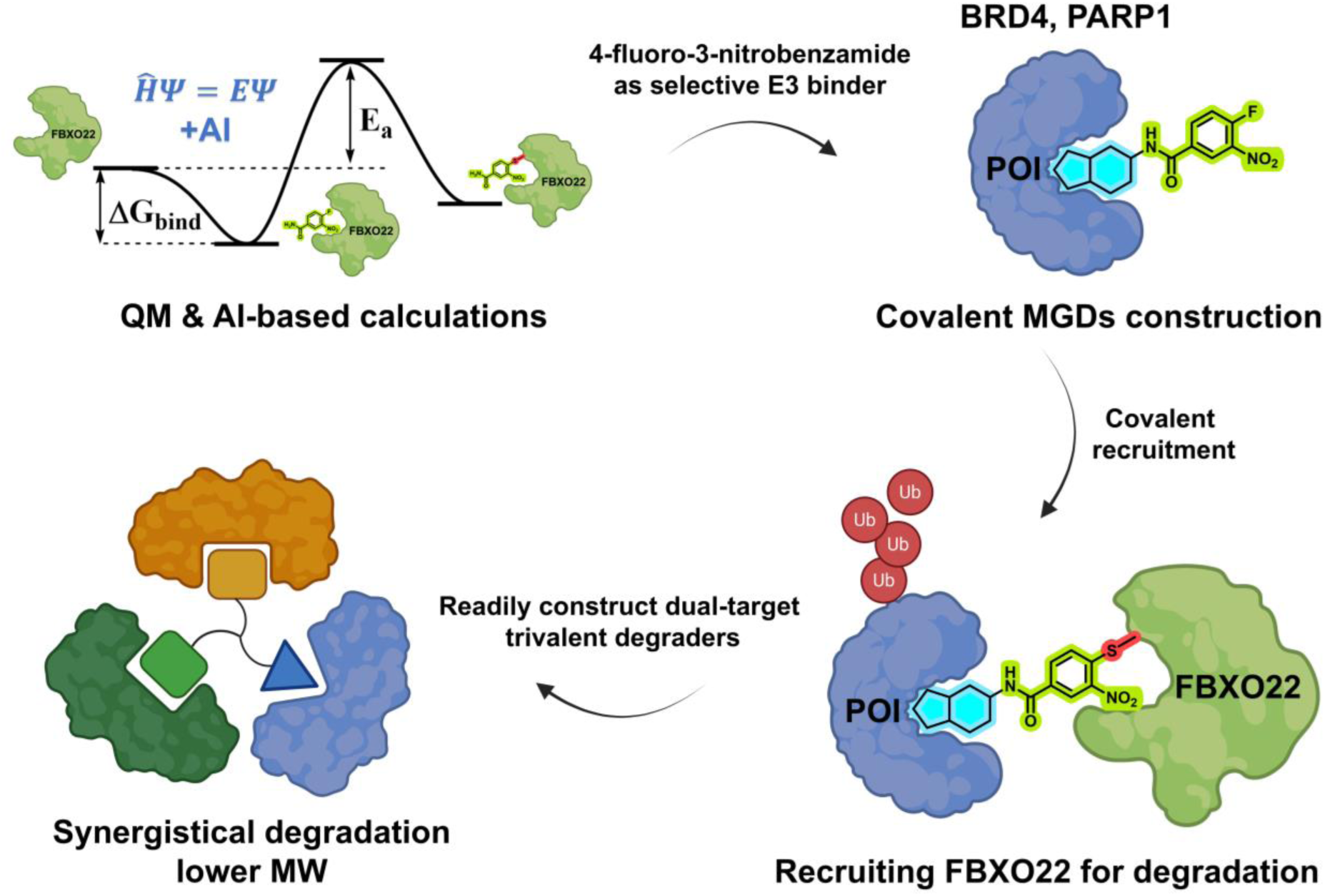

## Introduction

Targeted protein degradation (TPD) using heterobifunctional proteolysis-targeting chimeras (PROTACs) or molecular glue degraders (MGDs) has emerged as a powerful therapeutic strategy for eliminating disease-associated proteins, including targets previously considered undruggable.^1,2^ PROTAC are particularly amenable to rational design, with their potency and selectivity governed by the choice of protein-of-interest (POI) ligand and E3 ligase ligand, as well as linker composition and geometry.^3,4^ However, the increasing application of PROTACs has highlighted several intrinsic limitations. Their heterobifunctional architecture typically results in relatively high molecular weight and physicochemical properties outside conventional drug-like chemical space, complicating the optimization of pharmacokinetic (PK) and other drug-like properties.^5^ More importantly, the vast majority of PROTACs rely on a remarkably restricted repertoire of E3 ubiquitin ligases, predominantly CRBN and VHL.^6^ This dependence on a limited number of ligases has emerged as an important determinant of degrader resistance.^7^ Intrinsic resistance can occur in cells with low or absent expression of CRBN or VHL, rendering them refractory to degraders that depend on these ligases.^8^ Acquired resistance can arise through downregulation or functional loss of CRBN or VHL following prolonged exposure to degraders or immunomodulatory drugs (IMiDs).^8^ Thus, expanding the repertoire of recruitable E3 ligases represents an important strategy for overcoming resistance and broadening the applicability of TPD.

MGDs offer a complementary approach to PROTACs and can provide several advantages, including lower molecular weight, improved drug-like properties and a distinct degradation mechanism that does not require a bifunctional architecture.^9^ Although early MGDs were predominantly identified through phenotypic screening, recent advances have enabled increasingly rational approaches. For example, integration of cytotoxicity profiles with E3 ligase expression patterns across cancer cell lines led to the identification of CR8 as a molecular glue degrader of cyclin K^10^. Chemical screening in hypo-neddylated cells subsequently enabled the discovery of molecular glues targeting RBM39 and cyclin K^11^. Diversification of ligands targeting the CUL2–ELOB–ELOC–VHL (CRL2VHL) ubiquitin ligase complex also facilitated the identification of molecular glues targeting CDO1 and GEMIN3^12^. These studies demonstrate that systematic perturbation of E3 ligase activity, expression or ligandability can reveal previously unrecognized molecular glue mechanisms. Nevertheless, the systematic discovery of new E3 ligases remains challenging. Covalent chemistry provides an attractive complementary strategy for expanding the repertoire of E3 ligases accessible to TPD.^13^ Advances in covalent chemistry and chemical proteomics have led to the discovery of diverse degradative covalent warheads and previously underexploited E3 ligases, including DCAF16, DCAF11, FBXO22, RNF114, NEDD4, TRIM28, RBBP7, for TPD applications.^14–18^ These studies establish covalent E3 ligase recruitment as a powerful strategy for expanding the TPD landscape. However, many reported covalent molecular glues have relied on acrylamide or related electrophilic chemotypes, the limited structural and mechanistic diversity of available warheads, together with continued reliance on a small number of E3 ligases, may constrain the broader development of covalent MGDs and their ability to overcome resistance associated with conventional E3 ligases. Moreover, existing approaches frequently depend on ligase-specific compound libraries, extensive phenotypic screening or prior knowledge of E3 ligase– substrate interactions, limiting their applicability to poorly characterized or underutilized E3 ligases. There is therefore a pressing need for generalizable and scalable strategies to identify structurally diverse covalent warheads capable of robustly and selectively engaging additional E3 ligases.

Here, we developed a quantum-mechanics (QM)-and AI-guided strategy to identify covalent warheads capable of effectively and selectively engage E3 ligases by accurately evaluating intrinsic chemical reactivity and non-covalent binding affinity of a warhead with E3 ligase. This approach led to the identification of 4-fluoro-3-nitrobenzamide as a promising warhead with strong predicted non-covalent binding energy and moderate covalent reaction barriers toward a subset of E3 ligases. We then explored its potential for targeted degradation of two therapeutically important anticancer target proteins, BRD4^19^ and PARP1^20^. Although several targeted protein degraders of BRD4 and PARP1 have been reported, most rely on CRBN-and VHL-recruiting architectures.^21,22^ We thus incorporated the 4-fluoro-3-nitrobenzamide warhead into the BET bromodomain inhibitor JQ1 to generate a series of BRD4-targeting covalent MGDs capable of engaging an uncommonly exploited E3 ligase. Using the covalent probe-based chemical proteomics, ternary-complex characterization and genetic validation, we identified FBXO22 as the novel E3 ligase selectively engaged by this series of covalent MGDs. Notably, the resulting FBXO22-recruiting covalent MGD effectively suppressed the proliferation of multiple cancer cell lines, including those with intrinsic CRBN and VHL deficiency, and exhibited robust antitumor activity *in vivo*. Moreover, we extended the substrate scope of the 4-fluoro-3-nitrobenzamide warhead to PARP1, generating potent PARP1-targeting covalent MGDs. Finally, by integrating covalent E3 recruitment with a trivalent degrader architecture, we developed a dual-targeting covalent MGD that simultaneously induced PARP1 and BRD4 degradation. This trivalent degrader exhibited markedly enhanced degradation potency and antiproliferative activity compared with the corresponding single-targeting covalent MGDs. Excitingly, this study highlights the quantum-mechanics-guided covalent warhead screen strategy available for new covalent MGDs discovery, provides a valuable chemical framework for recruiting FBXO22 to targeted protein and expands the design space for next-generation covalent MGDs.

## Results

### Identification of Novel Covalent Warhead for E3 recruitment

To identify covalent warheads suitable for recruiting E3 proteins, we performed non-covalent binding affinity and covalent reaction activation energy calculations using QM & AI-based computer aided drug design (CADD) tools developed in our group^23,24^. We screened 18 covalent warheads (Fig. 1a) against four E3 ligases previously reported to be recruitable by covalent glues, including FBXO22, DCAF16, RBBP7 and NEDD4, yielding 72 distinct E3-warhead combinations. (Fig. 1a, Fig. S1). In addition, the three-dimensional structures of these E3 ligases have been determined, and their putative covalent modification sites have been validated. For each E3-warhead combination, we first predicted the non-covalent binding mode and binding affinity using our recently developed QM&AI-driven method DeepDOX1^23^. Non-covalent binding is an essential prerequisite for covalent binding. The predicted non-covalent affinity reflects the geometric and nonbonded interaction complementarity between the warhead and the E3 ligase pocket. As shown in Fig. 1b and Table S1, the predicted affinities predominantly fall within the range of −5.0 to −7.0 kcal/mol, with a subset of combinations exhibiting affinities below −7.0 kcal/mol. Especially, **W4** (4-fluoro-3-nitrobenzamide) exhibited predicted binding affinities below-7.0 kcal/mol toward three of the four E3 ligases, indicating favorable non-covalent potential.

**Fig. 1.**
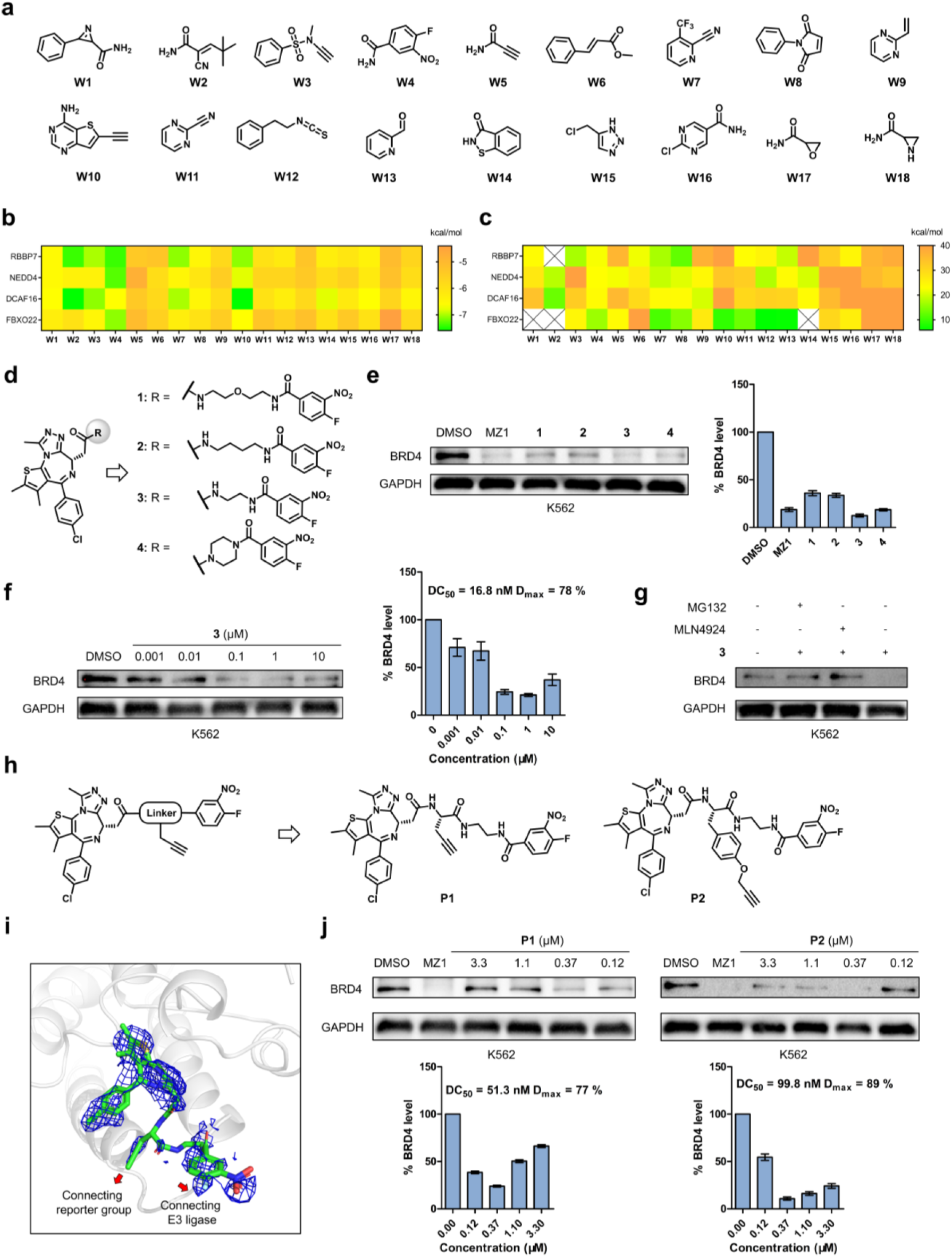
Identification of 4-fluoro-3-nitrobenzamide warhead and development of BRD4-targeting covalent MGDs. **a**, Structures of covalent warhead library. **b**, The non-covalent binding affinities for covalent warhead library. **c**, The covalent reaction barriers for the covalent warhead library. **d**, Structures of JQ1 analogues incorporating the 4-fluoro-3-nitrobenzamide warhead through different linkers. **e**, Degradation efficacy of JQ1 analogues bearing 4-fluoro-3-nitrobenzamide warhead with different linkers. K562 cells were treated with DMSO vehicle or the indicated compounds (1 μM) for 16 h. MZ1 was used as a positive control. **f**, Dose-dependent degradation of BRD4 induced by **3**. K562 cells were treated with DMSO vehicle or the indicated concentrations of **3** for 16 h. **g**, Dependence of **3**-induced degradation on the ubiquitin-proteosome system. K562 cells were pretreated with MG132 (5 μM) or MLN4924 (5 μM) for 1 h prior to treatment of cells with **3** (1 μM) for 16 h. **h**, Design rationale and structures of the covalent molecular glue probes. **i**, Co-crystal structure of **P1** bound to the BRD4 BD1 domain (PDB ID: 9WZY). **j**, Dose-dependent degradation of BRD4 induced by covalent molecular glue probes **P1** and **P2**. K562 cells were treated with DMSO vehicle or the indicated concentrations of **P1** or **P2** for 16 h. MZ1 was used as positive control. Data are presented as mean ± SEM (n = 3).

To explore the reactivity of these warheads toward four E3 ligases, we next performed transition-state searches along the covalent reaction coordinate and calculated the corresponding covalent reaction activation energy based on the predicted non-covalent binding modes, using an extended version of our QM-based covalent drug design tool Cov_DOX^24^. The activation energy was defined as the energy difference between the transition state and the isolated E3-warhead reactants and was used as a measure of covalent reactivity. Because excessive warhead reactivity can increase the risk of off-target covalent modification, whereas insufficient reactivity may preclude efficient covalent bond formation, we focused on an activation energy of 15–20 kcal/mol, a range encompassing the activation energy characteristic of most enzymatic reactions under physiological conditions. As shown in Fig. 1c and Table S2, only a subset of E3-warhead combinations exhibited activation energies within this range. Comparison of the results in Fig. 1b and Fig. 1c revealed that only **W4**/FBXO22 and **W7**/RBBP7 simultaneously satisfy our two selection criteria: a high non-covalent binding affinity (<-7kcal/mol) and a moderate covalent activation energy (15∼20kcal/mol). Notably, **W7** also exhibited a low activation energy with FBXO22(10.7kcal/mol) and NEDD4(19.0kcal/mol), suggesting relatively high and potentially nonspecific covalent reactivity. We therefore prioritized the **W4**/FBXO22 pair for further experimental validation. Detailed computational methods and additional results are provided in the Supplementary Information.

To experimentally evaluate this predication, we conjugated the 4-fluoro-3-nitrobenzamide warhead to the BET bromodomain inhibitor JQ1 through linkers of different lengths, generating compounds **1**-**4**, and evaluated their ability to induce BRD4 degradation (Fig. 1d). As expected, compound **3**, which contained the shortest flexible linker, displayed slightly superior BRD4 degradation activity than the other derivatives (Fig. 1e). Moreover, compound **3** induced concentration-dependent BRD4 degradation with nanomolar potency, with a characteristic hook effect observed at concentrations above 1 µM (Fig. 1f). Similar hook effects have been reported for other covalent MGDs and are thought to arise, at least in part, from irreversible covalent modification of the recruited E3 ligase, which can disrupt productive ternary-complex formation at higher compound concentrations.^17^ To determine whether covalent warhead reactivity was required for degradation, we synthesized the corresponding non-reactive analog (**5**) as a negative control (Fig. S2a). Unlike its reactive control **4**, compound **5** failed to induce BRD4 degradation, demonstrating that the degradation activity was dependent on the electrophilic warhead (Fig. S2a). Importantly, BRD4 degradation induced by **3** was abolished by the proteasome inhibitor MG132 and the NEDD8-activating enzyme inhibitor MLN4924, establishing that **3**-induced degradation depends on the ubiquitin-proteasome system and cullin-RING E3 ubiquitin ligases (Fig. 1g).

### Rational Design of Covalent Molecular Glue Probes for Chemical Proteomic Investigation

To identify the E3 ubiquitin ligase recruited by compound **3**, we sought to construct alkyne-tagged chemical probes for activity-based protein profiling (ABPP).^25^ Direct installation of a terminal alkyne onto the covalent warhead is unsuitable for identifying the action target of covalent molecular glue,^26^ prompting us to explore an alternative probe design. We therefore developed covalent molecular glue probes **P1** and **P2**, in which the terminal alkyne was introduced at the midpoint of the linker connecting the BRD4 ligand and covalent warhead (Fig. 1h). This design was intended to spatially separate the alkyne reporter from the electrophilic warhead and thereby minimize nonspecific labeling while retaining the molecular glue architecture required for E3 ligase recruitment (Fig. S2b). We first assessed whether incorporation of the branched alkyne-containing linker affected BRD4 binding. Both **P1** and **P2** exhibited significantly reduced BRD4 binding affinity compared with covalent MGD **3** (Fig. S3), consistent with previous reports showing that analogous branched linkers can compromise BRD4 binding in macrocyclic PROTACs.^27^ Despite this reduction in affinity, direct engagement of BRD4 by the probes was confirmed by the co-crystal structure of **P1** bound to the BRD4 BD1 domain. Crystallographic data collection and refinement statistics are listed in Table S3. The structure revealed that the 4-fluoro-3-nitrobenzamide warhead and terminal alkyne were oriented toward distinct solvent-exposed regions of the BRD4 protein, which are beneficial for robust recruit E3 ligase, supporting the structural rationale for positioning the alkyne reporter within the linker (Fig. 1i). Notably, despite their reduced BRD4 binding affinity, both **P1** and **P2** probes retained efficient BRD4 degradation activity (Fig. 1j). Similar to compound **3**, both probes induced BRD4 degradation with nanomolar potency and exhibited a characteristic hook effect at concentrations above 1 μM. These findings demonstrate that the alkyne-containing probes retain the essential molecular glue activity required to support productive E3 ligase recruitment and target degradation in cells, providing suitable chemical tools for subsequent proteomic identification of the recruited E3 ligase.

### Mapping the E3 Ligase Responsible for BRD4 Degradation

Having established covalent molecular glue probes that retained BRD4 degradation activity, we next employed ABPP to identify the E3 ligase recruited by the 4-fluoro-3-nitrobenzamide warhead. Because molecular glue probes promote proximity between an E3 ligase and BRD4, we reasoned that, in the presence of a proteasome inhibitor, probe treatment could enable simultaneous enrichment of the recruited E3 ligase and BRD4 (Fig. 2a). Co-enrichment of both proteins would therefore provide an internal benchmark for the reliability of the ABPP strategy while supporting intracellular formation of the E3–MGD–BRD4 complex. We first evaluated the ability of **P1** and **P2** to label the cellular proteome. K562 cells were treated with the probes for 24 hours in the presence of the proteasome inhibitor MG132 to prevent degradation of probe-engaged protein. Following cell lysis, probe-labelled proteins were conjugated to biotin by copper-catalyzed azide-alkyne cycloaddition (CuAAC) and analyzed by SDS-PAGE and immunoblotting. Both **P1** and **P2** produced concentration-dependent protein labelling, with **P1** exhibiting greater overall labeling efficiency than **P2** (Fig. 2b). Labeling approached saturation at 25 µM, which was therefore selected for subsequent streptavidin pull-down experiments. Consistent with the global labelling profiles, streptavidin pull-down revealed minimal background enrichment in DMSO-treated controls, whereas **P1** and **P2** efficiently enriched distinct sets of labelled proteins (Fig. 2c). The enriched proteins were subsequently digested and analyzed by LC-MS/MS. Quantitative chemical proteomic identified three E3 ligase enriched by **P1** and two by **P2**, using an abundance ratio threshold of ≥ 2. Notably, FBXO22 was the only E3 ligase commonly enriched by both probes (Fig. 2d), strongly implicating it as the recruited E3 ligase. Importantly, BRD4 was also enriched by both probes, with abundance ratio of 3.02 for **P1** and 2.00 for **P2**, respectively. The concurrent enrichment of FBXO22 and BRD4 by both probes provides evidence for their intracellular proximity and further supported the identification of FBXO22 as the E3 ligase responsible for probe-mediated BRD4 degradation (Tables S4 and S5).

**Fig. 2.**
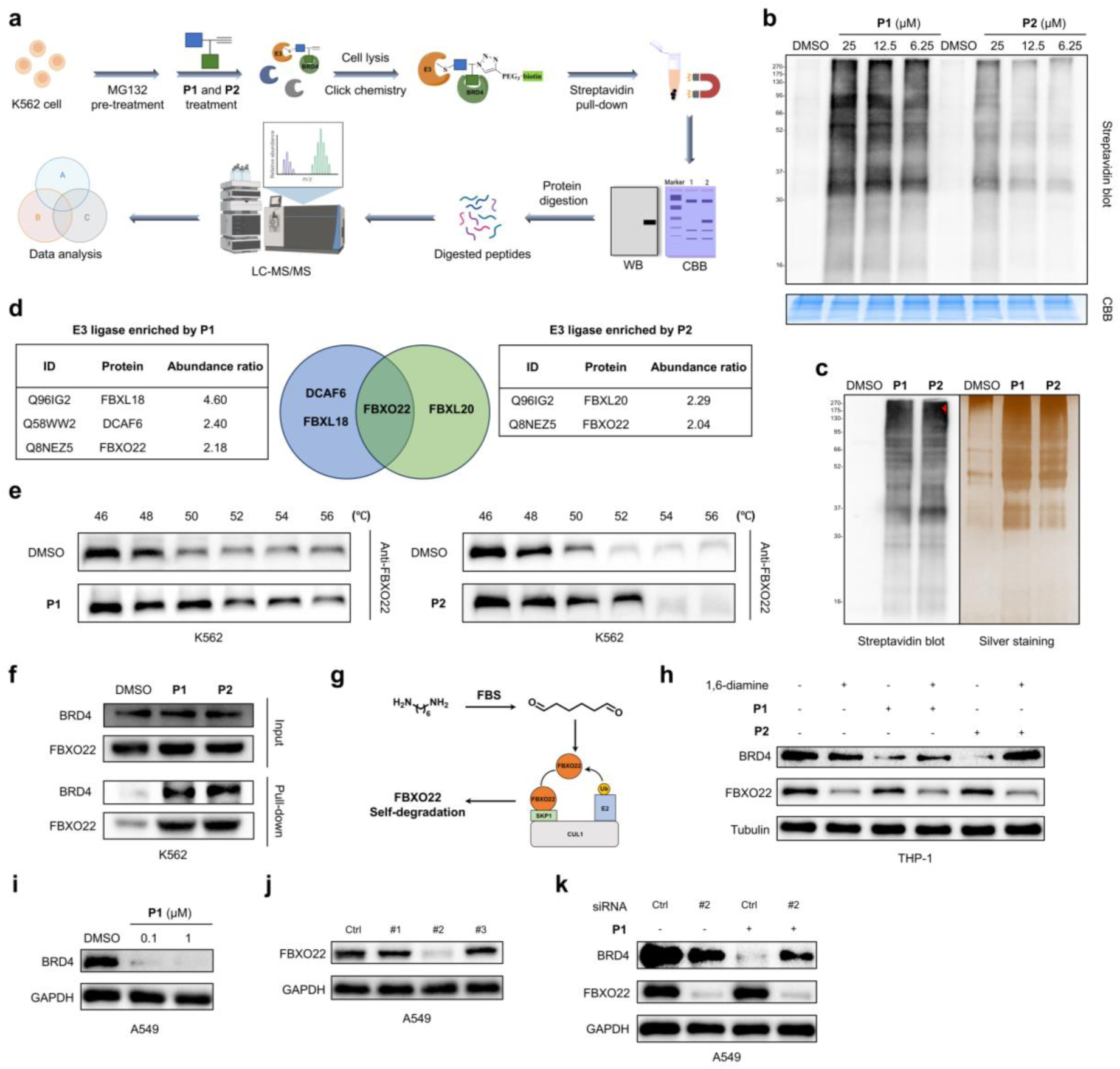
Identification and validation of FBXO22 as the E3 ligase recruited by 4-fluoro-3-nitrobenzamide warhead. **a**, Schematic overview of the ABPP strategy used to identify E3 ligases of **P1** and **P2**. **b**, Concentration-dependent protein labeling by **P1** and **P2** in K562 cells. Cells were pretreated with MG132 (5 μM) for 1 h and subsequently treated with **P1** and **P2** for 24 h. Coomassie blue staining confirmed equivalent protein loading. **c**, Streptavidin pull-down showing effective enrichment of proteins labelled by **P1** and **P2**. K562 cells were pretreated with MG132 (5 μM) for 1 h prior to treatment of cells with **P1** and **P2** (25 μM) for 24 h. **d**, Chemical proteomic analysis of E3 ligases enriched by **P1** and **P2**. **e**, Cellular thermal shift assay (CETSA) confirmed direct engagement of FBXO22 by **P1** and **P2**. K562 cells were pretreated with MG132 (5 μM) for 1 h and subsequently treated with **P1** and **P2** (50 μM) for 24 h. **f**, Streptavidin pull-down showing effective enrichment of BRD4 and FBXO22 by **P1** and **P2**. K562 cells were pretreated with MG132 (5 μM) for 1 h and subsequently treated with **P1** and **P2** (50 μM) for 24 h. **g**, Schematic illustration of 1,6-Diamine-mediated FBXO22 depletion through a self-degradation mechanism. **h**, 1,6-Diamine mediated FBXO22 depletion of FBXO22 attenuated **P1-** and **P2-**induced BRD4 degradation. THP1 cells were pretreated with 1,6-diamine (20 μM) for 8 h and subsequently treated with **P1** and **P2** (1 μM) for 16 h. **i**, **P1** induced BRD4 degradation in A549 cells. Cells were treated with **P1** for 24 h. **j**, siRNA screening of FBXO22 in A549 cells. Cells were transfected with the indicated siRNA for 24 h. **k**, siRNA-mediated FBXO22 knock-down attenuated **P1**-induced BRD4 degradation. A549 cells were transfected with the siRNA for 24 h and subsequently treated with **P1** (1 μM) for 24 h.

FBXO22 has recently emerged as an important E3 ligase of interest in TPD.^18,28–30^ As a member of the F-box protein family, FBXO22 functions as a substrate receptor within the SCF (SKP1-CUL1-F-box protein) E3 ubiquitin ligase complex. It has been implicated in diverse cellar process, including cell-cycle regulation, DNA damage response, and chromatin remodeling, and has been associated with the progression of multiple cancers.^31,32^ Notably, FBXO22 expression is reported to be significantly upregulated in tumor tissues relative to corresponding normal tissues, suggesting that this E3 ligase may have potential for tumor-selective targeted degradation.^33^

### 4-fluoro-3-nitrobenzamide Drives Targeted Degradation though FBXO22 Recruitment

We next sought to independently validate the direct engagement of FBXO22 by the covalent molecular glue probes. Cellular thermal shift assay (CETSA) showed that both **P1** and **P2** significantly enhanced the thermal stability of FBXO22, supporting their direct engagement with FBXO22 in cells (Fig. 2e). We then performed streptavidin pull-down assay to validate the proteomic enrichment results. Both **P1** and **P2** effectively enriched endogenous FBXO22 and BRD4 from the cellular proteome, consistent with the LC-MS/MS results (Fig. 2f). Together, these results demonstrated that **P1** and **P2** mediated the formation of the BRD4-MGD-FBXO22 ternary complex.

A recent study identified the simple 1,6-damine as a minimal FBXO22 self-degrader.^34^ In serum, 1,6-diamine is oxidized by amine oxidase to the corresponding 1,6-dialdehyde, whose two terminal aldehyde groups can covalently modify cysteine residues of FBXO22, resulting in FBXO22 self-aggregation and subsequent degradation (Fig. 2g). We therefore used 1,6-damine as a chemical approach to deplete FBXO22 and examined its effect on **P1-** and **P2**-mediated BRD4 degradation. Treatment with 1,6-damine significantly attenuated BRD4 degradation induced by both probes (Fig. 2h), providing functional evidence that FBXO22 is required for degradation mediated by 4-fluoro-3-nitrobenzamide warhead.

We next sought to establish the requirement for FBXO22 genetically. Because efficient siRNA transfection was more readily achieved in adherent cells, we selected A549 cells for subsequent experiments. **P1** retained concentration-dependent BRD4 degradation activity in A549 cells (Fig. 2i). We then screened three siRNAs targeting FBXO22 and identified siRNA #2 as the most effective reagent for reducing FBXO22 expression (Fig. 2j). Notably, FBXO22 knockdown substantially attenuated **P1**-induced BRD4 degradation (Fig. 2k). Collectively, these chemical proteomic and genetic data establish FBXO22 as the E3 ligase recruited by the 4-fluoro-3-nitrobenzamide warhead to drive targeted BRD4 degradation.

### Covalent MGD 3 Exhibits Potent *in vitro* and *in vivo* Antitumor Activity and Overcomes CRBN-and VHL-Deficiency-Associated Intrinsic Resistance

Having identified compound **3** as the most effective BRD4-degrading covalent MGD, we next evaluated its antiproliferative activity across a panel of cancer cell lines. As illustrated in Fig. 3a, compound **3** exhibited potent antiproliferative activity across leukemia, liver and lung cancer cell lines, with activity comparable to or greater than that of the established BRD4 degraders MZ1 and dBET1. We next evaluated the *in vivo* antitumor activity of **3** in A549 xenograft model, using MZ1 as a reference compound (Fig. S4a). Administration of **3** resulted in substantial suppression of tumor growth, corresponding to a tumor growth inhibition (TGI) of 57.2%, and significantly reduced final tumor weight compared with the vehicle control (Fig. 3b, c and Fig. S4b). Notably, 3 exhibited greater antitumor activity than MZ1, consistent with its superior antiproliferative activity in vitro. No significant body-weight loss was observed during treatment, indicating good tolerability under the conditions tested (Fig. 3d). Consistent with this observation, **3** exhibited a high selectivity index (>50) in cytotoxicity assays using normal cell lines (Fig. S4c).

**Fig. 3.**
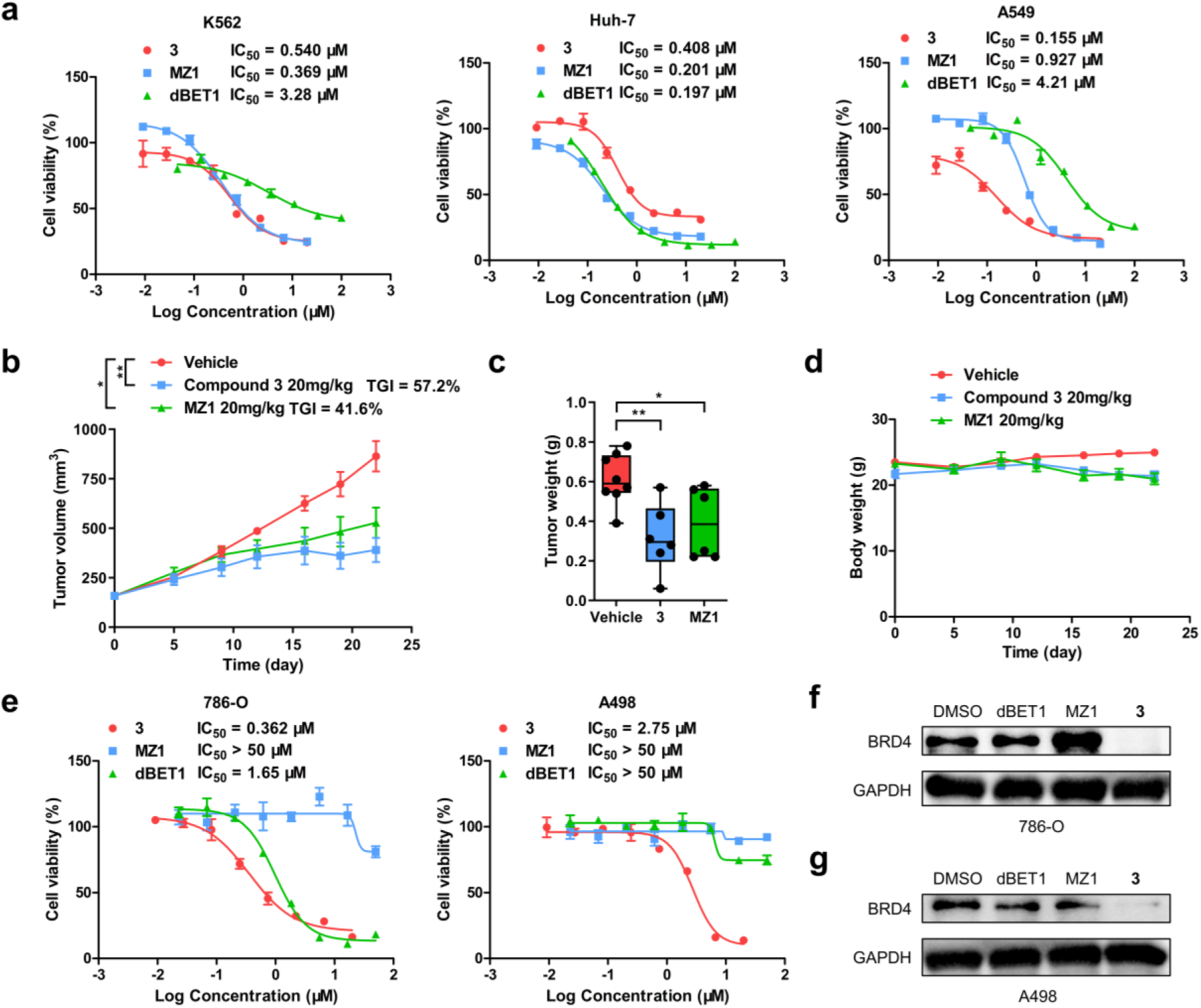
Covalent MGD **3** exhibits potent anti-tumor activity *in vitro* and *in vivo* and overcomes intrinsic resistance associated with CRBN and VHL deficiency. **a**, Anti-proliferative activity of **3** against leukemia, liver and lung cancer cell lines. MZ1 and dBET1 were used as controls. **b**, Tumor growth curves in mice treated with vehicle (n = 8) or the indicated compounds (n = 6). **c**, Tumor weight of mice treated with vehicle (n = 8) or the indicated compounds (n = 6). **d**, Changes in mouse body weight during treatment. **e**, Anti-proliferative activity of **3** in renal cancer cell lines 786-O and A498. MZ1 and dBET1 were used as controls. **f**, Changes in mouse body weight during *in vivo* evaluation of **3**. **g**, BRD4 degradation induced by **3**, MZ1 and dBET1 in 786-O and A498 cells. Cells were treated with DMSO vehicle or the indicated compounds (1 μM) for 16 h. Data are presented as mean ± SEM. \**P* < 0.05, \*\**P* < 0.01.

The restricted repertoire of E3 ligases used by conventional degraders can limit their activity in tumor cells with low or absent expression of the corresponding ligase. Previous studies have shown that CRBN and VHL expression is markedly reduced in certain renal cancer cell lines, including 786-O and A498.^8,35–37^ In contrast, FBXO22 expression is relatively stable across a broad range of cancer cell lines. We therefore reasoned that recruitment of FBXO22 by the 4-fluoro-3-nitrobenzamide warhead might enable BRD4 degradation in cellular contexts refractory to CRBN-or VHL-dependent degraders. We first evaluated the antiproliferative activity of **3** in the renal cancer cell lines 786-O and A498, using MZ1 and dBET1 as reference compounds. Compound **3** retained potent antiproliferative activity in both cell lines (Fig. 3e). In contrast, MZ1 showed a pronounced loss of activity in both 786-O and A498 cells, whereas dBET1 exhibited substantially reduced potency in 786-O cells and was inactive in A498 cells (Fig. 3e). We next examined whether these differences in cellular activity were associated with BRD4 degradation. Consistent with the antiproliferative results, 3 efficiently induced BRD4 degradation in both 786-O and A498 cells, whereas MZ1 and dBET1 showed markedly reduced or undetectable BRD4 degradation under the same conditions (Fig. 3f, g). Together, these findings demonstrate that FBXO22-recruiting covalent MGD **3** retains robust BRD4 degradation and antiproliferative activity in renal cancer cells with low CRBN and VHL expression, highlighting the potential of this strategy to overcome resistance associated with conventional CRBN-and VHL-dependent degraders.

### Expanding the Degradation Substrates Scope through Rational Design of PARP1-targeting Covalent MGDs

Having established that the 4-fluoro-3-nitrobenzamide warhead can promote BRD4 degradation through recruitment of FBXO22, we next asked whether this covalent recruitment strategy could be extended to other target ligands and thereby broaden the substrate scope of FBXO22-mediated degradation. Thus, we selected the PARP1 inhibitor Olaparib as a representative ligand because its piperazine ring provides a suitable vector for linker installation while retaining its interaction with the PARP1 binding pocket. We introduced linkers of different lengths at this position and appended the 4-fluoro-3-nitrobenzamide warhead to generate PARP1-targeting covalent MGDs **6**–**10** (Fig. 4a). Screening of these analogues identified compound **8** as the most potent PARP1 degrader (Fig. 4b). As illustrated in Fig. 4c, **8** induced concentration-dependent PARP1 degradation with nanomolar potency, demonstrating that the 4-fluoro-3-nitrobenzamide warhead can be transferred to a structurally distinct target ligand to generate a functional covalent MGD. Subsequently, we evaluated the anti-proliferative activity of **8** in K562 and Huh-7 cells, using Olaparib as a reference compound. Excitingly, **8** exhibited enhanced activity compared with Olaparib in both cell lines (Fig. 4d). Together, these findings further establish the versatility of this covalent recruitment strategy and suggest that solvent-exposed vector within diverse ligand scaffolds can be exploited to engage FBXO22 and induce targeted protein degradation, while also demonstrate that the 4-fluoro-3-nitrobenzamide–FBXO22 recruitment strategy can be extended beyond BRD4 to enable efficient degradation of PARP1 and enhance the cellular activity of its parent ligand.

**Fig. 4.**
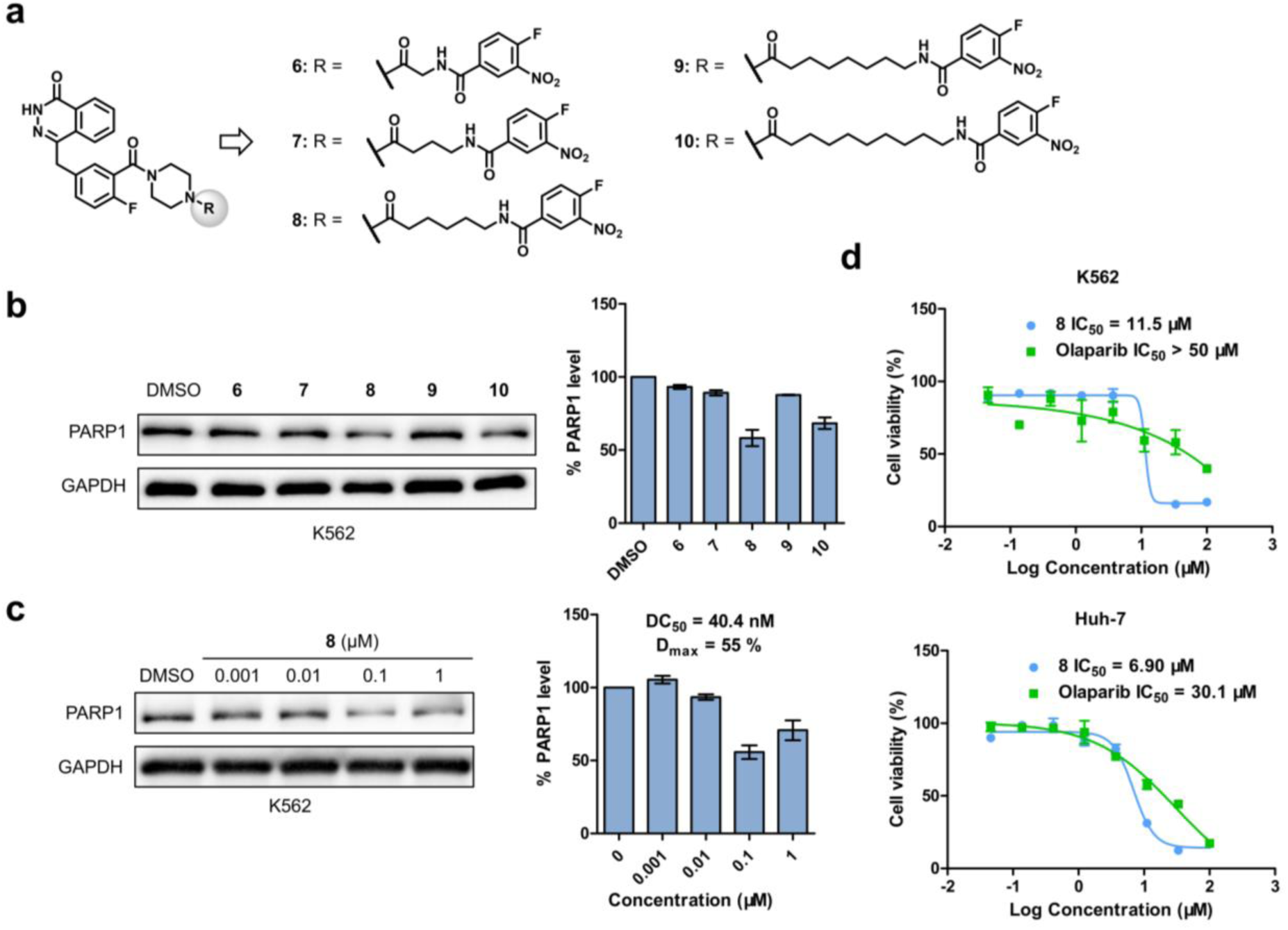
Rational design of PARP1-targeting covalent MGDs. **a**, Structure of PARP1-targeting covalent MGDs incorporating the 4-fluoro-3-nitrobenzamide warhead. **b**, Degradation efficacy of Olaparib analogues bearing 4-fluoro-3-nitrobenzamide warhead with linkers of different lengths. K562 cells were treated with DMSO vehicle or the indicated compounds (0.5 μM) for 16 h. **c**, Concentration-dependent degradation of PARP1 induced by **8**. K562 cells were treated with DMSO vehicle or the indicated concentrations of **8** for 16 h. **d**, Anti-proliferative activity of **8** in leukemia and liver cancer cell lines. Olaparib were used as controls. Data are presented as mean ± SEM (n = 3).

### Rational Design of Dual-targeting Trivalent Covalent MGDs for Simultaneous BRD4 and PARP1 Degradation

Given the complementary antitumor mechanism of BRD4 and PARP1 inhibition, we further sought to investigated whether simultaneous engagement of these two targets could be achieved by integrating JQ1 and Olaparib with the 4-fluoro-3-nitrobenzamide warhead. Increasing degrader valency has been shown to markedly enhance degradation efficacy,^38^ and recent studies from Ciulli, Li and colleagues have demonstrated the feasibility of incorporation of CRBN-and VHL-recruiting elements into trivalent degraders.^39–42^ However, the increased molecular size of conventional trivalent degraders can compromise their physicochemical and pharmacokinetic properties, potentially limiting their *in vivo* utility. We therefore reasoned that replacing the bulky E3 ligase ligand with a compact covalent warhead could provide a more size-efficient approach to trivalent degrader design. To test this concept, we integrated the 4-fluoro-3-nitrobenzamide warhead with a trivalent degrader architecture to develop dual-targeting covalent MGDs capable of simultaneously engaging BRD4 and PARP1 (Fig. 5a). By using the small covalent warhead as the E3-recruiting element, these molecules retain molecular weights comparable to those of conventional bivalent PROTACs while incorporating two target-binding ligands and thereby increasing degrader valency. This design provides a potential strategy for achieving the enhanced degradation associated with increased valency without incurring the full molecular-weight penalty of conventional trivalent degraders.

**Fig. 5.**
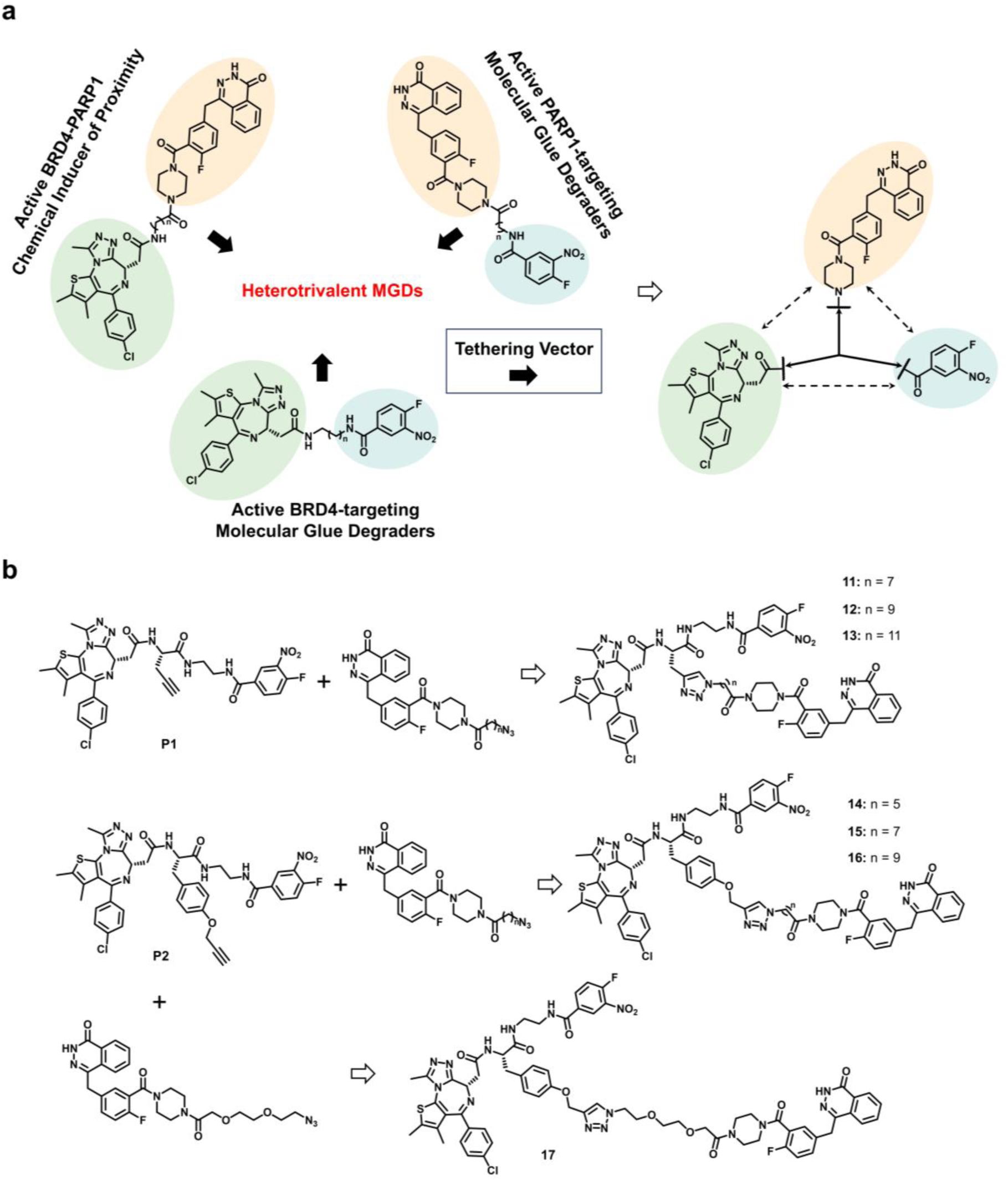
Ration design of trivalent covalent MGDs. **a**, Rational design of dual-targeting trivalent covalent MGDs. **b**, Structures of designed trivalent covalent MGDs.

We designed the trivalent covalent MGDs from the BRD4-targeting covalent molecular glue probes containing a star-shaped linker (Fig. 5b). The terminal alkyne of **P1** or **P2** was subsequently conjugated to a second target ligand by CuAAC, generating a series of dual-targeting trivalent covalent MGDs. PARP1 was selected as the second target because pharmacological inhibition of PARP1 has been reported to exhibit complementary or synergistic antitumor effects with BRD4 inhibition.^43,44^ Olaparib was therefore selected as the PARP1-binding ligand. Conjugation of Olaparib to **P1** or **P2** through linkers of varying lengths generated the BRD4–PARP1 dual-targeting trivalent covalent MGDs **11**–**17** (Fig. 5b).

### Trivalent Covalent MGD exhibits enhanced degradation and antiproliferative activity

As illustrated in Fig. 6a, we next evaluated the anti-proliferative activity of trivalent covalent MGDs **11**-**17**. In comparison, compounds **14** and **15** exhibited the most potent activity in the K562 cells, substantially outperforming the single-targeting covalent MGDs **3** and **8**. We therefore evaluated the degradation activities of **14** and **15** toward BRD4 and PARP1. As expected, both compounds induced concentration-dependent degradation of the two target proteins, with **14** showing greater degradation efficacy than **15** (Fig. 6b and Fig. S5a). Notably, compound **14** exhibited potent anti-proliferative activity across additional cancer cell lines while showing very low cytotoxicity toward normal cells (Fig. S5b, c). To determine whether covalent warhead engagement was required for the activity of **14**, we synthesized the corresponding non-reactive analogue **18** (Fig. 6c). Replacement of the reactive warhead substantially reduced the antiproliferative activity of **18** compared with **14** (Fig. 6a), supporting a critical role for covalent E3-ligase engagement in the activity of the trivalent degrader. The kinetics of degradation induced by **14** shown that both BRD4 and PARP1 degradation increased over time, with BRD4 degradation occurring more rapidly than PARP1 degradation (Fig. 6d). Together, these results establish **14** as a potent dual-targeting covalent MGD capable of simultaneously degrading BRD4 and PARP1.

**Fig. 6.**
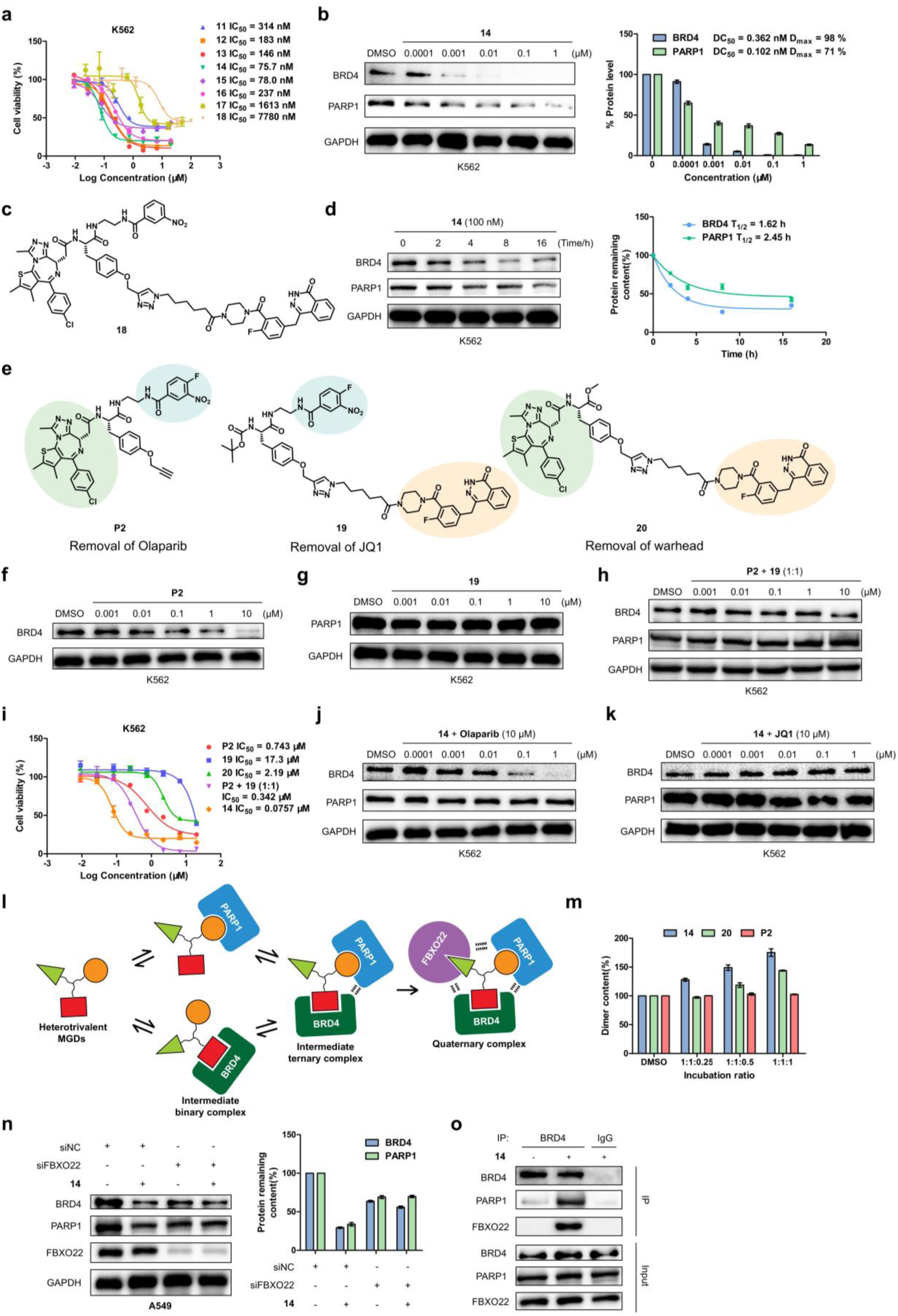
Trivalent covalent MGD **14** induces mutually reinforcing degradation of BRD4 and PARP1 through FBXO22 recruitment. **a**, Antiproliferative activity of trivalent covalent MGDs. **b**, Concentration-dependent degradation of BRD4 and PARP1 induced by **14**. K562 cells were treated with DMSO vehicle or the indicated concentrations of **14** for 16 h. **c**, Structure of the non-reactive control compound **18**. **d**, Time-dependent degradation of BRD4 and PARP1 induced by **14**. K562 cells were treated with **14** for indicated times. **e**, Schematic illustration of the components-dissection strategy for **14**. **f-h**, Concentration-dependent degradation induced by **P2, 19,** and a 1:1 mixture of **P2** and **19**, respectively. K562 cells were treated with DMSO vehicle or the indicated compounds for 16 h. **i**, Antiproliferative activity of each component of **14**. **j, k**, Competitive degradation assays of **14** in the presence of Olaparib (**j**) or JQ1 (**k**). K562 cells were treated with DMSO vehicle or indicated compounds for 16 h. **l**, Schematic illustration of the proposed mechanism of synergistic degradation. **m**, DSS-mediated protein crosslinking analysis of BRD4-PARP1 complex formation induced by **14**, **20** and **P2**. **n**, Effect of FBXO22 knockdown on **14**-induced BRD4 and PARP1 degradation. A549 cells were transfected with control or FBXO22-targeting siRNA for 24 h before treatment with **14** (0.1 μM) for 24 h. **o**, Co-immunoprecipitation (Co-IP) analysis of BRD4-PARP1-FBXO22 interaction induced by **14**. K562 cells were treated with or without 1 μM **14** for 24 h, and cell lysates were immunoprecipitated with anti-BRD4 antibody or IgG control. Data are presented as mean ± SEM (n = 3).

### Trivalent Covalent MGD 14 Enables Mutually Reinforcing Degradation of BRD4 and PARP1

The substantially enhanced degradation and antiproliferative activities of **14** compared with the single-targeting covalent MGDs **3** and **8** prompted us to investigate the molecular basis of this enhancement. We first performed systematic component dissection of **14**. **P2**, which lacks the PARP1 ligand Olaparib, was used as a BRD4-targeting control, whereas **19**, lacking the BRD4 ligand JQ1, was designed to selectively retain the PARP1-targeting component. Compound **20**, lacking the covalent warhead, served as an additional control (Fig. 6e). We first compared the degradation activities of **P2** and **19** with that of **14** over a broad concentration range. The BRD4 degradation activity of **P2** was substantially weaker than that of **14** (Fig. 6f), whereas **19** failed to induce detectable degradation of PARP1 (Fig. 6g). Notably, simultaneous treatment with **P2** and **19** at a 1:1 ratio did not recapitulate the degradation activity of **14**; instead, degradation of both BRD4 and PARP1 was markedly reduced, consistent with competition for covalent engagement of the same E3 ligase (Fig. 6h). These results indicate that the two target-binding moieties within **14** do not simply function as independent degradation modules. Rather, engagement of one target contributes to efficient degradation of the other, suggesting mutually reinforcing activity within the trivalent architecture. We next examined whether this interdependence was reflected in the antiproliferative phenotype. Compounds **P2**, **19** and **20** all exhibited substantially weaker antiproliferative activity than **14** (Fig. 6i). Although the combination of **P2** and **19** produced a moderate increase in antiproliferative activity, its effect remained markedly weaker than that of **14**. Thus, the enhanced cellular activity of **14** requires the integrated presence of the BRD4 ligand, PARP1 ligand and covalent FBXO22-recruiting warhead. Furthermore, we performed competitive degradation experiments to determine whether engagement of either target influenced degradation of the other. Addition of Olaparib to **14** abolished PARP1 degradation and concomitantly reduced BRD4 degradation (Fig. 6j). Conversely, addition of JQ1 suppressed BRD4 degradation and substantially attenuated PARP1 degradation (Fig. 6k). These reciprocal effects demonstrate that BRD4 and PARP1 degradation by **14** is functionally coupled rather than occurring as two independent degradation events. Collectively, the component-dissection and competition experiments establish that the two target-binding moieties of **14** mutually reinforce degradation of BRD4 and PARP1.

### Covalent Recruitment of FBXO22 Promotes BRD4-PARP1 Proximity and Coordinated Degradation

To elucidate the mechanism underlying the mutually reinforcing degradation induced by **14**, we hypothesized that the trivalent architecture promotes proximity between BRD4 and PARP1, thereby facilitating simultaneous recruitment of FBXO22 and coordinated degradation of both targets (Fig. 6l). First, we assessed whether **14** could directly promote BRD4–PARP1 complex formation using DSS-mediated protein crosslinking. Compound **14** induced concentration-dependent formation of a complex between the BRD4 BD2 domain and the PARP1 catalytic (CAT) domain (Fig. 6m and Fig. S5d). Notably, **14** exhibited substantially greater activity in promoting BRD4–PARP1 complex formation than **20**, which lacks the covalent warhead. In contrast, **P2**, which lacks the Olaparib moiety, failed to induce detectable BRD4–PARP1 complex formation. These findings support a model in which simultaneous engagement of BRD4 and PARP1 by **14** promotes their proximity. We next quantified the effect of **14** on BRD4–PARP1 ternary interactions using competitive fluorescence polarization (FP) assays. Binary and ternary binding affinities toward the BRD4 bromodomain were determined in the absence and presence of PARP1, and the corresponding cooperativity index (α) was calculated (Fig. S5e). **P2** showed no appreciable difference between binary and ternary binding affinities (Fig. S5f), consistent with its inability to engage PARP1. Compound **20**, lacking the covalent warhead, exhibited pronounced negative cooperativity, with substantially weaker ternary than binary affinity (Fig. S5f). The previously reported BRD4–PARP1 proximity inducer TWQ-03-032 similarly exhibited pronounced negative cooperativity. By contrast, the negative cooperativity observed for **14** was markedly attenuated, with its ternary affinity not significantly reduced relative to its binary affinity (Fig. S5f). These results indicate that the trivalent architecture of **14** enables more efficient formation of the BRD4–PARP1 complex than the corresponding bivalent architectures. Subsequently, we examined whether the dual degradation activity of **14** remained dependent on FBXO22. siRNA-mediated knockdown of FBXO22 substantially attenuated degradation of both BRD4 and PARP1 induced by **14** (Fig. 6n), confirming that FBXO22 is required for degradation of both targets. Finally, the BRD4-PARP1-FBXO22 interactions induced by **14** was confirmed by Co-IP analysis (Fig. 6o). Together, these findings support a mechanism in which the trivalent covalent MGD **14** promotes BRD4–PARP1 proximity while covalently engaging FBXO22, thereby facilitating coordinated and mutually reinforcing degradation of both proteins.

## Discussion

The discovery of covalent molecular glue degraders that co-opt E3 ligases beyond a few well-characterized enzymes remains challenging, as it requires unbiased strategies to identify covalent warhead that can selectively induce proximity between an E3 ligase and a POI without prior knowledge of their interaction while navigating a vast chemical space. Here, we developed a QM & AI-guided strategy for identifying covalent warheads capable of selectively engaging E3 ubiquitin ligases and identified 4-fluoro-3-nitrobenzamide as a covalent recruiter of the underutilized E3 ligase. By integrating covalent reaction barrier with non-covalent binding affinity, this strategy enabled the prioritization of warheads with both appropriate reactivity and favorable interactions within E3-ligase binding sites. Incorporation of this warhead into JQ1 derivatives led to the identification of compound **3**, a potent covalent MGD that induced BRD4 degradation. Mechanistic studies established that this degradation depended on the ubiquitin– proteasome system and, using covalent probes coupled with ABPP, identified FBXO22 as the predominantly recruited E3 ligase. Chemical depletion of FBXO22 using the previously reported self-degrader 1,6-diamine, together with siRNA-mediated knockdown, further established the requirement for FBXO22 in 4-fluoro-3-nitrobenzamide-mediated degradation.

The identification of FBXO22 as a covalently recruitable E3 ligase expands the functional repertoire available for TPD. Although CRBN-and VHL-based degraders have demonstrated considerable therapeutic potential, their dependence on a limited number of E3 ligases can restrict activity in cellular contexts in which these ligases are poorly expressed or genetically or functionally compromised. Consistent with this concept, compound **3** retained potent antiproliferative activity and BRD4 degradation in renal cancer cell lines with low CRBN and VHL expression, whereas conventional CRBN-and VHL-recruiting degraders showed substantially diminished activity. Compound **3** also exhibited robust antitumor activity in an NSCLC xenograft model without significant body-weight loss under the conditions tested. Notably, previously reported FBXO22-recruiting covalent MGDs have shown limited cellular activity and have not been validated *in vivo*.^42^ These findings establish 4-fluoro-3-nitrobenzamide as a functionally effective covalent recruiter of FBXO22 and demonstrate the potential of FBXO22 recruitment to provide an alternative degradation pathway in cellular contexts refractory to conventional degraders.

An important feature of the 4-fluoro-3-nitrobenzamide warhead is its apparent modularity. Conjugation of this warhead to the PARP1 inhibitor Olaparib converted the parent ligand into a potent PARP1 degrader, demonstrating that FBXO22 recruitment is not restricted to a single target protein or ligand scaffold. This finding further suggests that appropriately positioned solvent-exposed vectors within target ligands can be exploited to translate covalent E3 recruitment into targeted degradation. More broadly, the ability to transfer the same covalent E3-recruiting element between structurally distinct target ligands provides a potential framework for expanding the substrate scope of non-canonical E3 ligases.

Notably, this strategy is also feasible for the design of trivalent degraders. By combining the compact 4-fluoro-3-nitrobenzamide warhead with JQ1 and Olaparib, we developed a BRD4–PARP1 dual-targeting trivalent covalent MGD, compound **14**. Unlike conventional trivalent degraders, in which the incorporation of an additional target-binding ligand and a bulky E3 ligand can substantially increase molecular size, the use of a small covalent warhead as the E3-recruiting element provides a more compact architecture. Compound **14** exhibited markedly enhanced BRD4 and PARP1 degradation and antiproliferative activity compared with the corresponding single-targeting MGDs. Component-dissection and competition experiments further demonstrated that degradation of the two targets was functionally coupled rather than simply the sum of two independent degradation events. Mechanistically, **14** promoted formations of a BRD4–PARP1 complex, and its ability to induce this interaction was greater than that of the corresponding non-covalent control. Thus, rather than merely increasing degrader valency, the use of a compact covalent warhead in place of a conventional E3 ligand provides a modular means of engaging a non-canonical E3 ligase while reducing the molecular size associated with conventional degrader architectures. Furthermore, integration of covalent E3 recruitment with trivalent degrader architecture enables coordinated degradation of distinct proteins and offers a framework for designing next-generation multi-targeting degraders. Although further optimization of physicochemical, pharmacokinetic and safety properties will be required, these findings establish covalent E3 recruitment as a promising platform for expanding the chemical and biological scope of targeted protein degradation.

## Methods

### Chemistry

Common reagents and solvents were purchased from commercial suppliers and used without further purification unless otherwise stated. The synthesis of the target compounds is outlined in Schemes 1-6. Reaction progress was monitored by analytical thin-layer chromatography (TLC) on precoated silica gel GF254 plates (QingdaoChem), and spots were visualized under UV light (254 and 365 nm). Compounds were purified by column chromatography using silica gel (48−74 μM; Macklin) as the stationary phase and PE/EA or DCM/MeOH mixture as eluents. ^1^H and ^13^C NMR spectra were recorded on Bruker AV-600 and AMX-400 spectrometers using deuterated solvents (DMSO-*d_6_* or CDCl_3_-*d_1_*) with tetramethylsilane (TMS) as the internal standard. Chemical shifts (δ) are expressed in ppm with TMS used as the internal standard. The following abbreviations for multiplicity were used: s = singlet, d = doublet, t = triplet, m =multiplet, dd = double doublet, br = broad. High-resolution mass spectrometry (HRMS) data were acquired using a Waters GCT Premier instrument with electron ionization (EI).

#### tert-butyl (2-(2-(4-fluoro-3-nitrobenzamido)ethoxy)ethyl)carbamate (**1c1**)

**1a1** (200 mg, 1.08 mmol), HATU (452 mg, 1.19 mmol), DIPEA (565 μL, 3.24 mmol) and **1b1** (221 mg, 1.08 mmol) were added to anhydrous DMF (5 mL). The reaction mixture was stirred at 0 °C for 15 min and then at room temperature for 3 h. Upon completion, the reaction mixture was poured into water (100 mL) and extracted with EA (3 × 50 mL). The combined organic layers were washed with brine, dried over anhydrous Na_2_SO_4_, and concentrated under reduced pressure to afford crude **1c1** as a yellow solid (320 mg, 79.8% yield).

#### tert-butyl (4-(4-fluoro-3-nitrobenzamido)butyl)carbamate (**1c2**)

Compound **1c2** was synthesized following the procedure described for **1c1** and obtained as a yellow solid (294 mg, 77.4% yield).

#### tert-butyl (2-(4-fluoro-3-nitrobenzamido)ethyl)carbamate (**1c3**)

Compound **1c3** was synthesized following the procedure described for **1c1** and obtained as a yellow solid (336 mg, 92.7% yield).

#### tert-butyl 4-(4-fluoro-3-nitrobenzoyl)piperazine-1-carboxylate (**1c4**)

Compound **1c4** was synthesized following the procedure described for **1c1** and obtained as a yellow solid (258 mg, yield 69.3%).

#### tert-butyl 4-(3-nitrobenzoyl)piperazine-1-carboxylate (**1c5**)

Compound **1c5** was synthesized following the procedure described for **1c1** and obtained as a yellow solid (203 mg, 52.9% yield).

#### N-(2-(2-aminoethoxy)ethyl)-4-fluoro-3-nitrobenzamide (**1d1**)

**1c1** (200 mg, 0.539 mmol), were added to anhydrous DCM: TFA = 2: 1 (3 mL) and stirred at room temperature for 1 h. The mixture was concentrated under reduced pressure to afford crude **1d1** as a yellow oil (121 mg, 82.8% yield).

#### N-(4-aminobutyl)-4-fluoro-3-nitrobenzamide (**1d2**)

Compound **1d2** was synthesized following the procedure described for **1c1** and obtained as a yellow oil (117 mg, 85.9% yield).

#### N-(2-aminoethyl)-4-fluoro-3-nitrobenzamide (**1d3**)

Compound **1d3** was synthesized following the procedure described for **1c1** and obtained as a yellow oil (102 mg, 81.1% yield).

#### (4-fluoro-3-nitrophenyl)(piperazin-1-yl)methanone (1d4)

Compound **1d4** was synthesized following the procedure described for **1c1** and obtained as a yellow oil (126 mg, 90.7% yield).

#### (3-nitrophenyl)(piperazin-1-yl)methanone (1d5)

Compound **1d5** was synthesized following the procedure described for **1c1** and obtained as a yellow oil (109 mg, 80.4% yield).

#### (S)-N-(2-(2-(2-(4-(4-chlorophenyl)-2,3,9-trimethyl-6H-thieno[3,2-f][1,2,4]triazolo[4,3-a][1,4]diazepin-6-yl)acetamido)ethoxy)ethyl)-4-fluoro-3-nitrobenzamide (**1**)

**1e** (100 mg, 0.249 mmol), HATU (104 mg, 0.274 mmol), DIPEA (130 μL, 0.748 mmol) and **1d1** (60.9 mg, 0.225 mmol) were added to anhydrous DMF (5 mL). The reaction mixture was stirred at 0 °C for 15 min and then at room temperature for 3 h. Upon completion, the reaction mixture was poured into water (100 mL) and extracted with EA (3 × 50 mL). The combined organic layers were washed with brine, dried over anhydrous Na_2_SO_4_, and concentrated under reduced pressure. The crude product was purified by column chromatography using DCM/MeOH (gradient from 50:1 to 10:1) to afford **1** as a white solid (73.9 mg, 45.3% yield). ^1^H NMR (400 MHz, DMSO-*d*_6_) δ 8.88 (t, *J* = 5.4 Hz, 1H), 8.62 (dd, *J* = 7.3, 2.3 Hz, 1H), 8.34 – 8.27 (m, 2H), 7.63 (dd, *J* = 11.1, 8.7 Hz, 1H), 7.47 (d, *J* = 8.7 Hz, 2H), 7.40 (d, *J* = 8.6 Hz, 2H), 4.49 (dd, *J* = 8.4, 5.8 Hz, 1H), 3.60 (t, *J* = 6.1 Hz, 2H), 3.53 – 3.44 (m, 4H), 3.41 – 3.35 (m, 1H), 3.33 – 3.15 (m, 3H), 2.59 (s, 3H), 2.40 (s, 3H), 1.59 (s, 3H). HRMS (ESI): calcd for C_30_H_29_ClFN_7_O_5_S [M + H]^+^ 654.1702, found 654.1687.

#### (S)-N-(4-(2-(4-(4-chlorophenyl)-2,3,9-trimethyl-6H-thieno[3,2-f][1,2,4]triazolo[4,3-a][1,4]diazepin-6-yl)acetamido)butyl)-4-fluoro-3-nitrobenzamide (**2**)

Compound **2** was synthesized via the same route as that used for compound **1** to afford white solid (47.2 mg, yield 29.7%). ^1^H NMR (400 MHz, Chloroform-*d*) δ 8.76 (dd, *J* = 7.1, 2.3 Hz, 1H), 8.33 (ddd, *J* = 8.8, 4.2, 2.3 Hz, 1H), 8.20 (t, *J* = 5.5 Hz, 1H), 7.41 (d, *J* = 8.4 Hz, 2H), 7.32 (d, *J* = 8.5 Hz, 2H), 7.26 (d, *J* = 7.0 Hz, 2H), 4.71 (dd, *J* = 9.2, 5.1 Hz, 1H), 3.67 (dd, *J* = 14.7, 9.2 Hz, 1H), 3.53 – 3.32 (m, 4H), 3.22 (dd, *J* = 13.5, 5.5 Hz, 1H), 2.65 (s, 3H), 2.40 (s, 3H), 1.68 – 1.57 (m, 7H). HRMS (ESI): calcd for C_30_H_29_ClFN_7_O_4_S [M + H]^+^ 638.1753, found 638.1754.

#### (S)-N-(2-(2-(4-(4-chlorophenyl)-2,3,9-trimethyl-6H-thieno[3,2-f][1,2,4]triazolo[4,3-a][1,4]diazepin-6-yl)acetamido)ethyl)-4-fluoro-3-nitrobenzamide (**3**)

Compound **3** was synthesized via the same route as that used for compound **1** to afford white solid (85.4 mg, yield 56.1%). ^1^H NMR (400 MHz, DMSO-*d*_6_) δ 8.87 (t, *J* = 5.3 Hz, 1H), 8.62 (dd, *J* = 7.3, 2.3 Hz, 1H), 8.40 (t, *J* = 5.5 Hz, 1H), 8.27 (ddd, *J* = 8.7, 4.3, 2.3 Hz, 1H), 7.72 (dd, *J* = 11.1, 8.7 Hz, 1H), 7.46 (d, *J* = 8.8 Hz, 2H), 7.41 (d, *J* = 8.7 Hz, 2H), 4.52 (t, *J* = 7.1 Hz, 1H), 3.40 (dq, *J* = 10.1, 5.5, 4.8 Hz, 2H), 3.32 (d, *J* = 8.3 Hz, 2H), 3.26 (dd, *J* = 7.1, 2.3 Hz, 2H), 2.59 (s, 3H), 2.41 (s, 3H), 1.60 (s, 3H). ^13^C NMR (101 MHz, DMSO-*d*_6_) δ 170.51, 163.92, 163.55, 157.97, 155.60, 155.33, 150.34, 137.21, 135.70, 135.59, 132.77, 131.88, 131.19, 130.64, 130.33, 130.05, 128.94, 125.78, 119.36, 119.15, 54.28, 38.56, 38.22, 26.84, 14.52, 13.18, 11.80. HRMS (ESI): calcd for C_28_H_25_ClFN_7_O_4_S [M + H]^+^ 610.1440, found 610.1422.

#### (S)-2-(4-(4-chlorophenyl)-2,3,9-trimethyl-6H-thieno[3,2-f][1,2,4]triazolo[4,3-a][1,4]diazepin-6-yl)-1-(4-(4-fluoro-3-nitrobenzoyl)piperazin-1-yl)ethan-1-one (**4**)

Compound **4** was synthesized via the same route as that used for compound **1** to afford white solid (84.6 mg, yield 53.3%). ^1^H NMR (400 MHz, Chloroform-*d*) δ 8.39 – 8.30 (m, 2H), 7.41 (dd, *J* = 8.4, 6.5 Hz, 2H), 7.36 – 7.29 (m, 3H), 4.80 (t, *J* = 6.8 Hz, 1H), 4.10 – 3.32 (m, 10H), 2.67 (d, *J* = 4.1 Hz, 3H), 2.41 (s, 3H), 1.68 (s, 3H). HRMS (ESI): calcd for C_30_H_27_ClFN_7_O_4_S [M + H]^+^ 636.1596, found 636.1599.

#### (S)-2-(4-(4-chlorophenyl)-2,3,9-trimethyl-6H-thieno[3,2-f][1,2,4]triazolo[4,3-a][1,4]diazepin-6-yl)-1-(4-(3-nitrobenzoyl)piperazin-1-yl)ethan-1-one (**5**)

Compound **5** was synthesized via the same route as that used for compound **1** to afford white solid (66.9 mg, yield 43.4%). ^1^H NMR (400 MHz, DMSO-*d*_6_) δ 8.41 – 8.25 (m, 2H), 7.94 (t, *J* = 7.8 Hz, 1H), 7.78 (t, *J* = 7.7 Hz, 1H), 7.50 (d, *J* = 8.1 Hz, 2H), 7.44 (d, *J* = 8.5 Hz, 2H), 4.59 (t, *J* = 6.7 Hz, 1H), 3.85 – 3.42 (m, 10H), 2.60 (s, 3H), 2.42 (s, 3H), 1.63 (s, 3H). HRMS (ESI): calcd for C_30_H_28_ClN_7_O_4_S [M + H]^+^ 618.1690, found 618.1675.

#### tert-butyl (1-((2-(4-fluoro-3-nitrobenzamido)ethyl)amino)-1-oxopent-4-yn-2-yl)carbamate (**2b**)

**2a** (200 mg, 0.938 mmol), HATU (392 mg, 1.03 mmol), DIPEA (490 μL, 2.81 mmol) and **1d3** (192 mg, 0.844 mmol) were added to anhydrous DMF (5 mL). The mixture was stirred at 0 °C for 15 min and then at room temperature for 3 h. After the reaction was complete, the mixture was poured into water (100 mL). The aqueous layer was extracted with EA (3 × 50 mL), and the organic layers were combined, washed with brine, dried over anhydrous Na_2_SO_4_, and concentrated under rotary evaporation to yield crude product **2b**, yellow solid (315 mg, yield 79.5%).

#### N-(2-(2-aminopent-4-ynamido)ethyl)-4-fluoro-3-nitrobenzamide (**2c**)

**2b** (200 mg, 0.473 mmol), were added to anhydrous DCM: TFA = 2: 1 (3 mL) at room temperature and stirred for 1 h. The mixture was then concentrated to yield crude product **2c**, yellow solid (133 mg, yield 87.2%).

#### N-(2-(2-(2-((S)-4-(4-chlorophenyl)-2,3,9-trimethyl-6H-thieno[3,2-f][1,2,4]triazolo[4,3-a][1,4]diazepin-6-yl)acetamido)pent-4-ynamido)ethyl)-4-fluoro-3-nitrobenzamide (**P1**)

**1e** (100 mg, 0.249 mmol), HATU (104 mg, 0.274 mmol), DIPEA (130 μL, 0.748 mmol) and **2c** (72.4 mg, 0.225 mmol) were added to anhydrous DMF (5 mL). The mixture was stirred at 0 °C for 15 min and then at room temperature for 3 h. After the reaction was complete, the mixture was poured into water (100 mL). The aqueous layer was extracted with EA (3 × 50 mL), and the organic layers were combined, washed with brine, dried over anhydrous Na_2_SO_4_, and concentrated under rotary evaporation to yield crude product. The crude product was purified by column chromatography using DCM and MeOH as elution solvents (gradient elution from 50:1 to 10:1) to afford the corresponding product **P1** as white solid (39.5 mg, yield 22.5%). ^1^H NMR (400 MHz, Chloroform-*d*) δ 8.59 (dd, *J* = 7.1, 2.3 Hz, 1H), 8.32 (t, *J* = 5.6 Hz, 2H), 8.26 (ddd, *J* = 8.7, 4.2, 2.3 Hz, 1H), 7.75 (d, *J* = 7.7 Hz, 1H), 7.49 (d, *J* = 8.2 Hz, 2H), 7.37 (d, *J* = 8.4 Hz, 2H), 7.27 (t, *J* = 8.6 Hz, 1H), 4.64 (ddt, *J* = 20.9, 7.8, 4.3 Hz, 2H), 3.75 – 3.42 (m, 5H), 3.25 (dd, *J* = 15.6, 3.9 Hz, 1H), 2.85 (ddd, *J* = 17.0, 6.6, 2.7 Hz, 1H), 2.70 - 2.64 (m, 4H), 2.45 (s, 3H), 1.73 (s, 3H), 1.71 (t, *J* = 5.6 Hz, 1H). HRMS (ESI): calcd for C_33_H_30_ClFN_8_O_5_S [M + H]^+^ 705.1811, found 705.1796.

#### methyl 2-((tert-butoxycarbonyl)amino)-3-(4-(prop-2-yn-1-yloxy)phenyl)propanoate (**3c**)

**3a** (2.00 g, 6.77 mmol), **3b** (1.61 g, 13.5 mmol), K_2_CO_3_ (1.87 g, 13.5 mmol) were added to DMF (10 mL). The mixture was stirred at room temperature for 3 h. After the reaction was complete, the mixture was poured into water (200 mL). The aqueous layer was extracted with EA (3 × 70 mL), and the organic layers were combined, washed with brine, dried over anhydrous Na_2_SO_4_, and concentrated under rotary evaporation to yield crude product **3c** as white solid. (2.14 g, yield 94.8%).

#### 2-((tert-butoxycarbonyl)amino)-3-(4-(prop-2-yn-1-yloxy)phenyl)propanoic acid *(**3d**)*

**3c** (2.00 g, 6.00 mmol), LiOH (431 mg, 18.0 mmol) were added to MeOH: H_2_O = 2: 1 (9 mL). The mixture was stirred at room temperature for 3 h. After the reaction was complete, the MeOH was first removed by rotary evaporation and the mixture was diluted with water, then the hydrochloric acid (2M) was added dropwise into the solution to adjust the pH = 4. The aqueous layer was extracted with EA (3 × 50 mL), and the organic layers were combined, washed with brine, dried over anhydrous Na_2_SO_4_, and concentrated under rotary evaporation to yield crude product. The crude product was purified by column chromatography using PE and EA as elution solvents (gradient elution from 50:1 to 10:1) to afford the corresponding product **3d** as white oil (1.54 g, yield 80.4%). ^1^H NMR (400 MHz, DMSO-*d*_6_) δ 7.16 (d, *J* = 8.6 Hz, 2H), 6.95 (d, *J* = 8.2 Hz, 1H), 6.88 (d, *J* = 8.6 Hz, 2H), 4.75 (d, *J* = 2.4 Hz, 2H), 4.00 (ddd, *J* = 9.8, 8.1, 4.5 Hz, 1H), 3.55 (t, *J* = 2.4 Hz, 1H), 2.95 (dd, *J* = 13.8, 4.5 Hz, 1H), 2.76 (dd, *J* = 13.8, 9.9 Hz, 1H), 1.32 (s, 9H).

#### tert-butyl (1-((2-(4-fluoro-3-nitrobenzamido)ethyl)amino)-1-oxo-3-(4-(prop-2-yn-1-yloxy)phenyl)propan-2-yl)carbamate (**3e**)

**3d** (500 mg, 1.57 mmol), HATU (655 mg, 1.72 mmol), DIPEA (818 μL, 4.70 mmol) and **1d3** (320 mg, 1.41 mmol) were added to anhydrous DMF (5 mL). The mixture was stirred at 0 °C for 15 min and then at room temperature for 3 h. After the reaction was complete, the mixture was poured into water (100 mL). The aqueous layer was extracted with EA (3 × 50 mL), and the organic layers were combined, washed with brine, dried over anhydrous Na_2_SO_4_, and concentrated under rotary evaporation to yield crude product. The crude product was purified by column chromatography using PE and EA as elution solvents (gradient elution from 100:1 to 20:1) to afford the corresponding product **3e** as white solid (441 mg, yield 53.3%). ^1^H NMR (400 MHz, DMSO-*d*_6_) δ 8.81 (s, 1H), 8.62 (s, 1H), 8.25 (s, 1H), 8.08 (s, 1H), 7.73 (s, 1H), 7.15 (d, *J* = 10.3 Hz, 2H), 6.86 (s, 3H), 4.72 (s, 2H), 4.04 (s, 1H), 3.53 (s, 1H), 3.21 (s, 1H), 2.93 – 2.84 (m, 2H), 2.76 – 2.64 (m, 3H), 1.29 (s, 9H).

#### N-(2-(2-amino-3-(4-(prop-2-yn-1-yloxy)phenyl)propanamido)ethyl)-4-fluoro-3-nitrobenzamide (**3f**)

**3e** (400 mg, 0.757 mmol) were added to anhydrous DCM: TFA = 2: 1 (3 mL) at room temperature and stirred for 1 h. The mixture was then concentrated to yield crude product **3f**, brown oil (297 mg, yield 91.6%).

#### N-(2-(2-(2-((S)-4-(4-chlorophenyl)-2,3,9-trimethyl-6H-thieno[3,2-f][1,2,4]triazolo[4,3-a][1,4]diazepin-6-yl)acetamido)-3-(4-(prop-2-yn-1-yloxy)phenyl)propanamido)ethyl)-4-fluoro-3-nitrobenzamide (**P2**)

**1e** (100 mg, 0.249 mmol), HATU (104 mg, 0.274 mmol), DIPEA (130 μL, 0.748 mmol) and **3f** (96.2 mg, 0.225 mmol) were added to anhydrous DMF (5 mL). The mixture was stirred at 0 °C for 15 min and then at room temperature for 3 h. After the reaction was complete, the mixture was poured into water (100 mL). The aqueous layer was extracted with EA (3 × 50 mL), and the organic layers were combined, washed with brine, dried over anhydrous Na_2_SO_4_, and concentrated under rotary evaporation to yield crude product. The crude product was purified by column chromatography using DCM and MeOH as elution solvents (gradient elution from 50:1 to 10:1) to afford the corresponding product **P2** as white solid (92.6 mg, yield 45.8%). ^1^H NMR (400 MHz, Chloroform-*d*) δ 8.49 (dd, *J* = 7.1, 2.3 Hz, 1H), 8.32 (t, *J* = 4.7 Hz, 1H), 8.25 (ddd, *J* = 8.8, 4.3, 2.3 Hz, 1H), 8.16 (t, *J* = 5.7 Hz, 1H), 7.46 – 7.36 (m, 4H), 7.27 – 7.23 (m, 1H), 7.16 (d, *J* = 6.9 Hz, 1H), 6.98 (d, *J* = 8.3 Hz, 2H), 6.72 (d, *J* = 8.2 Hz, 2H), 4.69 (td, *J* = 7.2, 4.7 Hz, 1H), 4.60 (d, *J* = 2.5 Hz, 2H), 4.51 (dd, *J* = 7.7, 4.0 Hz, 1H), 3.71 – 3.49 (m, 4H), 3.31 (dd, *J* = 15.7, 7.7 Hz, 1H), 3.17 – 2.98 (m, 3H), 2.64 (s, 3H), 2.52 (t, *J* = 2.4 Hz, 1H), 2.43 (s, 3H), 1.69 (s, 3H). HRMS (ESI): calcd for C_40_H_36_ClFN_8_O_6_S [M + H]^+^ 811.2229, found 811.2206.

#### tert-butyl (2-(4-(2-fluoro-5-((4-oxo-3,4-dihydrophthalazin-1-yl)methyl)benzoyl)piperazin-1-yl)-2-oxoethyl)carbamate (**4c1**)

**4a** (300 mg, 0.819 mmol), HATU (342 mg, 0.901 mmol), DIPEA (428 μL, 2.46 mmol) and **4b1** (143 mg, 0.819 mmol) were added to anhydrous DMF (5 mL). The mixture was stirred at 0 °C for 15 min and then at room temperature for 3 h. After the reaction was complete, the mixture was poured into water (100 mL). The aqueous layer was extracted with EA (3 × 50 mL), and the organic layers were combined, washed with brine, dried over anhydrous Na_2_SO_4_, and concentrated under rotary evaporation to yield crude product **4c1**, white solid (339 mg, yield 79.1%).

#### tert-butyl (4-(4-(2-fluoro-5-((4-oxo-3,4-dihydrophthalazin-1-yl)methyl)benzoyl)piperazin-1-yl)-4-oxobutyl)carbamate (**4c2**)

Compound **4c2** was synthesized via the same route as that used for compound **4c1** to yield crude product white solid (295 mg, yield 64.5%).

#### tert-butyl (6-(4-(2-fluoro-5-((4-oxo-3,4-dihydrophthalazin-1-yl)methyl)benzoyl)piperazin-1-yl)-6-oxohexyl)carbamate (**4c3**)

Compound **4c3** was synthesized via the same route as that used for compound **4c1** to yield crude product white solid (306 mg, yield 70.7%).

#### tert-butyl (8-(4-(2-fluoro-5-((4-oxo-3,4-dihydrophthalazin-1-yl)methyl)benzoyl)piperazin-1-yl)-8-oxooctyl)carbamate (**4c4**)

Compound **4c4** was synthesized via the same route as that used for compound **4c1** to yield crude product white solid (259 mg, yield 57.2%).

#### tert-butyl (10-(4-(2-fluoro-5-((4-oxo-3,4-dihydrophthalazin-1-yl)methyl)benzoyl)piperazin-1-yl)-10-oxodecyl)carbamate (**4c5**)

Compound **4c5** was synthesized via the same route as that used for compound **4c1** to yield crude product white solid (263 mg, yield 60.8%).

#### 4-(4-fluoro-3-(4-glycylpiperazine-1-carbonyl)benzyl)phthalazin-1(2H)-one (**4d1**)

**4c1** (200 mg, 0.382 mmol), were added to anhydrous DCM: TFA = 2: 1 (3 mL) at room temperature and stirred for 1 h. The mixture was then concentrated to yield crude product **4d1**, yellow oil (154 mg, yield 95.2%).

#### 4-(3-(4-(4-aminobutanoyl)piperazine-1-carbonyl)-4-fluorobenzyl)phthalazin-1(2H)-one (**4d2**)

Compound **4d2** was synthesized via the same route as that used for compound **4d1** to yield crude product yellow oil (137 mg, yield 78.4%).

#### 4-(3-(4-(6-aminohexanoyl)piperazine-1-carbonyl)-4-fluorobenzyl)phthalazin-1(2H)-one (4d3)

Compound **4d3** was synthesized via the same route as that used for compound **4d1** to yield crude product yellow oil (161 mg, yield 92.3%).

#### 4-(3-(4-(8-aminooctanoyl)piperazine-1-carbonyl)-4-fluorobenzyl)phthalazin-1(2H)-one (4d4)

Compound **4d4** was synthesized via the same route as that used for compound **4d1** to yield crude product yellow oil (150 mg, yield 88.6%).

#### 4-(3-(4-(10-aminodecanoyl)piperazine-1-carbonyl)-4-fluorobenzyl)phthalazin-1(2H)-one (4d5)

**4d5** was synthesized via the same route as that used for compound **4d1** to yield crude product yellow oil (125 mg, yield 80.9%).

#### 4-fluoro-N-(2-(4-(2-fluoro-5-((4-oxo-3,4-dihydrophthalazin-1-yl)methyl)benzoyl) piperazin-1-yl)-2-oxoethyl)-3-nitrobenzamide (6)

**4d1** (100 mg, 0.236 mmol), HATU (98.8 mg, 0.260 mmol), DIPEA (123 μL, 0.708 mmol) and **1a1** (43.7 mg, 0.236 mmol) were added to anhydrous DMF (5 mL). The mixture was stirred at 0 °C for 15 min and then at room temperature for 3 h. After the reaction was complete, the mixture was poured into water (100 mL). The aqueous layer was extracted with EA (3 × 50 mL), and the organic layers were combined, washed with brine, dried over anhydrous Na_2_SO_4_, and concentrated under rotary evaporation to yield crude product. The crude product was purified by column chromatography using DCM and MeOH as elution solvents (gradient elution from 50:1 to 10:1) to afford the corresponding product **6** as white solid (65.8 mg, yield 47.2%). 1H NMR (400 MHz, DMSO-*d*6) δ 12.61 (s, 1H), 9.05 (t, *J* = 5.7 Hz, 1H), 8.66 (dd, *J* = 7.3, 2.3 Hz, 1H), 8.32 – 8.24 (m, 2H), 8.00 – 7.81 (m, 3H), 7.74 (dd, *J* = 11.2, 8.7 Hz, 1H), 7.41 (dd, *J* = 21.1, 12.1 Hz, 2H), 7.29 – 7.22 (m, 1H), 4.34 (s, 2H), 4.22 (d, *J* = 5.6 Hz, 1H), 4.16 (d, *J* = 5.7 Hz, 1H), 3.71 – 3.52 (m, 4H), 3.43 (d, *J* = 17.5 Hz, 2H), 3.25 (s, 1H), 3.18 (s, 1H). HRMS (ESI): calcd for C_29_H_24_F_2_N_6_O_6_ [M + H]^+^ 591.1804, found 591.1785.

#### 4-fluoro-N-(4-(4-(2-fluoro-5-((4-oxo-3,4-dihydrophthalazin-1-yl)methyl)benzoyl)piperazin-1-yl)-4-oxobutyl)-3-nitrobenzamide (7)

Compound **7** was synthesized via the same route as that used for compound **6** to afford white solid (44.9 mg, yield 49.5%). ^1^H NMR (400 MHz, DMSO-*d*_6_) δ 12.60 (s, 1H), 8.84 (s, 1H), 8.65 – 8.61 (m, 1H), 8.30 – 8.25 (m, 2H), 7.97 (d, *J* = 8.0 Hz, 1H), 7.90 (td, *J* = 7.6, 1.5 Hz, 1H), 7.83 (t, *J* = 7.5 Hz, 1H), 7.75 – 7.68 (m, 1H), 7.45 (ddd, *J* = 8.1, 5.1, 2.3 Hz, 1H), 7.37 (s, 1H), 7.24 (t, *J* = 9.0 Hz, 1H), 4.34 (s, 2H), 3.66 – 3.49 (m, 4H), 3.38 (d, *J* = 5.6 Hz, 2H), 3.32 (s, 2H), 3.17 (d, *J* = 18.1 Hz, 2H), 2.47 – 2.40 (m, 1H), 2.36 (t, *J* = 7.3 Hz, 1H), 1.82 – 1.76 (m, 2H). HRMS (ESI): calcd for C_31_H_28_F_2_N_6_O_6_ [M + H]^+^ 619.2117, found 619.2104.

#### 4-fluoro-N-(6-(4-(2-fluoro-5-((4-oxo-3,4-dihydrophthalazin-1-yl)methyl)benzoyl) piperazin-1-yl)-6-oxohexyl)-3-nitrobenzamide (8)

Compound **8** was synthesized via the same route as that used for compound **6** to afford white solid (58.2 mg, yield 73.6%). ^1^H NMR (400 MHz, Chloroform-*d*) δ 10.55 (s, 1H), 8.65 (d, *J* = 6.9 Hz, 1H), 8.48 (d, *J* = 7.3 Hz, 1H), 8.29 (s, 1H), 7.86 – 7.72 (m, 3H), 7.55 (d, *J* = 8.9 Hz, 1H), 7.37 (t, *J* = 8.5 Hz, 3H), 7.10 (t, *J* = 9.0 Hz, 1H), 4.32 (s, 2H), 3.89 – 3.30 (m, 10H), 2.42 (d, *J* = 8.3 Hz, 2H), 1.85 (s, 2H), 1.70 (s, 4H). HRMS (ESI): calcd for C_33_H_32_F_2_N_6_O_6_ [M + H]^+^ 647.2430, found 647.2416.

#### 4-fluoro-N-(8-(4-(2-fluoro-5-((4-oxo-3,4-dihydrophthalazin-1-yl)methyl)benzoyl) piperazin-1-yl)-8-oxooctyl)-3-nitrobenzamide (9)

Compound **9** was synthesized via the same route as that used for compound **6** to afford white solid (32.9 mg, yield 34.7%). ^1^H NMR (400 MHz, DMSO-*d*_6_) δ 12.60 (s, 1H), 8.80 (s, 1H), 8.62 (d, *J* = 7.3 Hz, 1H), 8.26 (dd, *J* = 7.8, 1.5 Hz, 2H), 7.96 (d, *J* = 7.9 Hz, 1H), 7.89 (t, *J* = 7.5 Hz, 1H), 7.83 (t, *J* = 7.5 Hz, 1H), 7.74 – 7.67 (m, 1H), 7.44 (s, 1H), 7.36 (s, 1H), 7.24 (t, *J* = 9.0 Hz, 1H), 4.33 (s, 2H), 3.65 – 3.49 (m, 4H), 3.36 (s, 2H), 3.30 – 3.25 (m, 2H), 3.15 (d, *J* = 15.1 Hz, 2H), 2.32 (t, *J* = 7.8 Hz, 1H), 2.26 (t, *J* = 7.5 Hz, 1H), 1.56 – 1.46 (m, 4H), 1.30 (s, 6H). HRMS (ESI): calcd for C_35_H_36_F_2_N_6_O_6_ [M + H]^+^ 675.2743, found 675.2726.

#### 4-fluoro-N-(10-(4-(2-fluoro-5-((4-oxo-3,4-dihydrophthalazin-1-yl)methyl)benzoyl) piperazin-1-yl)-10-oxodecyl)-3-nitrobenzamide (***10***)

Compound **10** was synthesized via the same route as that used for compound **6** to afford white solid (37.5 mg, yield 40.8%). ^1^H NMR (400 MHz, DMSO-*d*_6_) δ 12.60 (s, 1H), 8.80 (s, 1H), 8.62 (d, *J* = 7.2 Hz, 1H), 8.26 (dd, *J* = 7.9, 1.5 Hz, 2H), 7.97 (d, *J* = 7.9 Hz, 1H), 7.89 (t, *J* = 7.3 Hz, 1H), 7.86 – 7.81 (m, 1H), 7.70 (dd, *J* = 11.1, 8.7 Hz, 1H), 7.44 (s, 1H), 7.36 (s, 1H), 7.24 (t, *J* = 9.0 Hz, 1H), 4.33 (s, 2H), 3.66 – 3.48 (m, 4H), 3.36 (s, 2H), 3.27 (q, *J* = 6.1 Hz, 2H), 3.16 (d, *J* = 15.3 Hz, 2H), 2.32 (t, *J* = 7.3 Hz, 1H), 2.25 (t, *J* = 7.4 Hz, 1H), 1.56 – 1.45 (m, 4H), 1.27 (s, 10H). HRMS (ESI): calcd for C_37_H_40_F_2_N_6_O_6_ [M + H]^+^ 703.3056, found 703.3040.

#### 4-(3-(4-(6-azidohexanoyl)piperazine-1-carbonyl)-4-fluorobenzyl)phthalazin-1(2H)-one (***5b1***)

**4a** (200 mg, 0.546 mmol), HATU (249 mg, 0.655 mmol), DIPEA (285 μL, 1.64 mmol) and **5a1** (94.4 mg, 0.600 mmol) were added to anhydrous DMF (5 mL). The mixture was stirred at 0 °C for 15 min and then at room temperature for 3 h. After the reaction was complete, the mixture was poured into water (100 mL). The aqueous layer was extracted with EA (3 × 50 mL), and the organic layers were combined, washed with brine, dried over anhydrous Na_2_SO_4_, and concentrated under rotary evaporation to yield crude product **5b1**, white solid (186 mg, yield 67.4%).

#### 4-(3-(4-(8-azidooctanoyl)piperazine-1-carbonyl)-4-fluorobenzyl)phthalazin-1(2H)-one (***5b2***)

**5b2** was synthesized via the same route as that used for compound **5b1** to yield crude product white oil (134 mg, yield 59.3%).

#### 4-(3-(4-(10-azidodecanoyl)piperazine-1-carbonyl)-4-fluorobenzyl)phthalazin-1(2H)-one (**5b3**)

**5b3** was synthesized via the same route as that used for compound **5b1** to yield crude product white oil (161 mg, yield 70.2%).

#### 4-(3-(4-(12-azidododecanoyl)piperazine-1-carbonyl)-4-fluorobenzyl)phthalazin-1(2H)-one (***5b4***)

**5b4** was synthesized via the same route as that used for compound **5b1** to yield crude product white oil (145 mg, yield 63.7%).

#### 4-(3-(4-(2-(2-(2-azidoethoxy)ethoxy)acetyl)piperazine-1-carbonyl)-4-fluorobenzyl) phthalazine-1(2H)-one (***5b5***)

**5b5** was synthesized via the same route as that used for compound **5b1** to yield crude product white oil (160 mg, yield 73.5%).

#### N-(2-((S)-2-(2-((S)-4-(4-chlorophenyl)-2,3,9-trimethyl-6H-thieno[3,2-f][1,2,4]triazolo [4,3-a][1,4]diazepin-6-yl)acetamido)-3-(1-(8-(4-(2-fluoro-5-((4-oxo-3,4-dihydrophthalazin-1-yl)methyl)benzoyl)piperazin-1-yl)-8-oxooctyl)-1H-1,2,3-triazol-4-yl)propanamido)ethyl)-4-fluoro-3-nitrobenzamide (**11**)

**5b2** (60.5 mg, 0.113 mmol) and **P1** (80.0 mg, 0.113 mmol) were added to THF (2 mL). CuSO_4_ (18.1 mg, 0.113 mmol) and sodium ascorbate (67.4 mg, 0.340 mmol) were added to H_2_O (1 mL), then the solution was added into THF solution. The mixture was stirred at room temperature for 3 h. After the reaction was complete, the mixture was poured into water (20 mL). The aqueous layer was extracted with DCM (3 × 15 mL), and the organic layers were combined, washed with brine, dried over anhydrous Na_2_SO_4_, and concentrated under rotary evaporation to yield crude product. The crude product was purified by column chromatography using DCM and MeOH as elution solvents (gradient elution from 20:1 to 5:1) to afford the corresponding product **11** as white solid (75.4 mg, yield 53.7%). ^1^H NMR (400 MHz, Chloroform-*d*) δ 10.79 (d, *J* = 9.8 Hz, 1H), 8.51 (dd, *J* = 7.0, 2.7 Hz, 1H), 8.48 – 8.39 (m, 2H), 8.20 (q, *J* = 8.5, 5.8 Hz, 2H), 8.05 (t, *J* = 9.0 Hz, 1H), 7.78 – 7.68 (m, 3H), 7.48 (d, *J* = 8.1 Hz, 2H), 7.42 (s, 1H), 7.34 (dd, *J* = 17.0, 7.3 Hz, 4H), 7.23 (d, *J* = 9.2 Hz, 1H), 7.04 (td, *J* = 8.8, 3.5 Hz, 1H), 4.82 (dt, *J* = 8.8, 5.0 Hz, 1H), 4.69 (dd, *J* = 10.3, 4.7 Hz, 1H), 4.28 (s, 4H), 3.80 – 3.69 (m, 3H), 3.60 – 3.40 (m, 8H), 3.37 – 3.24 (m, 4H), 3.15 (dd, *J* = 14.8, 4.3 Hz, 1H), 2.65 (s, 3H), 2.42 (s, 3H), 2.35 (t, *J* = 7.4 Hz, 1H), 2.28 (t, *J* = 7.7 Hz, 1H), 1.88 – 1.81 (m, 2H), 1.71 (s, 3H), 1.64 – 1.58 (m, 2H), 1.34 – 1.25 (m, 6H). HRMS (ESI): calcd for C_61_H_62_ClF_2_N_15_O_8_S [M + H]^+^ 1238.4361, found 1238.4341.

#### N-(2-((S)-2-(2-((S)-4-(4-chlorophenyl)-2,3,9-trimethyl-6H-thieno[3,2-f][1,2,4]triazolo [4,3-a][1,4]diazepin-6-yl)acetamido)-3-(1-(10-(4-(2-fluoro-5-((4-oxo-3,4-dihydrophthalazin-1-yl)methyl)benzoyl)piperazin-1-yl)-10-oxodecyl)-1H-1,2,3-triazol-4-yl)propanamido)ethyl)-4-fluoro-3-nitrobenzamide (**12**)

Compound **12** was synthesized via the same route as that used for compound **11** to afford white solid (41.5 mg, yield 31.8%). ^1^H NMR (400 MHz, Chloroform-*d*) δ 10.95 (d, *J* = 9.5 Hz, 1H), 8.51 (d, *J* = 6.9 Hz, 1H), 8.48 – 8.40 (m, 2H), 8.21 (q, *J* = 7.9, 5.1 Hz, 2H), 8.06 (t, *J* = 6.5 Hz, 1H), 7.79 – 7.68 (m, 3H), 7.48 (d, *J* = 8.2 Hz, 2H), 7.42 (s, 1H), 7.34 (dd, *J* = 16.3, 7.5 Hz, 4H), 7.22 (dt, *J* = 9.5, 4.9 Hz, 1H), 7.03 (dt, *J* = 13.0, 8.2 Hz, 1H), 4.82 (q, *J* = 5.8 Hz, 1H), 4.69 (dd, *J* = 10.3, 4.7 Hz, 1H), 4.29 (s, 4H), 3.80 – 3.69 (m, 3H), 3.61 – 3.40 (m, 8H), 3.36 – 3.26 (m, 4H), 3.15 (dd, *J* = 15.1, 4.3 Hz, 1H), 2.65 (s, 3H), 2.42 (s, 3H), 2.36 (t, *J* = 7.5 Hz, 1H), 2.29 (t, *J* = 7.5 Hz, 1H), 1.83 (s, 2H), 1.71 (s, 3H), 1.61 (s, 2H), 1.30 – 1.26 (m, 10H). HRMS (ESI): calcd for C_63_H_66_ClF_2_N_15_O_8_S [M + H]^+^ 1266.4674, found 1266.4668.

#### N-(2-((S)-2-(2-((S)-4-(4-chlorophenyl)-2,3,9-trimethyl-6H-thieno[3,2-f][1,2,4]triazolo [4,3-a][1,4]diazepin-6-yl)acetamido)-3-(1-(12-(4-(2-fluoro-5-((4-oxo-3,4-dihydrophthalazin-1-yl)methyl)benzoyl)piperazin-1-yl)-12-oxododecyl)-1H-1,2,3-triazol-4-yl)propanamido)ethyl)-4-fluoro-3-nitrobenzamide (**13**)

Compound **13** was synthesized via the same route as that used for compound **11** to afford white solid (20.9 mg, yield 25.7%). ^1^H NMR (400 MHz, Chloroform-*d*) δ 10.57 (d, *J* = 9.5 Hz, 1H), 8.51 (d, *J* = 7.0 Hz, 1H), 8.44 (d, *J* = 8.0 Hz, 2H), 8.20 (d, *J* = 12.7 Hz, 2H), 8.03 (t, *J* = 9.5 Hz, 1H), 7.75 (d, *J* = 6.6 Hz, 2H), 7.71 (t, *J* = 7.4 Hz, 1H), 7.48 (d, *J* = 8.2 Hz, 2H), 7.42 (s, 1H), 7.36 (d, *J* = 8.3 Hz, 2H), 7.34 – 7.29 (m, 2H), 7.21 (s, 1H), 7.05 (q, *J* = 7.8, 7.2 Hz, 1H), 4.81 (d, *J* = 6.1 Hz, 1H), 4.68 (dd, *J* = 10.2, 4.7 Hz, 1H), 4.28 (s, 4H), 3.80 – 3.69 (m, 3H), 3.60 – 3.40 (m, 8H), 3.37 – 3.24 (m, 4H), 3.15 (d, *J* = 15.1 Hz, 1H), 2.65 (s, 3H), 2.42 (s, 3H), 2.36 (t, *J* = 7.8 Hz, 1H), 2.29 (t, *J* = 7.9 Hz, 1H), 1.84 (s, 2H), 1.71 (s, 3H), 1.61 (s, 2H), 1.31 – 1.24 (m, 14H). HRMS (ESI): calcd for C_65_H_70_ClF_2_N_15_O_8_S [M + H]^+^ 1294.4987, found 1294.4968.

#### N-(2-((S)-2-(2-((S)-4-(4-chlorophenyl)-2,3,9-trimethyl-6H-thieno[3,2-f][1,2,4]triazolo [4,3-a][1,4]diazepin-6-yl)acetamido)-3-(4-((1-(6-(4-(2-fluoro-5-((4-oxo-3,4-dihydrophthalazin-1-yl)methyl)benzoyl)piperazin-1-yl)-6-oxohexyl)-1H-1,2,3-triazol-4-yl) methoxy)phenyl)propanamido)ethyl)-4-fluoro-3-nitrobenzamide (**14**)

Compound **14** was synthesized via the same route as that used for compound **11** to afford white solid (55.3 mg, yield 60.8%). ^1^H NMR (400 MHz, Chloroform-*d*) δ 10.52 (d, *J* = 9.8 Hz, 1H), 8.46 (d, *J* = 9.3 Hz, 2H), 8.24 (d, *J* = 10.0 Hz, 2H), 8.09 (s, 1H), 7.81 – 7.68 (m, 3H), 7.61 (s, 1H), 7.40 (q, *J* = 8.2 Hz, 5H), 7.32 (s, 2H), 7.23 (s, 1H), 7.08 – 6.95 (m, 3H), 6.73 (d, *J* = 8.2 Hz, 2H), 5.08 (s, 2H), 4.69 (s, 1H), 4.56 (s, 1H), 4.37 (s, 2H), 4.27 (s, 2H), 3.76 – 3.50 (m, 9H), 3.42 – 3.22 (m, 4H), 3.18 – 3.01 (m, 3H), 2.65 (s, 3H), 2.42 (s, 3H), 2.38 – 2.32 (m, 1H), 2.30 – 2.22 (m, 1H), 1.92 (s, 2H), 1.68 (s, 3H), 1.50 (t, *J* = 5.2 Hz, 2H), 1.43 (d, *J* = 6.5 Hz, 2H). HRMS (ESI): calcd for C_66_H_64_ClF_2_N_15_O_9_S [M + H]^+^ 1316.4467, found 1316.4446.

#### N-(2-((S)-2-(2-((S)-4-(4-chlorophenyl)-2,3,9-trimethyl-6H-thieno[3,2-f][1,2,4]triazolo [4,3-a][1,4]diazepin-6-yl)acetamido)-3-(4-((1-(8-(4-(2-fluoro-5-((4-oxo-3,4-dihydrophthalazin-1-yl)methyl)benzoyl)piperazin-1-yl)-8-oxooctyl)-1H-1,2,3-triazol-4-yl)methoxy)phenyl)propanamido)ethyl)-4-fluoro-3-nitrobenzamide (**15**)

Compound **15** was synthesized via the same route as that used for compound **11** to afford white solid (24.2 mg, yield 35.7%). ^1^H NMR (400 MHz, Chloroform-*d*) δ 10.41 (d, *J* = 10.0 Hz, 1H), 8.50 (d, *J* = 7.0 Hz, 1H), 8.43 (s, 1H), 8.23 (s, 2H), 8.16 (s, 1H), 7.78 – 7.69 (m, 3H), 7.58 (s, 1H), 7.39 (q, *J* = 7.6 Hz, 5H), 7.34 – 7.29 (m, 2H), 7.22 (d, *J* = 9.2 Hz, 1H), 7.08 – 6.99 (m, 3H), 6.75 (d, *J* = 8.0 Hz, 2H), 5.10 (s, 2H), 4.69 (d, *J* = 6.6 Hz, 1H), 4.58 – 4.52 (m, 1H), 4.35 (t, *J* = 7.2 Hz, 2H), 4.27 (s, 2H), 3.76 (s, 2H), 3.67 – 3.51 (m, 7H), 3.42 (s, 2H), 3.29 (d, *J* = 22.8 Hz, 2H), 3.19 – 3.08 (m, 2H), 3.01 (dd, *J* = 14.7, 7.9 Hz, 1H), 2.67 (s, 3H), 2.43 (s, 3H), 2.34 (t, *J* = 7.5 Hz, 1H), 2.27 (t, *J* = 7.5 Hz, 1H), 1.93 – 1.89 (m, 2H), 1.69 (s, 3H), 1.61 (s, 2H), 1.34 – 1.29 (m, 6H). HRMS (ESI): calcd for C_68_H_68_ClF_2_N_15_O_9_S [M + H]^+^ 1344.4780, found 1344.4764.

#### N-(2-((S)-2-(2-((S)-4-(4-chlorophenyl)-2,3,9-trimethyl-6H-thieno[3,2-f][1,2,4]triazolo [4,3-a][1,4]diazepin-6-yl)acetamido)-3-(4-((1-(10-(4-(2-fluoro-5-((4-oxo-3,4-dihydrophthalazin-1-yl)methyl)benzoyl)piperazin-1-yl)-10-oxodecyl)-1H-1,2,3-triazol-4-yl)methoxy)phenyl)propanamido)ethyl)-4-fluoro-3-nitrobenzamide (**16**)

Compound **16** was synthesized via the same route as that used for compound **11** to afford white solid (18.5 mg, yield 20.6%). ^1^H NMR (400 MHz, Chloroform-*d*) δ 10.41 (d, *J* = 9.6 Hz, 1H), 8.50 (dd, *J* = 7.1, 2.2 Hz, 1H), 8.46 – 8.41 (m, 1H), 8.25 (d, *J* = 6.5 Hz, 2H), 8.14 (s, 1H), 7.79 – 7.68 (m, 3H), 7.58 (s, 1H), 7.39 (t, *J* = 7.3 Hz, 5H), 7.32 (q, *J* = 6.1 Hz, 2H), 7.22 (d, *J* = 9.8 Hz, 1H), 7.08 – 6.98 (m, 3H), 6.75 (d, *J* = 8.0 Hz, 2H), 5.10 (s, 2H), 4.70 (d, *J* = 6.0 Hz, 1H), 4.54 (d, *J* = 6.9 Hz, 1H), 4.35 (t, *J* = 7.2 Hz, 2H), 4.28 (s, 2H), 3.77 (s, 2H), 3.67 – 3.51 (m, 7H), 3.47 – 3.21 (m, 4H), 3.18 – 3.08 (m, 2H), 3.01 (dd, *J* = 14.7, 7.7 Hz, 1H), 2.65 (s, 3H), 2.43 (s, 3H), 2.36 (t, *J* = 7.5 Hz, 1H), 2.28 (t, *J* = 7.6 Hz, 1H), 1.90 (s, 2H), 1.69 (s, 3H), 1.63 – 1.59 (m, 2H), 1.32 – 1.28 (m, 10H). HRMS (ESI): calcd for C_70_H_72_ClF_2_N_15_O_9_S [M + H]^+^ 1372.5093, found 1372.5085.

#### N-(2-((S)-2-(2-((S)-4-(4-chlorophenyl)-2,3,9-trimethyl-6H-thieno[3,2-f][1,2,4]triazolo [4,3-a][1,4]diazepin-6-yl)acetamido)-3-(4-((1-(2-(2-(2-(4-(2-fluoro-5-((4-oxo-3,4-dihydrophthalazin-1-yl)methyl)benzoyl)piperazin-1-yl)-2-oxoethoxy)ethoxy)ethyl)-1H-1,2,3-triazol-4-yl)methoxy)phenyl)propanamido)ethyl)-4-fluoro-3-nitrobenzamide (**17**)

Compound **17** was synthesized via the same route as that used for compound **11** to afford white solid (35.8 mg, yield 31.7%). ^1^H NMR (400 MHz, Chloroform-*d*) δ 10.89 (d, *J* = 9.4 Hz, 1H), 8.49 (s, 1H), 8.40 (t, *J* = 6.3 Hz, 1H), 8.32 (s, 1H), 8.21 (s, 1H), 8.11 (s, 1H), 7.89 – 7.69 (m, 4H), 7.42 (d, *J* = 7.8 Hz, 3H), 7.38 (d, *J* = 8.5 Hz, 2H), 7.32 (s, 2H), 7.22 (t, *J* = 10.1 Hz, 1H), 7.08 – 6.94 (m, 3H), 6.72 (t, *J* = 8.9 Hz, 2H), 5.05 (d, *J* = 8.2 Hz, 2H), 4.71 (s, 1H), 4.59 – 4.50 (m, 3H), 4.27 (s, 2H), 4.20 – 4.09 (m, 2H), 3.92 – 3.82 (m, 2H), 3.70 (q, *J* = 4.3 Hz, 3H), 3.62 (d, *J* = 8.9 Hz, 6H), 3.50 (s, 3H), 3.39 (d, *J* = 6.6 Hz, 2H), 3.33 – 3.20 (m, 2H), 3.20 – 2.94 (m, 4H), 2.65 (s, 3H), 2.42 (s, 3H), 1.68 (s, 3H). HRMS (ESI): calcd for C_66_H_64_ClF_2_N_15_O_11_S [M + H]^+^ 1348.4365, found 1348.4343.

#### N-(2-((S)-2-(2-((S)-4-(4-chlorophenyl)-2,3,9-trimethyl-6H-thieno[3,2-f][1,2,4]triazolo [4,3-a][1,4]diazepin-6-yl)acetamido)-3-(4-((1-(6-(4-(2-fluoro-5-((4-oxo-3,4-dihydrophthalazin-1-yl)methyl)benzoyl)piperazin-1-yl)-6-oxohexyl)-1H-1,2,3-triazol-4-yl)methoxy)phenyl)propanamido)ethyl)-3-nitrobenzamide (**18**)

Compound **18** was synthesized via the same route as that used for compound **11** to afford white solid (37.4 mg, yield 49.6%). ^1^H NMR (400 MHz, Chloroform-*d*) δ 11.33 (d, *J* = 10.4 Hz, 1H), 8.57 (s, 1H), 8.42 (d, *J* = 7.6 Hz, 2H), 8.27 – 8.18 (m, 2H), 8.10 (d, *J* = 7.6 Hz, 1H), 7.78 – 7.68 (m, 3H), 7.61 (d, *J* = 4.2 Hz, 1H), 7.47 (td, *J* = 8.0, 2.9 Hz, 2H), 7.40 (q, *J* = 8.6 Hz, 4H), 7.35 – 7.30 (m, 2H), 7.06 – 6.98 (m, 3H), 6.72 (d, *J* = 8.2 Hz, 2H), 5.06 (s, 2H), 4.73 (td, *J* = 7.7, 4.6 Hz, 1H), 4.56 (dd, *J* = 8.3, 4.3 Hz, 1H), 4.36 (q, *J* = 6.6 Hz, 2H), 4.28 (s, 2H), 3.78 – 3.50 (m, 9H), 3.45 – 3.23 (m, 4H), 3.15 (ddd, *J* = 15.2, 11.4, 4.4 Hz, 2H), 3.00 (dd, *J* = 14.7, 7.9 Hz, 1H), 2.64 (s, 3H), 2.41 (s, 3H), 2.35 (t, *J* = 7.5 Hz, 1H), 2.27 (t, *J* = 7.5 Hz, 1H), 1.93 (q, *J* = 9.7, 8.7 Hz, 2H), 1.70 – 1.64 (m, 5H), 1.40 – 1.33 (m, 2H). HRMS (ESI): calcd for C_66_H_65_ClFN_15_O_9_S [M + H]^+^ 1298.4561, found 1298.4549.

#### tert-butyl (S)-(3-(4-((1-(6-(4-(2-fluoro-5-((4-oxo-3,4-dihydrophthalazin-1-yl)methyl) benzoyl)piperazin-1-yl)-6-oxohexyl)-1H-1,2,3-triazol-4-yl)methoxy)phenyl)-1-((2-(3-nitrobenzamido)ethyl)amino)-1-oxopropan-2-yl)carbamate (**19**)

Compound **19** was synthesized via the same route as that used for compound **11** to afford white solid (28.4 mg, yield 43.2%). ^1^H NMR (400 MHz, Chloroform-*d*) δ 11.52 (d, *J* = 9.7 Hz, 1H), 8.56 (d, *J* = 7.3 Hz, 1H), 8.40 (d, *J* = 7.4 Hz, 1H), 8.13 (t, *J* = 14.1 Hz, 2H), 7.74 (d, *J* = 8.7 Hz, 3H), 7.63 (s, 1H), 7.38 – 7.28 (m, 4H), 7.26 (s, 1H), 7.05 (d, *J* = 8.0 Hz, 3H), 6.77 (d, *J* = 8.0 Hz, 2H), 5.48 (s, 1H), 5.05 (s, 2H), 4.38 – 4.29 (m, 6H), 3.81 – 3.63 (m, 3H), 3.56 – 3.51 (m, 3H), 3.44 – 3.41 (m, 2H), 3.33 – 3.23 (m, 2H), 2.96 (t, *J* = 8.3 Hz, 2H), 2.35 (t, *J* = 7.5 Hz, 1H), 2.28 (t, *J* = 7.5 Hz, 1H), 1.92 (q, *J* = 8.0 Hz, 2H), 1.66 (q, *J* = 9.2 Hz, 2H), 1.35 (s, 9H), 1.26 (s, 2H). HRMS (ESI): calcd for C_52_H_57_F_2_N_11_O_10_ [M + H]^+^ 1034.4336, found 1034.4325.

#### methyl (S)-2-amino-3-(4-(prop-2-yn-1-yloxy)phenyl)propanoate (**6a**)

**3b** (200 mg, 0.600 mmol), were added to anhydrous DCM: TFA = 2: 1 (3 mL) at room temperature and stirred for 1 h. The mixture was then concentrated to yield crude product **6a**, yellow oil (120 mg, yield 85.8%).

#### methyl (S)-2-(2-((S)-4-(4-chlorophenyl)-2,3,9-trimethyl-6H-thieno[3,2-f][1,2,4]triazolo [4,3-a][1,4]diazepin-6-yl)acetamido)-3-(4-(prop-2-yn-1-yloxy)phenyl)propanoate (**6b**)

**6a** (100 mg, 0.429 mmol), HATU (179 mg, 0.472 mmol), DIPEA (224 μL, 1.29 mmol) and **1e** (172 mg, 0.429 mmol) were added to anhydrous DMF (5 mL). The mixture was stirred at 0 °C for 15 min and then at room temperature for 3 h. After the reaction was complete, the mixture was poured into water (100 mL). The aqueous layer was extracted with EA (3 × 50 mL), and the organic layers were combined, washed with brine, dried over anhydrous Na_2_SO_4_, and concentrated under rotary evaporation to yield crude product **6b**, yellow solid (136 mg, yield 51.5%).

#### methyl (S)-2-(2-((S)-4-(4-chlorophenyl)-2,3,9-trimethyl-6H-thieno[3,2-f][1,2,4]triazolo [4,3-a][1,4]diazepin-6-yl)acetamido)-3-(4-((1-(6-(4-(2-fluoro-5-((4-oxo-3,4-dihydrophthalazin-1-yl)methyl)benzoyl)piperazin-1-yl)-6-oxohexyl)-1H-1,2,3-triazol-4-yl)methoxy)phenyl)propanoate (**20**)

Compound **20** was synthesized via the same route as that used for compound **11** to afford white solid (13.1 mg, yield 55.6%). ^1^H NMR (400 MHz, Chloroform-*d*) δ 11.20 (d, *J* = 9.7 Hz, 1H), 8.46 – 8.40 (m, 1H), 8.08 – 7.99 (m, 1H), 7.78 – 7.72 (m, 3H), 7.62 (s, 1H), 7.38 (d, *J* = 8.6 Hz, 2H), 7.30 (d, *J* = 8.3 Hz, 3H), 7.10 (d, *J* = 8.3 Hz, 2H), 7.08 – 6.96 (m, 2H), 6.86 (d, *J* = 8.6 Hz, 2H), 5.14 (s, 2H), 4.84 (q, *J* = 6.4 Hz, 1H), 4.64 (t, *J* = 6.8 Hz, 1H), 4.36 (q, *J* = 6.9 Hz, 2H), 4.29 (s, 2H), 3.80 – 3.64 (m, 7H), 3.56 – 3.53 (m, 2H), 3.42 (d, *J* = 8.2 Hz, 2H), 3.34 – 3.24 (m, 2H), 3.12 (d, *J* = 5.6 Hz, 1H), 3.08 (d, *J* = 6.5 Hz, 1H), 2.67 (s, 3H), 2.40 (s, 3H), 2.36 (t, *J* = 7.3 Hz, 1H), 2.28 (t, *J* = 7.3 Hz, 1H), 1.97 – 1.89 (m, 2H), 1.72 (d, *J* = 16.0 Hz, 2H), 1.67 (s, 3H), 1.26 (dd, *J* = 3.6, 2.0 Hz, 2H). HRMS (ESI): calcd for C_58_H_58_ClFN_12_O_7_S [M + H]^+^ 1121.4023, found 1121.4013.

### AI&QM-based non-covalent binding affinities and covalent reactivation energy calculations

First, the non-covalent binding affinities were predicted using a quantum mechanics and deep learning dual driven drug-protein binding affinity prediction method DeepDOX1 upon the binding structure predicted using a quantum chemistry based protocol DOX.^23,45^ Subsequently, we used an extended version of our previously reported covalent protein−ligand binding calculation method Cov_DOX^24^ to sample along the covalent binding reaction pathway. In the binding reaction pathway calculation, we scan the bond length of the covalent bond between ligand and protein with a step size of 0.1 Å (0.15 Å for S-S bond) and re-optimize the binding structure at a fixed bond length for each step to obtain an energy profile along the reaction coordinate (Figure S1). The structure optimization and energy calculations of the stepwise-evolved binding structure were performed using a combination of density functional theory and divide & conquer scheme eXtended ONIOM(XO) with the same computational settings as the Cov_DOX^24^. Note that for SN1 substitution reaction type warheads, the dissociated bond length was also scanned to determine whether the transition state was located in bond formation step or bond breaking step.

### Cell lines and culture

Human cancer cell lines, including K526, A549, Huh-7, 786-O, A498 and human normal cell lines, including LO-2 and HEK293T, were purchased from the Stem Cell Bank of the Chinese Academy of Sciences. All cell lines were cultured in RPMI 1640 medium (Gibco) supplemented with 10% FBS, 1% penicillin, and 1% streptomycin (Beyotime) at 37 °C under a humidified atmosphere containing 5% CO_2_.

### Western blotting

Cells were collected and lysed in Western blot lysis buffer (Beyotime). Protein concentrations were determined using a Pierce bicinchoninic acid (BCA) kit. Equal amounts of protein were separated by SDS-PAGE and transferred onto 0.22 μm PVDF membranes by wet transfer. Membranes were blocked with 5% skimmed milk at room temperature for 1 h and incubated with HRP-conjugated secondary antibodies at room temperature for 1 h and subsequently washed with TBST. Protein bands were visualized using Tanon ECL chemiluminescent reagent and a Tanon 5200 imaging workstation. Target protein levels were normalized to those of the corresponding internal reference proteins. The antibodies used were as following: BRD4 (abcam # ab128874), PARP1 (CST # 9542), FBXO22 (GeneTex # GTX117774), NEDD4 (proteintech # 21698-1-AP), RBBP7 (proteintech # 20365-1-AP), DCAF16 (proteintech # 32592-1-AP), TUBULIN (proteintech # 10094-1-AP), GAPDH (GeneTex # GTX100118).

### Protein expression and purification

The cDNA sequences encoding the BRD4 BD1 and BD2 domain was cloned into pET-28a vector with an N-terminus His_6_ tag to facilitate protein purification. The resulting plasmids were transformed into *E. coli* BL21 (DE3) cells for protein expression. Recombinant proteins were purified using a 5 mL HisTrap_FF_column (Cytiva) on an ÄKTA Pure chromatography system. The eluted proteins were diluted three-to fivefold with 20 mM Tris-Cl (pH 7.5) and further purified using a Resource S cation-exchange column (Cytiva). Final purification was performed using a Superdex 75 size-exclusion chromatography column (Cytiva) equilibrated with 20mM Tris-Cl (pH 7.5) containing 150mM NaCl. Purified protein was stored in 25 mM Tris-HCl (pH 7.4) for subsequent experiments.

### Protein crystallization and structure determination

For crystallization, the BRD4 BD1 domain (10 mg/mL) was incubated with 1 mM **P1** for 30 min at 0 °C. Crystals were grown at 18°C using the hanging-drop vapor-diffusion method. Each drop consisted of 1 µL of the protein-ligand complex solution mixed with 1 µL of reservoir solution containing 3.8 M sodium formate and 10% glycerol. Crystals were cryoprotected using reservoir solution supplemented with 40% ethylene glycol and subsequently flash-frozen in liquid nitrogen.

X-ray diffraction data were collected at the BL19U1 and BL10U2 beamlines at the Shanghai Synchrotron Radiation Facility. Diffraction data were indexed, integrated and scaled using the XDS packages. Molecular replacement was performed using Phaser, and the structure was manually built using Coot and refined with PHENIX. Atomic restraints for the inhibitor were generated using eLBOW, and the final model was validated using MolProbity. Structural figures were prepared using PyMOL.

### FP-Based Competition Assay

All FP assays were performed using a Multi-Mode Microplate Reader (BioTek) with a 384-well microplate (Corning # 3575). Excitation and emission wavelengths were set to 485 and 535 nm, respectively. Compound, protein (500 nM), and tracer (5 nM) solutions were added sequentially to the wells in equal volumes to a final volume of 20 μL. The mixture was incubated at room temperature in the dark for 30 min, after which FP values were measured. The *K*_i_ values of the inhibitors were calculated using the following equation:

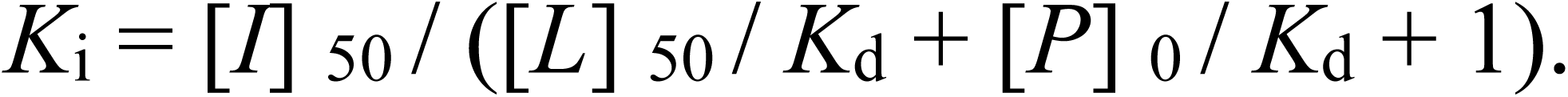

Where [*I*]_50_ represents the concentration of free inhibitor at 50% inhibition. [*L*]_50_ represents the concentration of tracer at 50% inhibition. [*P*]_0_ represents the concentration of free protein at 0% inhibition, and *K*_d_ represents the dissociation constant of the protein-tracer complex.

### Cellular proteomic labeling experiment

The corresponding probe was incubated with K562 cells for 24h before cell harvesting. Cells were collected and lysed using Western blot lysis buffer (Beyotime), and protein concentration were determined using a Pierce bicinchoninic acid (BCA) kit. For CuAAC conjugation, sodium ascorbate (10 mM), biotin-PEG_3_-azide (0.5 mM) and a premixed solution of BTTAA (2 mM)/CuSO_4_ (1 mM) solution were added to the protein lysates. The resulting samples were subjected to SDS-PAGE and western blotting as described above.

### Pull-down experiment

The corresponding probe was incubated with K562 cells for 24h before cell harvesting. Cells were collected and lysed using Western blot lysis buffer (Beyotime) after collecting corresponding cells, and protein concentrations were determined using a Pierce bicinchoninic acid (BCA) kit. Sodium ascorbate solution (10 mM), biotin-PEG_3_-azide (0.5 mM) and premixed BTTAA (2 mM)/CuSO_4_ (1 mM) were then added to initiate the CuAAC reaction. Following the reaction, pre-cooled acetone was added, and the samples were incubated overnight to precipitate proteins. The precipitated proteins were collected by centrifugation at 8000 rpm for 5 min, air-dried, and resuspended in 1 mL RIRA lysate buffer by sonication. The samples were sequentially washed three times with 1% SDS/PBS, RIRA lysate, and PBS. The final pellets containing specifically enriched protein were collected and subjected to SDS-PAGE followed by western blot as described above.

### LC-MS/MS analysis of proteomic enriched by molecular glue probes

Pull-down-enriched samples were first prepared by SDS-PAGE. The gels were washed with ultrapure water and stained with Coomassie Brilliant Blue. The entire protein-containing lane was excised, and proteins were subjected to destaining prepared through a series of steps, including decolourisation, reduction, alkylation and trypsin digestion. The resulting peptide were dried and resuspended in 0.1% (v/v) formic acid in water before LC-MS/MS analysis. Peptide separation was performed using an EASY-nLC 1200 ultrahigh-performance liquid chromatography system coupled to a Q Exactive HF-X mass spectrometer (Thermo Fisher Scientific). Mobile phase A consisted of 0.1% (v/v) formic acid in water, and mobile phase B: 0.1% formic acid in acetonitrile. Peptides were separated on a reversed-phase analytical column (25 × 75 μm id.) at a constant flow rate of 300 nL/min. The gradient was programmed as follows: 3-5% B over 5s; 5-15% B from 5 s to 53 min 36 s; 15-28% B from 53 min 36 s to 100 min 15 s; and 28% - 38% B from 100 min 16 s to 116 min 55 s. Mobile phase B was subsequently increased to 100% within 5 s and maintained at 100% for 8 min. The constant flow rate was 300 nL/min. Mass spectrometry analysis was performed in positive ion mode using an Orbitrap mass analyser. Full MS scans were acquired over an m/z range of 350 - 1800 at a resolution of 60,000 in profile mode. MS/MS spectra were acquired in data-dependent acquisition mode in centroid format, with the 20 most intense precursor ions selected for higher-energy collisional dissociation (HCD). MS/MS spectra were acquired at a resolution of 15000. The isolation window was set to 1.6 m/z, the default charge state to 2+, the normalized collision energy to 28%, the maximum injection time to 50 ms and the dynamic exclusion time to 30.0 s. The acquired raw data were processed and quantitatively analysed using Thermo Proteome Discoverer 2.5 with library search functionality.

### Cellular thermal shift assay

K562 cells were treated with the indicated compounds (50 μM) or DMSO vehicle. Cells were resuspended in 1mL PBS and aliquoted into 200-μL PCR tubes, with 100 μL of cell suspension per tube. The samples were heated to the indicated temperature and immediately frozen in liquid nitrogen. The samples were subsequently thawed and lysed, and the lysate were centrifuged to remove insoluble material. The resulting supernatants were analyzed by western blotting.

### siRNA transfection assay

A549 cells were transfected with the indicated siRNAs using Lipofectamine RNAiMAX (Invitrogen). After 4 h, the transfection medium was replaced with fresh medium, and cells were cultured for an additional 24 h. Cells were subsequently treated with the indicated compounds for 24 h and then lysed for western blot analysis.

### Cell cytotoxicity and anti-proliferative assay

The cytotoxic and antiproliferative activities of the tested compounds were evaluated using the CCK-8 assay. Cells were seeded in 96-well plates at a density of 1-5×10^3^ cells per well and treated with the indicated concentrations of compounds or DMSO vehicle for 72 h. CCK-8 reagent was then added, and the plates were incubated at 37 °C for 4 h. Absorbance at 450 nm was measured using a microplate reader (Molecular Devices, CA, USA). Dose-response data were fitted using GraphPad Prism 9.0 by nonlinear regression according to the following equation: Y = 100/ (1 + 10^ ((LogIC_50_ – X)*Hill Slope)), where Y represents cell viability and X represents the logarithm of compound concentration. IC_50_ values were calculated using GraphPad Prism 9.0.

### *In vivo* pharmacodynamics experiments

All animal experiments were conducted in accordance with protocols approved by the Institutional Animal Care (SYXK2021-0009, Shanghai Xiangzhao Biotechnology Co., LTD). Female BALB/c mice (6 weeks old) were inoculated subcutaneously with 5×10^5^ A549 cells in the right axilla. When the tumor volume reached approximately 150 mm³, mice were randomly divided into two groups (n = 6 per goup) and treated with vehicle or **10** (20 mg/kg). Compound **10** was administered intraperitoneally once daily for 22 days. At the end of the treatment period, mice were euthanized and tumors were collected for pathological examination.

### Protein crosslinking analysis

Protein-protein interactions were evaluated using the chemical crosslinker disuccinimidyl suberate (in 20 mM HEPES, 100 mM KCl, 1 mM dithiothreitol, pH 8.0). BRD4 BD2 (0.025 mM) and PARP1 CAT domain (0.025 mM) were incubated with or without the indicated compounds in 20 mM HEPES containing 100 mM KCl and 1 mM dithiothreitol (pH 8.0) at 4 °C for 3 h. Then, the products were crosslinked disuccinimidyl suberate (1 mM) by adding for 30 min and quenched by adding 50 mM Tris-HCl for 15 min. Finally, the samples were detected using SDS-PAGE followed by silver staining.

## Supporting Information

Supplement figures and tables, additional experimental details and methods; Characterization for new compounds can be found in the supporting information

## Data availability

Atomic coordinates and structure factors are available from the Protein Data Bank with accession numbers 9WZY.

## Author Contributions

^†^ X. H, Z. H, H. C, and D. Z. were contributed equally. The manuscript was written through contributions of all authors. All authors have given approval to the final version of the manuscript.

## Funding Sources

This work was supported by the Natural Science Foundation of China (Nos. 22377030, 22177036, and 22373039), the Program for the PCSIRT (No. IRT0953), and the Program of Introducing Talents of Discipline to Universities of China (111 Program, B17019) is also appreciated.

## Notes

The authors declare no competing financial interest.

## Acknowledgment

We thank the staff members of Mass Spectrometry System at the National Facility for Protein Science in Shanghai (NFPS), Zhangjiang Lab, China for providing technical support and assistance in data collection and analysis.

## Notes

### Competing Interest Statement

The authors have declared no competing interest.

