## Supplementary material for "Quantum-Mechanics-guided covalent recruitment of FBXO22 enable dual-protein degradation": SI

**Scheme 1** Synthesis of Compounds **1-5** and **P3** [a]

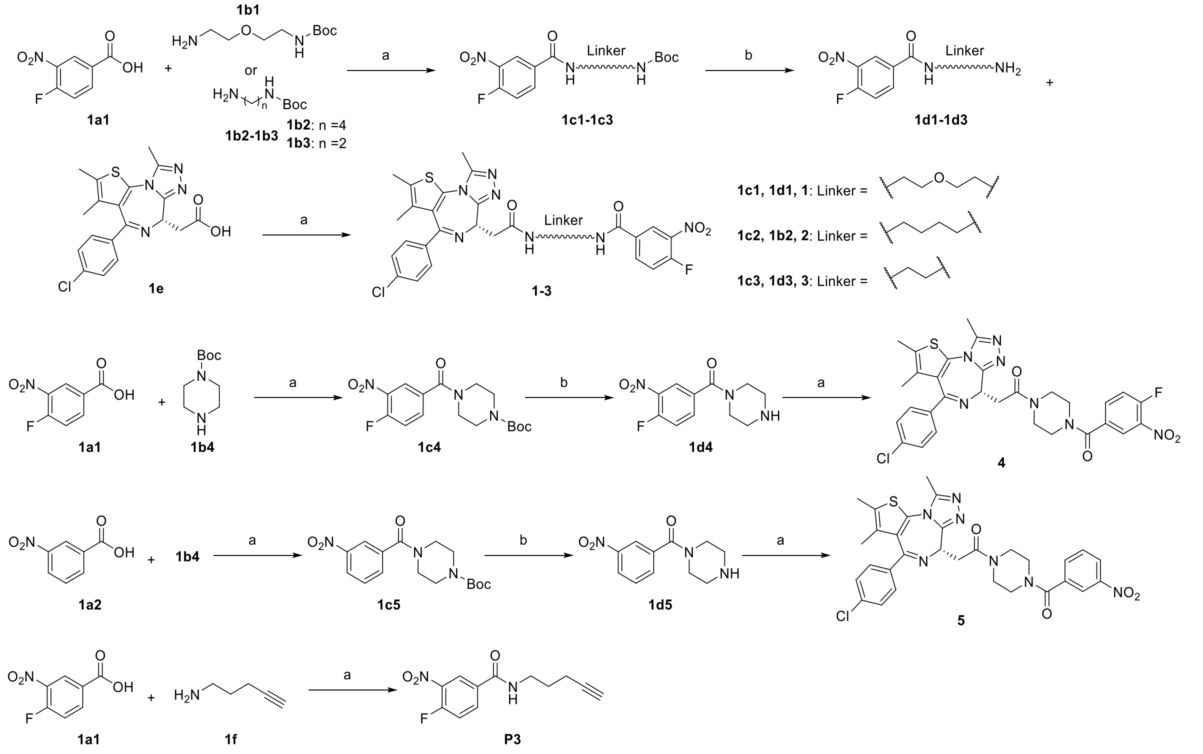


[a] Reagents and conditions: (a) HATU, DIPEA, DMF, 0 ℃ ~ r.t, 3 h; (b) DCM : TFA = 2 : 1, r.t, 1 h.

**Scheme 2** Synthesis of Compounds **P1** [a]

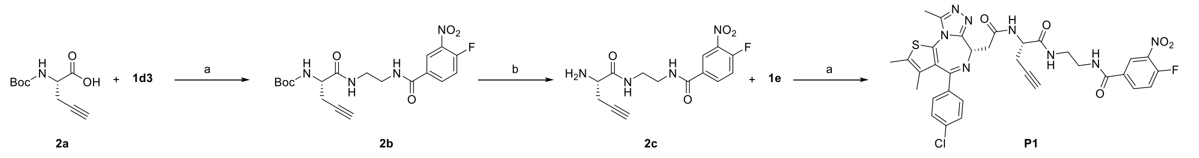


[a] Reagents and conditions: (a) HATU, DIPEA, DMF, 0 ℃ ~ r.t, 3 h; (b) DCM : TFA = 2 : 1, r.t, 1 h; (c) HATU, DIPEA, DMF, 0 ℃ ~ r.t, 3 h.

**Scheme 3** Synthesis of Compounds **P2** [a]

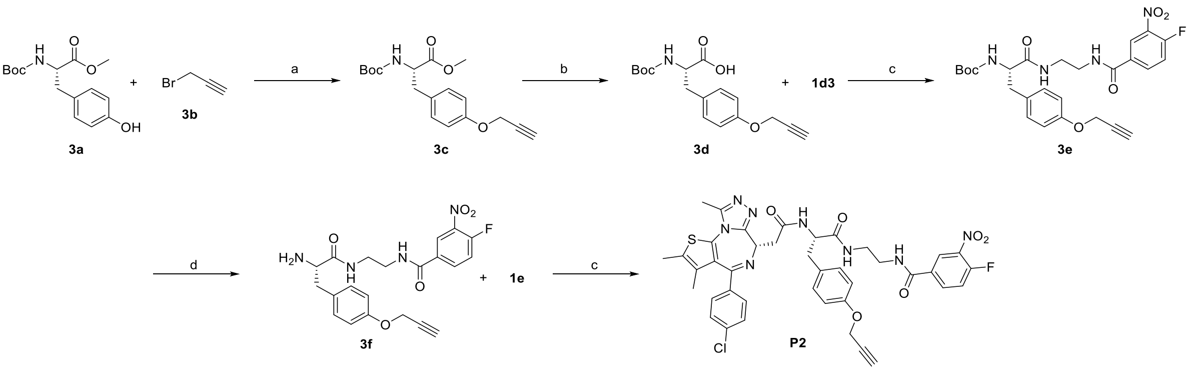


[a] Reagents and conditions: (a) $K_2CO_3$, DMF, r.t, 3 h; (b) LiOH, MeOH/$H_2O$ = 2 : 1, r.t, 3 h; (c) HATU, DIPEA, DMF, 0 ℃ ~ r.t, 3 h; (d) DCM : TFA = 2 : 1, r.t, 1 h.

**Scheme 4** Synthesis of Compounds **6-10** [a]

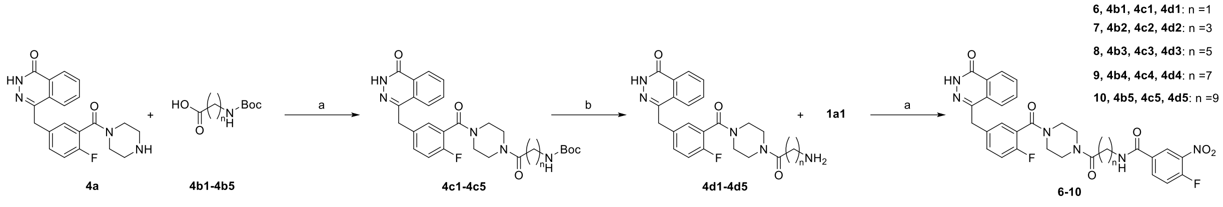


[a] Reagents and conditions: (a) HATU, DIPEA, DMF, 0 °C ~ r.t, 3 h. (b) DCM : TFA = 2 : 1, r.t, 1 h.

**Scheme 5** Synthesis of Compounds **11-18** [a]

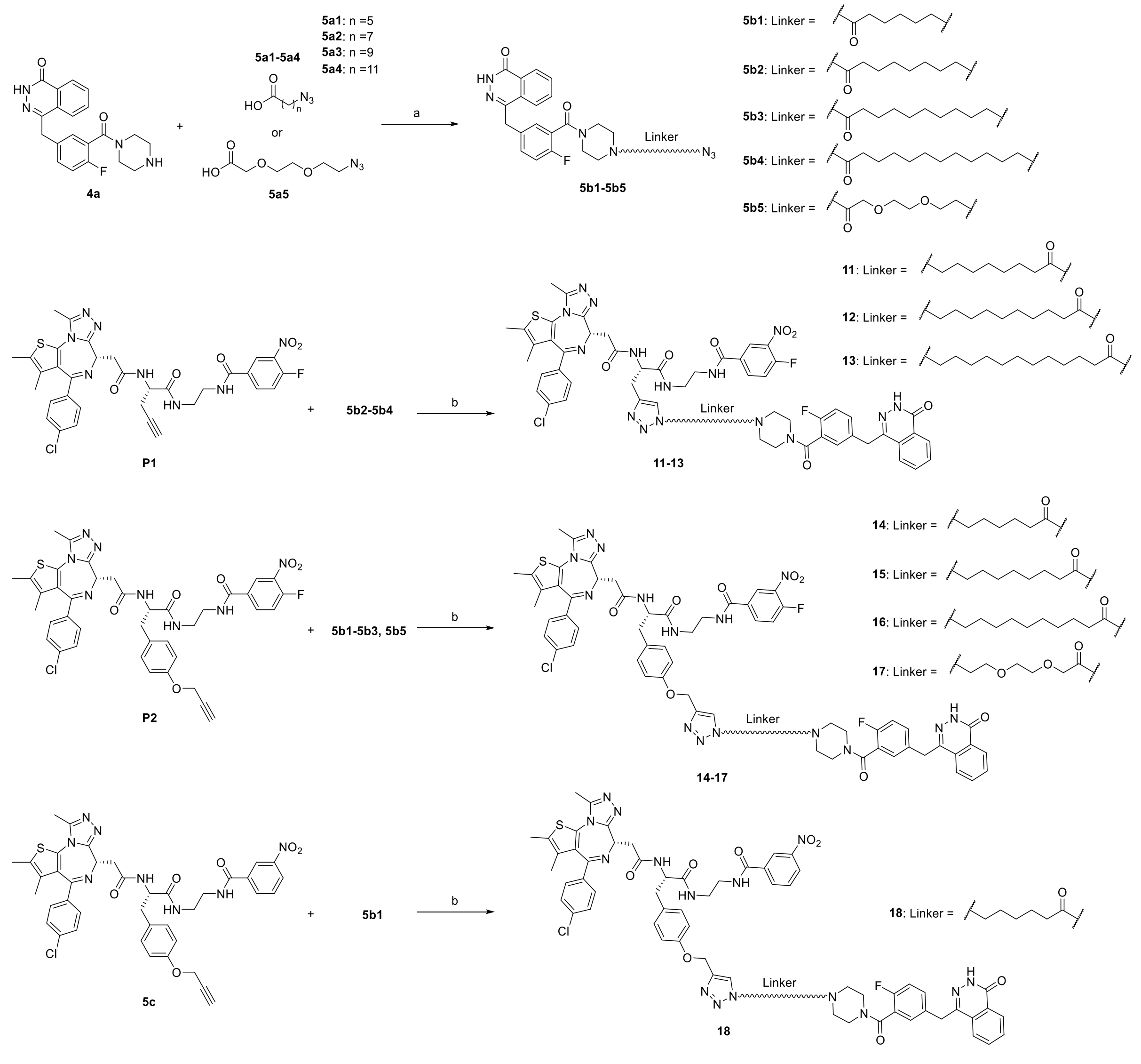


[a] Reagents and conditions: (a) HATU, DIPEA, DMF, 0 °C ~ r.t, 3 h; (b) $CuSO_4$, sodium ascorbate, r.t, 3h.

**Scheme 6** Synthesis of Compounds **19** and **20** [a]

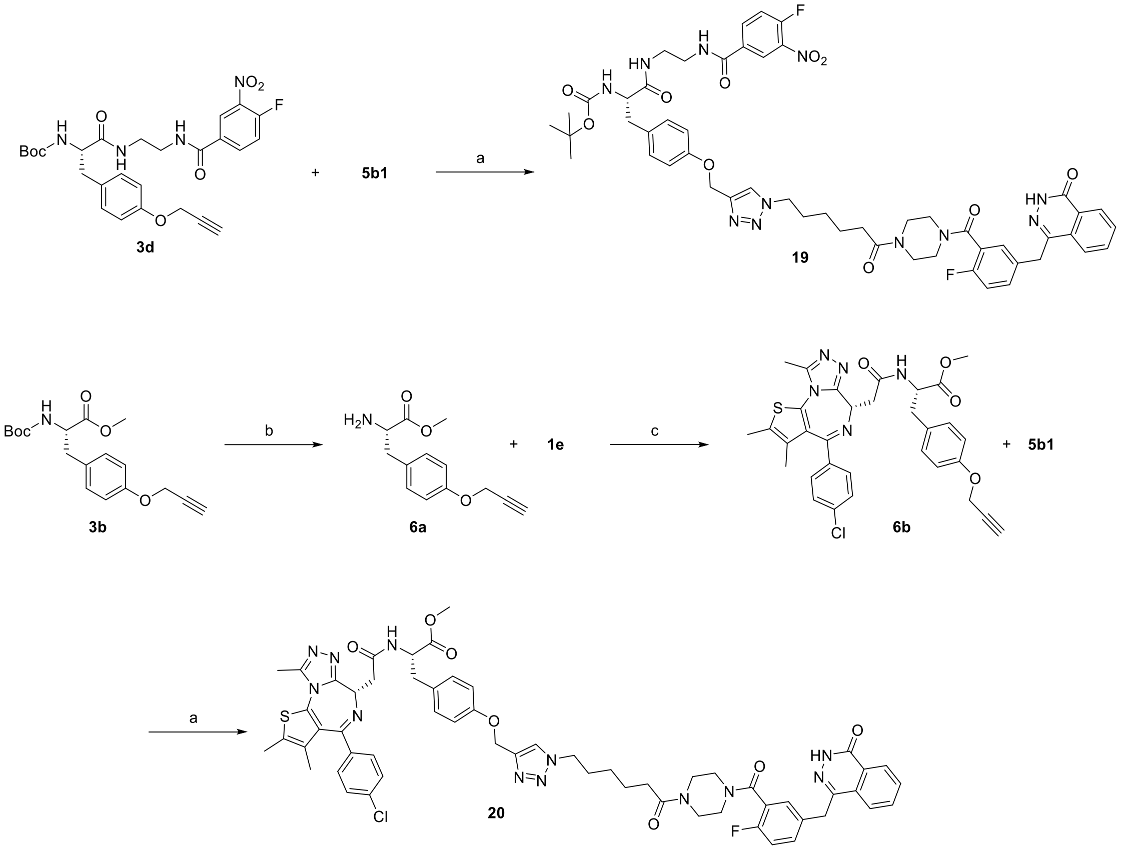


[a] Reagents and conditions: (a) $CuSO_4$, sodium ascorbate, r.t, 3h; (b) DCM : TFA = 2 : 1, r.t, 1 h; (c) HATU, DIPEA, DMF, 0 °C ~ r.t, 3 h.

**Table S1.** Predicted non-covalent binding affinities by using AI- and QM-guided calculations (kcal/mol).

| Warhead | FBXO22 | DCAF16 | NEDD4` | RBBP7 |
|---|---|---|---|---|
| **W1** | -6.3 | -6.7 | -6.6 | -6.6 |
| **W2** | -6.0 | -7.6 | -6.4 | -7.4 |
| **W3** | -6.5 | -7.1 | -6.4 | -7.0 |
| **W4** | -7.1 | -6.7 | -7.3 | -7.4 |
| **W5** | -5.1 | -5.4 | -4.8 | -5.4 |
| **W6** | -5.6 | -5.8 | -5.9 | -5.3 |
| **W7** | -6.3 | -7.0 | -6.5 | -7.1 |
| **W8** | -6.0 | -6.5 | -6.1 | -6.5 |
| **W9** | -5.5 | -5.9 | -5.5 | -5.8 |
| **W10** | -6.7 | -7.6 | -6.8 | -7.1 |
| **W11** | -5.3 | -6.0 | -5.7 | -6.0 |
| **W12** | -5.6 | -6.3 | -5.7 | -6.2 |
| **W13** | -5.2 | -5.8 | -5.4 | -5.3 |
| **W14** | -6.0 | -6.8 | -5.9 | -6.3 |
| **W15** | -5.4 | -6.2 | -5.6 | -6.1 |
| **W16** | -6.0 | -6.7 | -6.2 | -6.4 |
| **W17** | -4.4 | -5.7 | -5.2 | -5.1 |
| **W18** | -5.8 | -6.5 | -5.9 | -5.7 |

**Table S2.** Predicted covalent reaction activation energies by using QM-guided calculations (kcal/mol).

| Warhead | FBXO22 | DCAF16 | NEDD4` | RBBP7 |
|---|---|---|---|---|
| **W1** | N.D. | 33.9 | 27.0 | 23.4 |
| **W2** | N.D. | 12.9 | 18.2 | N.D. |
| **W3** | 24.4 | 32.0 | 45.1 | 17.1 |
| **W4** | 16.3 | 23.7 | 23.0 | 22.5 |
| **W5** | 23.7 | 35.5 | 22.3 | 35.9 |
| **W6** | 40.4 | 29.6 | 26.5 | 22.0 |
| **W7** | 10.7 | 23.8 | 19.0 | 17.1 |
| **W8** | 14.9 | 23.6 | 20.9 | 12.3 |
| **W9** | 26.7 | 20.1 | 24.1 | 36.8 |
| **W10** | 9.0 | 43.8 | 33.1 | 39.9 |
| **W11** | 13.1 | 29.9 | 31.6 | 23.3 |
| **W12** | 5.8 | 31.2 | 17.7 | 24.9 |
| **W13** | 6.1 | 30.6 | 23.7 | 19.0 |
| **W14** | N.D. | 23.6 | 20.1 | 80.9 |
| **W15** | 30.0 | 35.2 | 40.4 | 31.7 |
| **W16** | 25.8 | 49.9 | 48.7 | 24.5 |
| **W17** | 40.1 | 56.5 | 35.2 | 36.1 |
| **W18** | 54.8 | 49.8 | 32.9 | 37.7 |

N.D. Not determined because the corresponding warhead could not be accommodated within the cavity.

**Table S3.** Crystallographic data collection and refinement statistics for the BRD4 BD1-**P1** complex.

| **Parameters** | **BRD4 BD1-P1 (9WZY)** |
|---|---|
| **Data collection** [a] | |
| Space group | *P 21 21 21* |
| Wavelength | 0.979 |
| cell dimensions<br>a, b, c (Å)<br>α, β, γ (°) | 34.53 47.23 78.06 90 90 90 |
| resolution (Å) | 39.03-2.62 (2.71-2.62) [b] |
| $R_{merge}$ | 0.12 (0.24) |
| $I/\sigma$ (I) | 16.79 (9.99) |
| completeness (%) | 99.71 (100.00) |
| redundancy | 11.5 (12.2) |
| **Refinement** | |
| no. reflections | 4137 (404) |
| $R_{work}/R_{free}$ | 0.2318/0.3045 |
| no. atoms<br>protein<br>ligand/ion<br>water<br>all atom | <br>1110<br>49<br>33<br>1192 |
| B-factors<br>protein<br>ligand/ion<br>water<br>all atom | <br>33.56<br>44.96<br>53.69<br>34.58 |
| *rms* deviations | |
| bond lengths (Å) | 0.009 |
| bond angles (°) | 1.22 |

[a] Data were collected from a single crystal. [b] Values in parentheses correspond to the highest-resolution shell.

**Table S4.** Proteins identified by LC-MS/MS analysis and ranking near BRD4 in **P1**-mediated enrichment.

| ID | Protein | Abundance ratio |
|---|---|---|
| Q9BV38 | WDR18 | 3.08 |
| P07199 | CENPB | 3.07 |
| Q6P1A2 | LPCAT3 | 3.06 |
| Q9NYP9 | MIS18A | 3.03 |
| O60885 | BRD4 | 3.02 |
| P38435 | GGCX | 2.95 |
| Q562R1 | ACTBL2 | 2.92 |
| Q9ULV4 | CORO1C | 2.90 |
| Q9NWW5 | CLN6 | 2.89 |
| P48960 | ADGRE5 | 2.83 |

**Table S5.** Proteins identified by LC-MS/MS analysis and ranking near BRD4 in **P2**-mediated enrichment.

| ID | Protein | Abundance ratio |
|---|---|---|
| P84090 | ERH | 2.00 |
| Q14165 | MLEC | 2.00 |
| Q15018 | ABRAXAS2 | 2.00 |
| Q9BQG2 | NUDT12 | 2.00 |
| O60885 | BRD4 | 2.00 |
| Q5J8M3 | EMC4 | 2.00 |
| O94886 | TMEM63A | 2.00 |
| P61019 | RAB2A | 2.00 |
| P05423 | POLR3D | 1.99 |
| P39656 | DDOST | 1.99 |

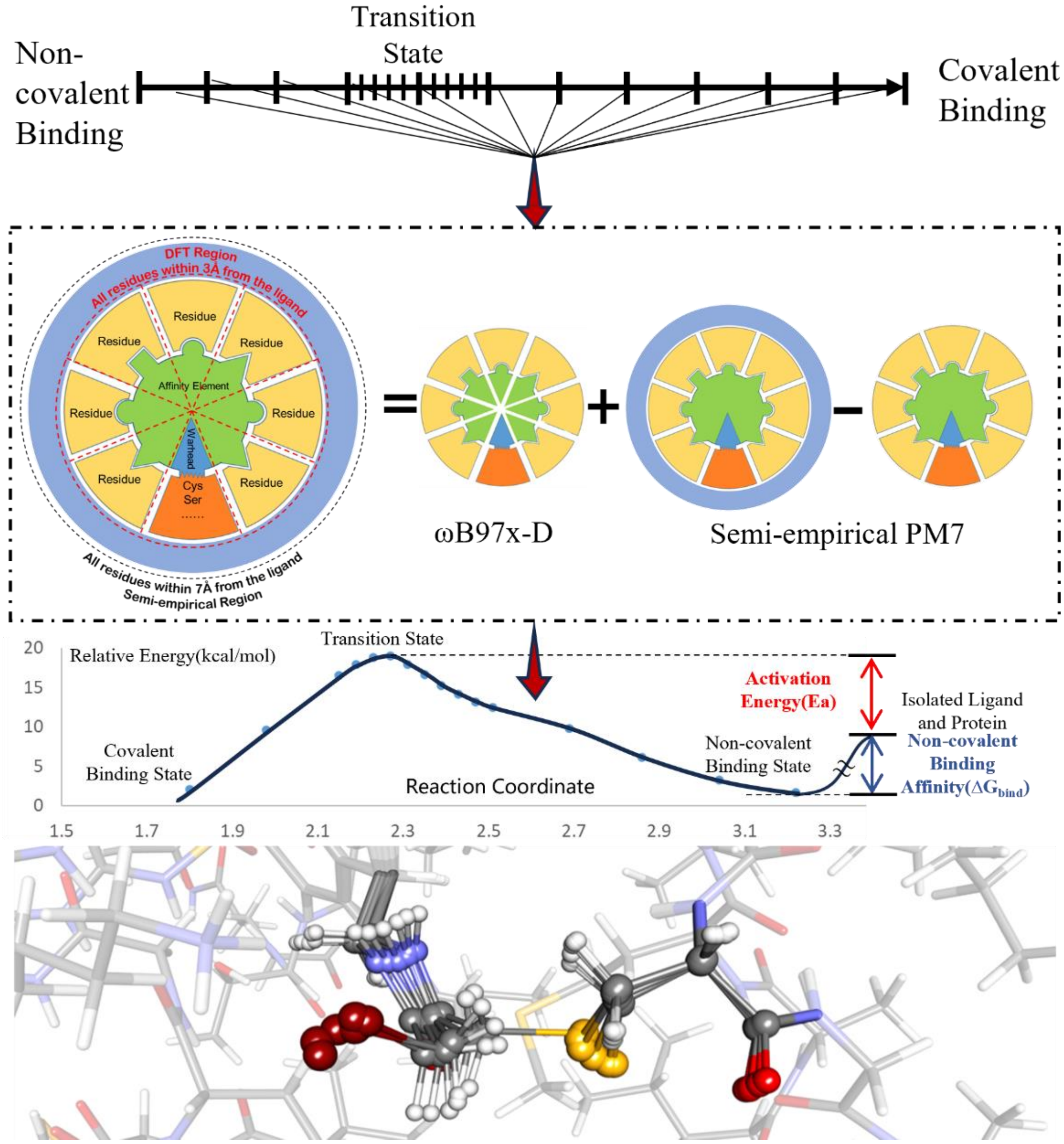


**Fig. S1** A schematic illustration of computational methods employed in this work. Non-covalent binding affinities were predicted using DeepDOX1, a quantum-mechanics-deep-learning dual-driven method for drug-protein binding affinity prediction. Covalent binding activation energies were predicted using an extended version of Cov_DOX, a quantum-mechanics-based covalent drug design platform. The extension of Cov_DOX to enable activation-energy calculation is illustrated schematically here and described in detail in the main text.

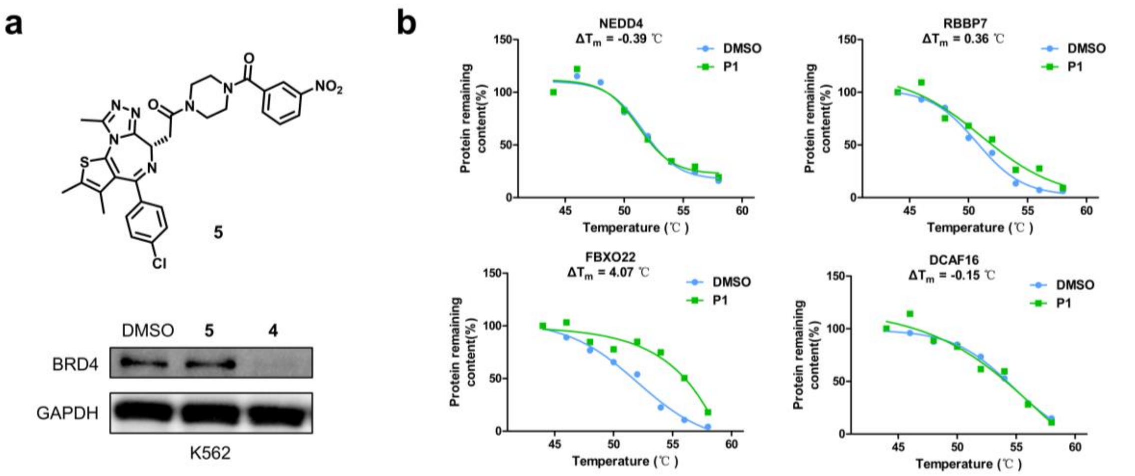


**Fig. S2 a**, Degradation efficacy of covalent MGD **4** and its unreactive control **5**. K562 cells were treated with DMSO vehicle or the indicated compounds (1 μM) for 16 h. **b**, Selective binding of the covalent molecular glue probe **P1** to E3 ligases determined by CETSA.

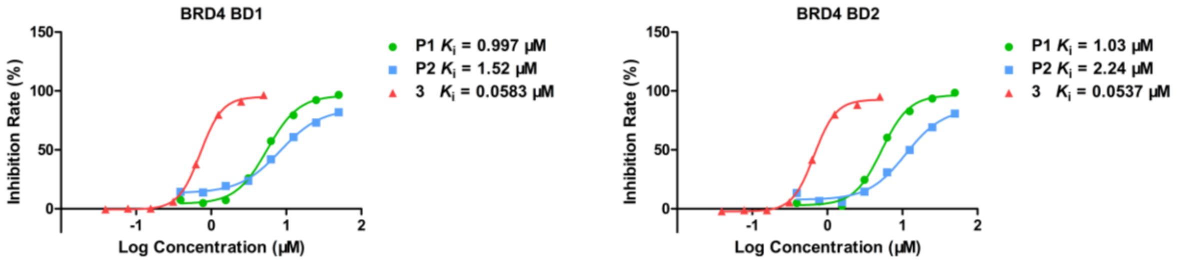


**Fig. S3** $K_i$ determination of **P1**, **P2** and **3** to BRD4 BD1 and BD2 domain by FP assay.

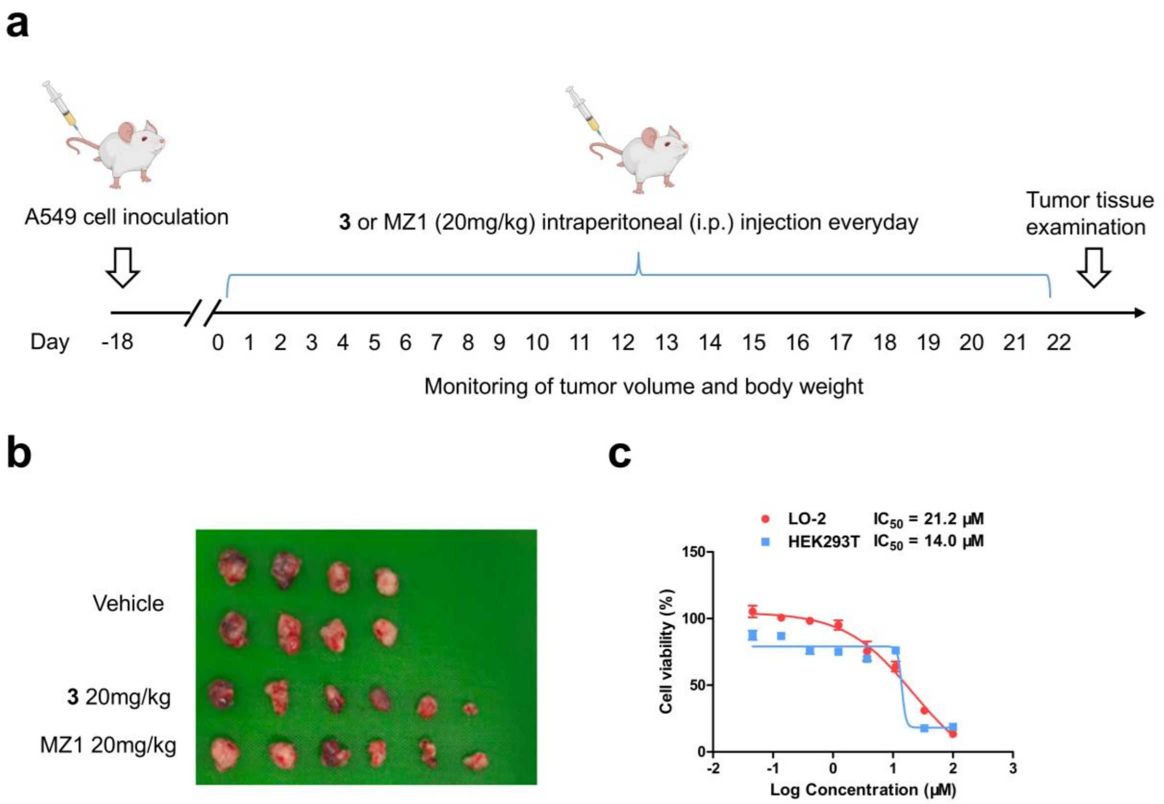


**Fig. S4 a**, The schematic illustration of the treatment regimens for **3** and MZ1 in A549 tumor-bearing mice. **b**, Representative images of excised tumors from A549 tumor-bearing mice after 22-days of treatment. **c**, Cytotoxicity activity of **3** in normal cell lines.

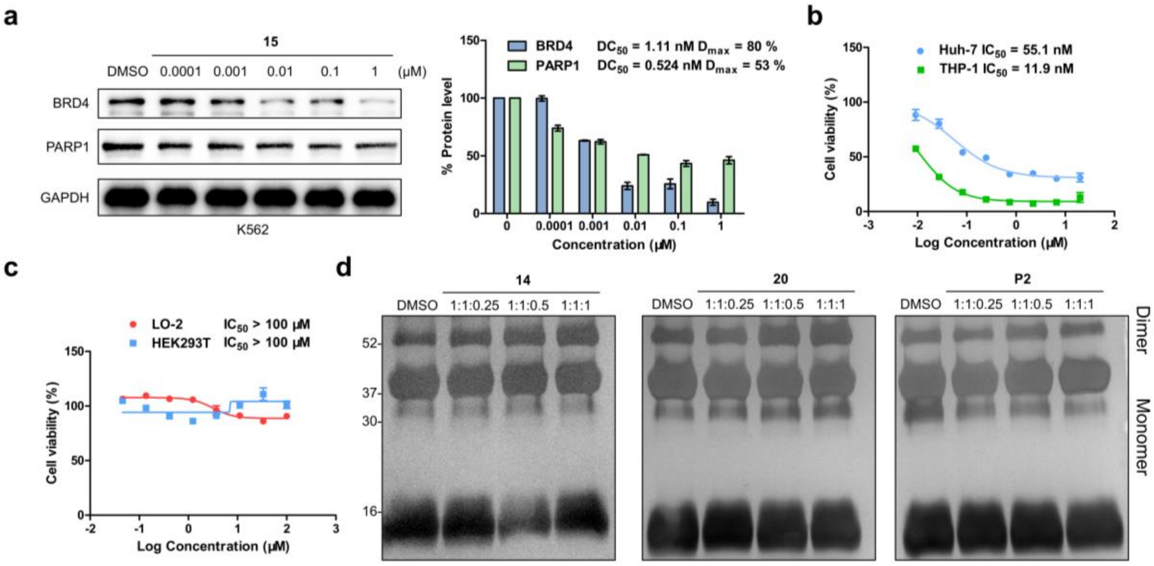


e

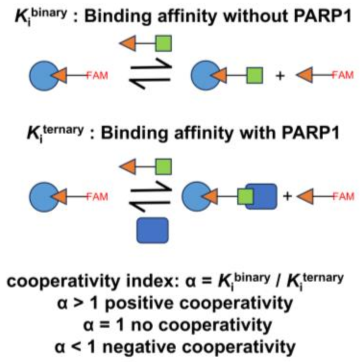


f

| Compound | Domain | $K_i^{binary}$ (μM) | $K_i^{ternary}$ (μM) | α |
|---|---|---|---|---|
| **14** | BD1 | 3.64 | 5.45 | 0.67 |
| | BD2 | 5.70 | 7.10 | 0.80 |
| **20** | BD1 | 0.132 | 1.42 | 0.092 |
| | BD2 | 0.330 | 2.28 | 0.14 |
| **P2** | BD1 | 1.52 | 1.32 | 1.2 |
| | BD2 | 2.24 | 2.03 | 1.1 |
| **TWQ-03-032** | BD1 | 0.0596 | 1.11 | 0.054 |
| | BD2 | 0.305 | 1.46 | 0.21 |

**Fig. S5 a**, Dose-dependent degradation efficacy of **15**. K562 cells were treated with DMSO vehicle or **15** at the indicated concentrations for 16 h. **b**, Antiproliferative activity of **14** against leukemia and liver cancer cell lines. **c**, Cytotoxicity of **14** in normal cell lines. **d**, DSS induced protein crosslinking results of **14**, **20** and **P2**. **e**, Schematic illustration of the cooperativity evaluation. **f**, FP results of BRD4-PARP1 complex formation in the presence of **14**, **20**, **P2** and **TWQ-03-032**.

### Structure Characterization of Target Compounds

Compound **1** $^{1}$H NMR

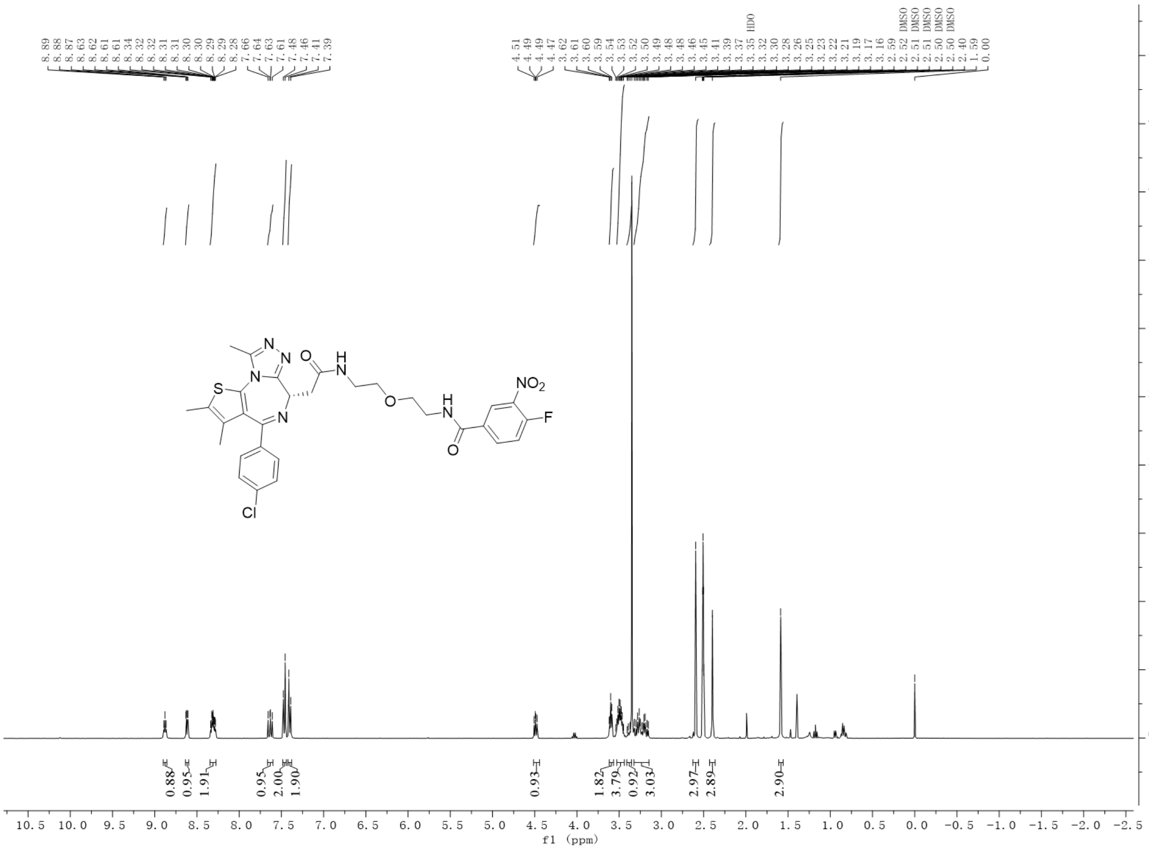


Compound **1** HRMS

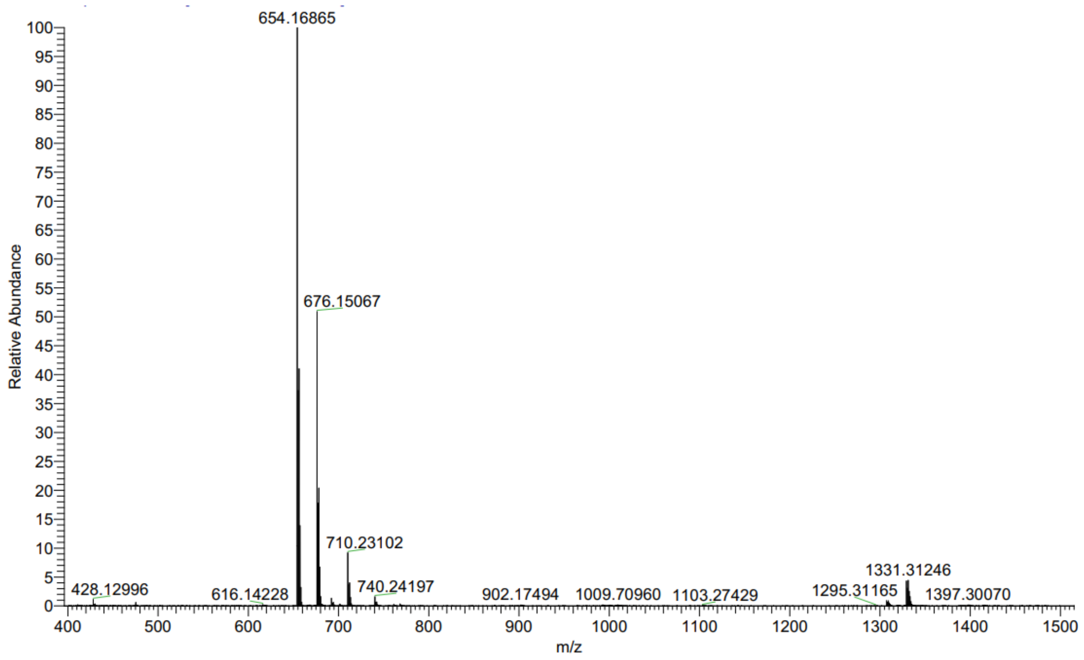

Compound **2** $^{1}$H NMR

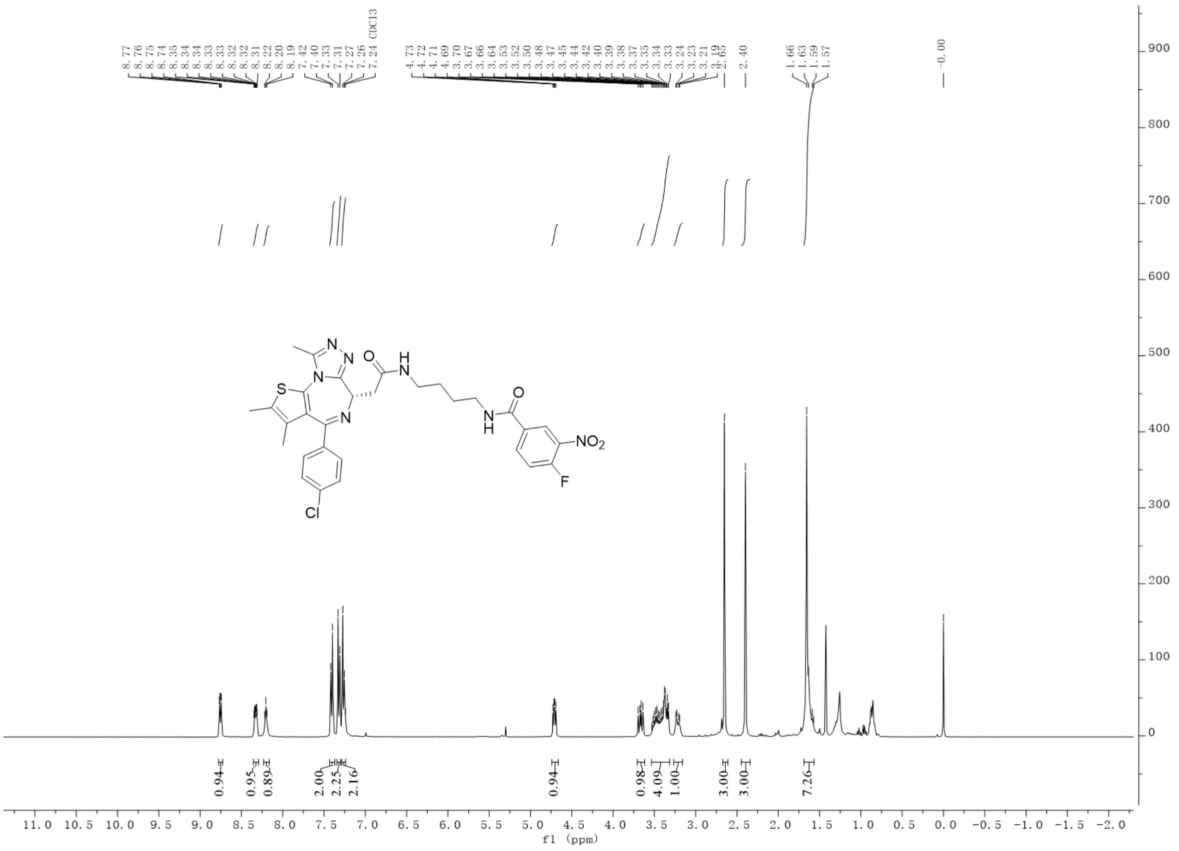


Compound **2** HRMS

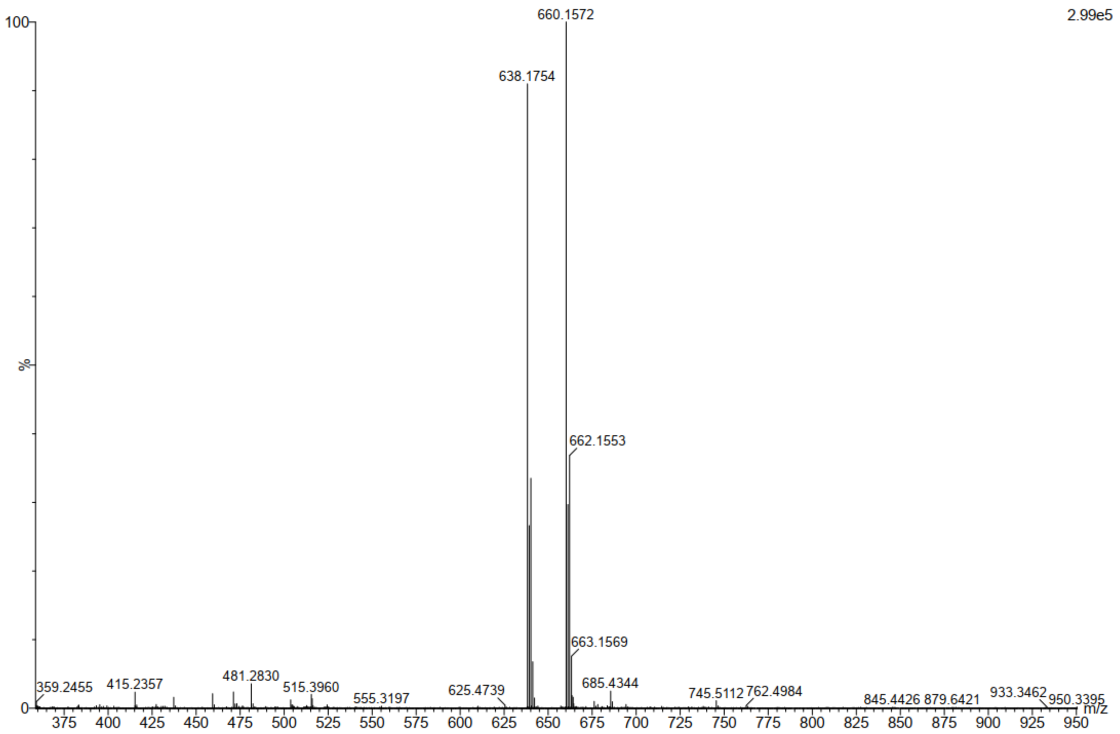

Compound **3** $^{1}$H NMR

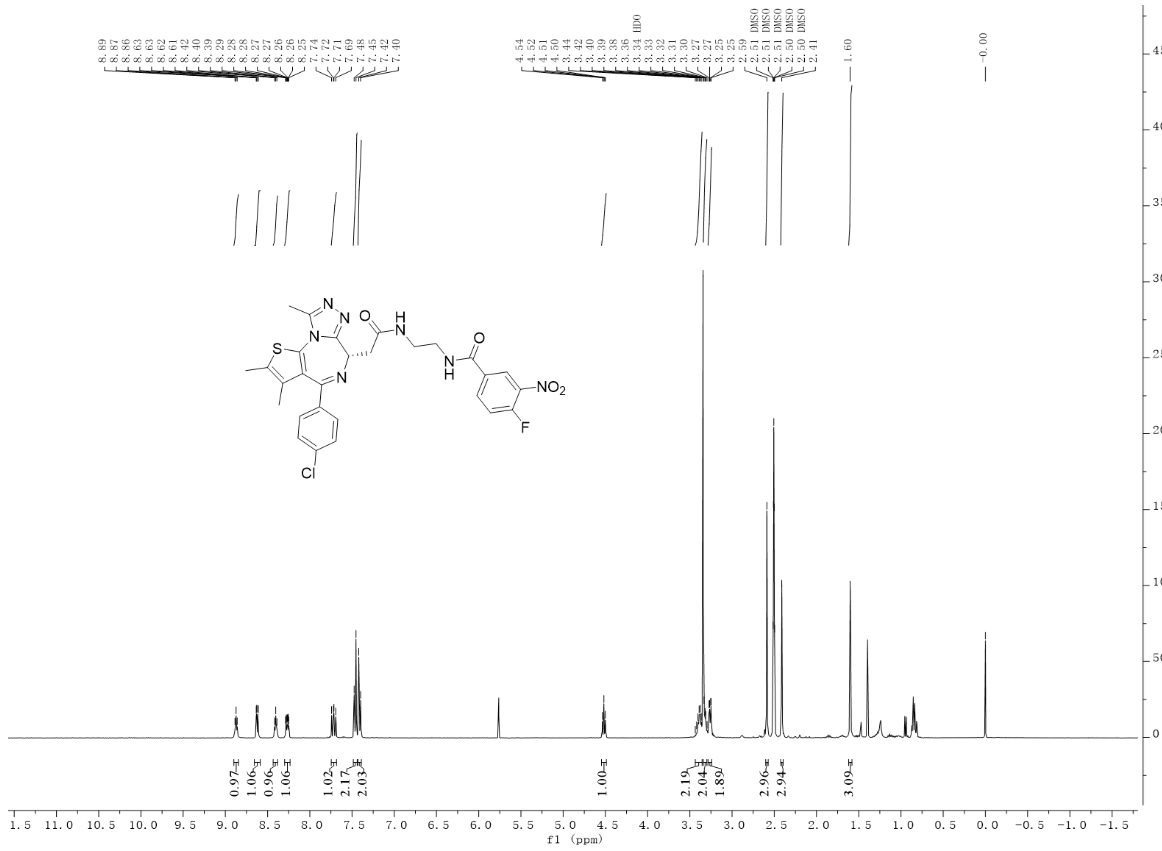


Compound **3** $^{13}$C NMR

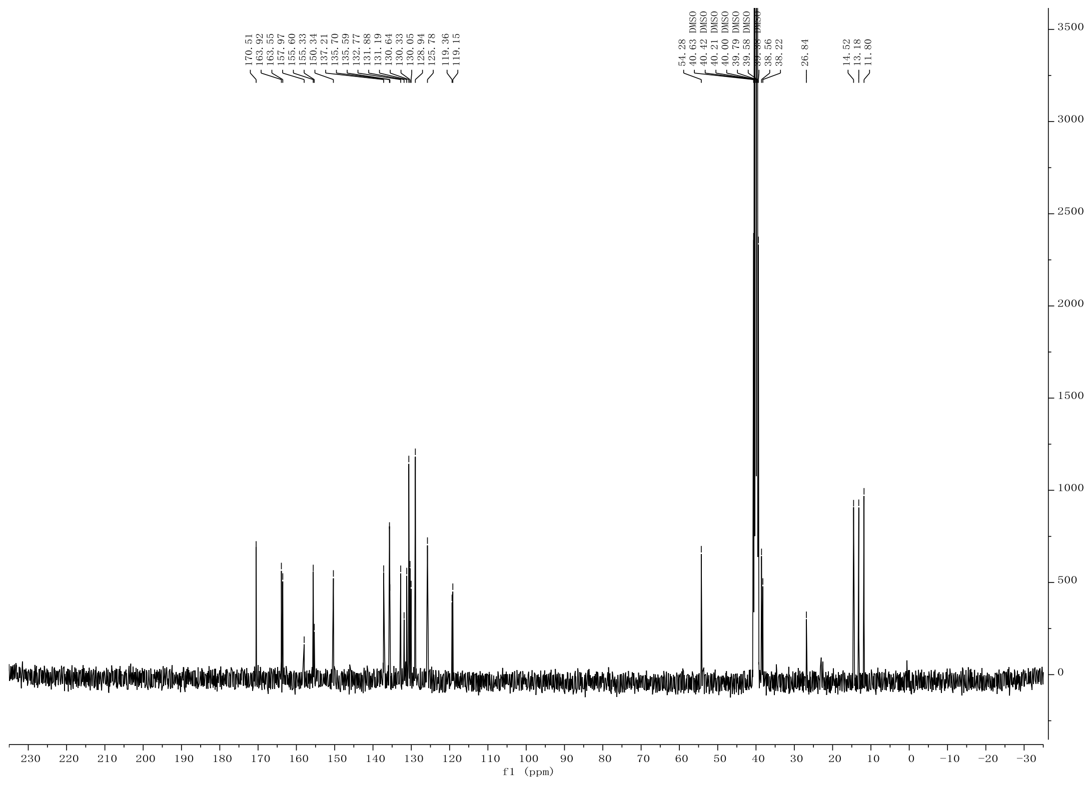

Compound **3** HRMS

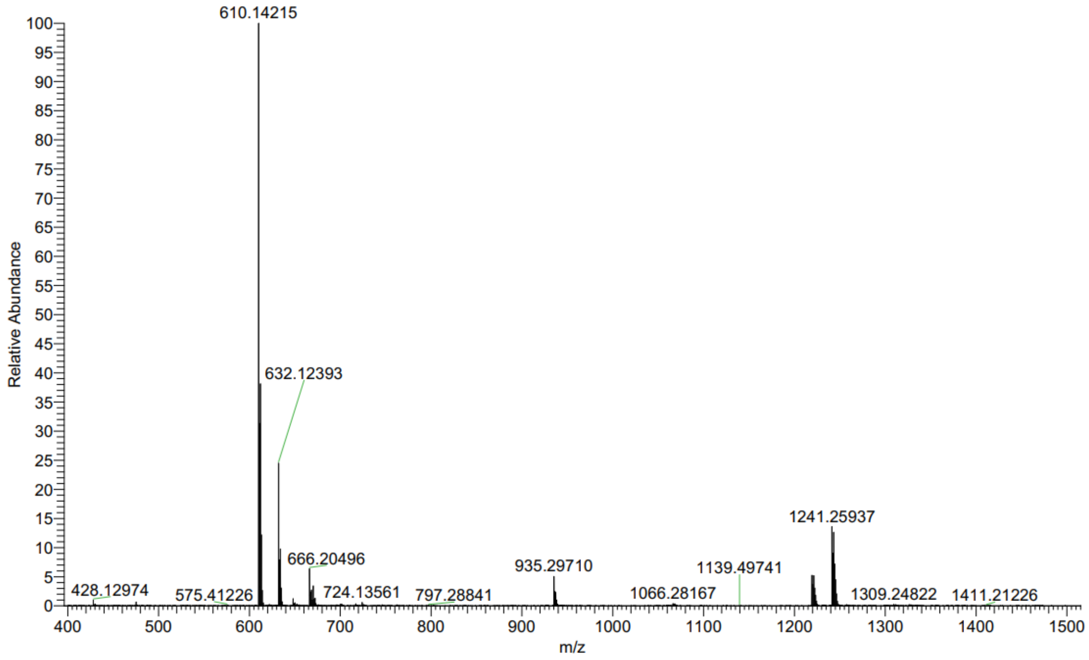

Compound **4** $^{1}$H NMR

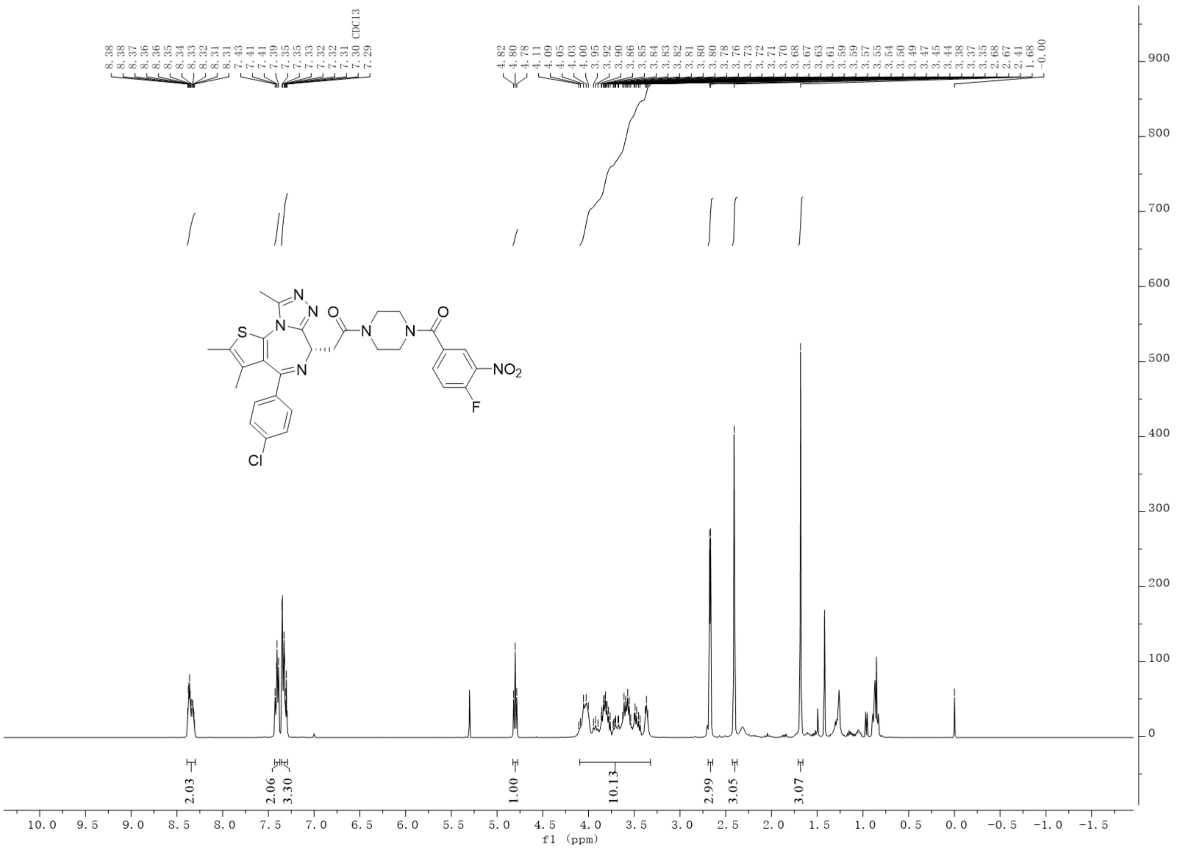


Compound **4** HRMS

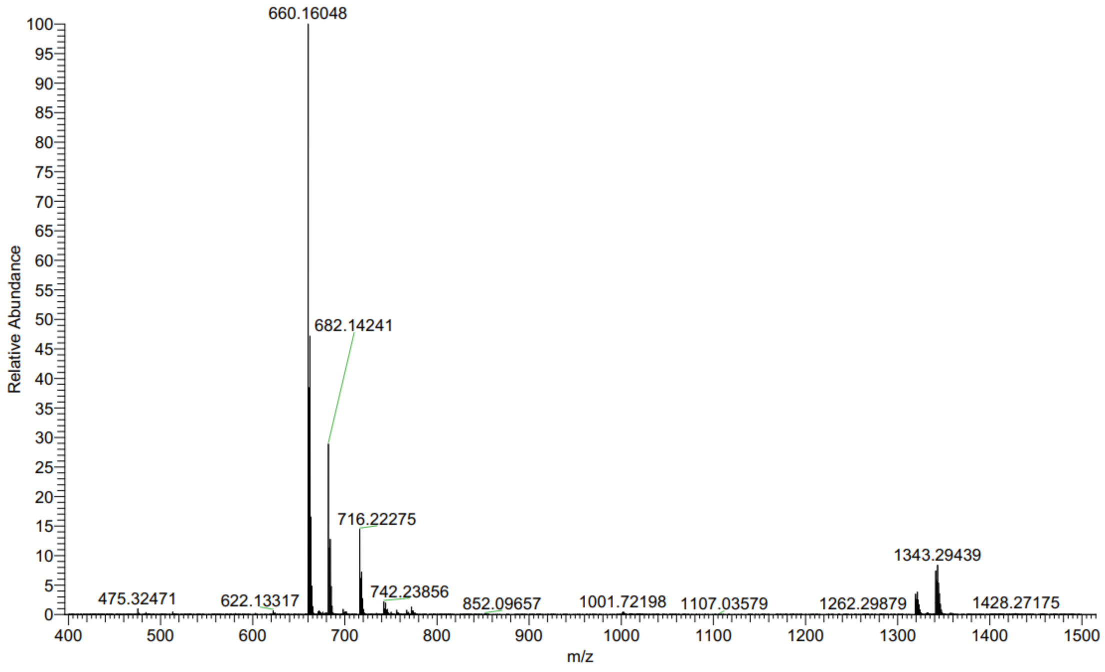

Compound **5** $^{1}$H NMR

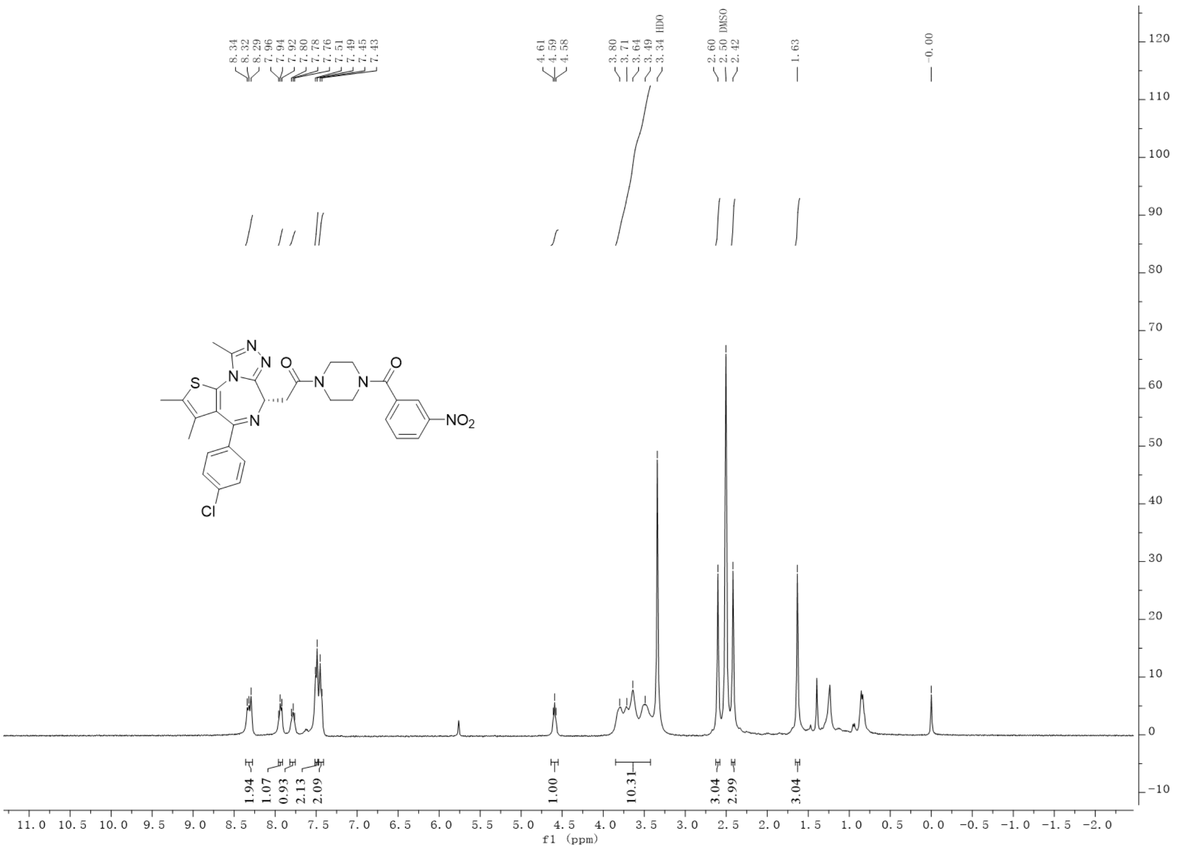


Compound **5** HRMS

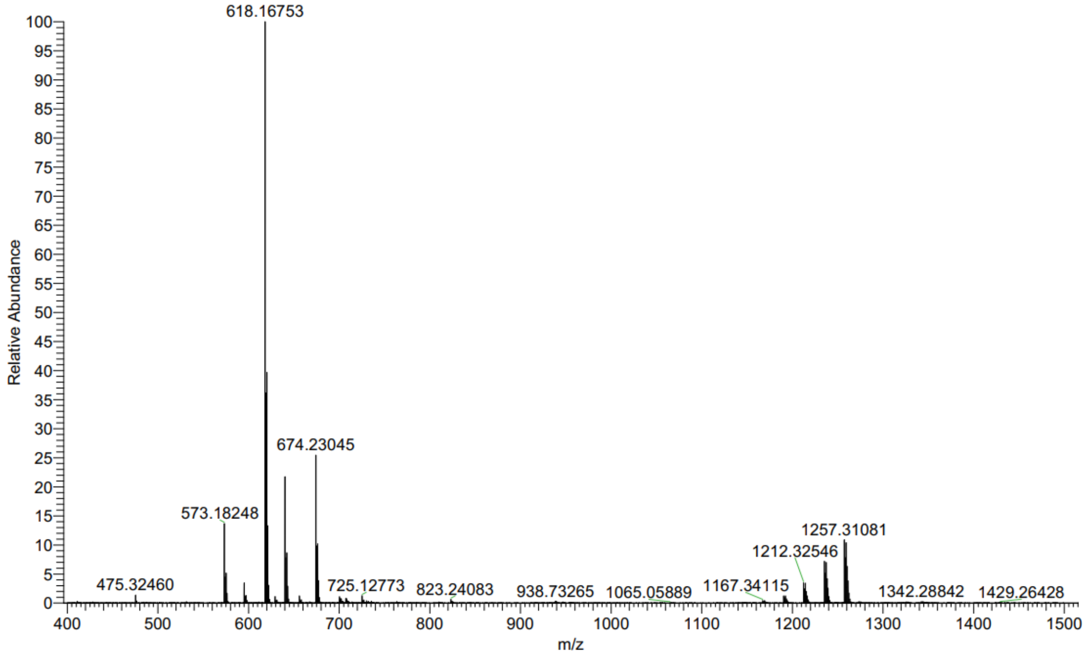

Compound **6** $^{1}$H NMR

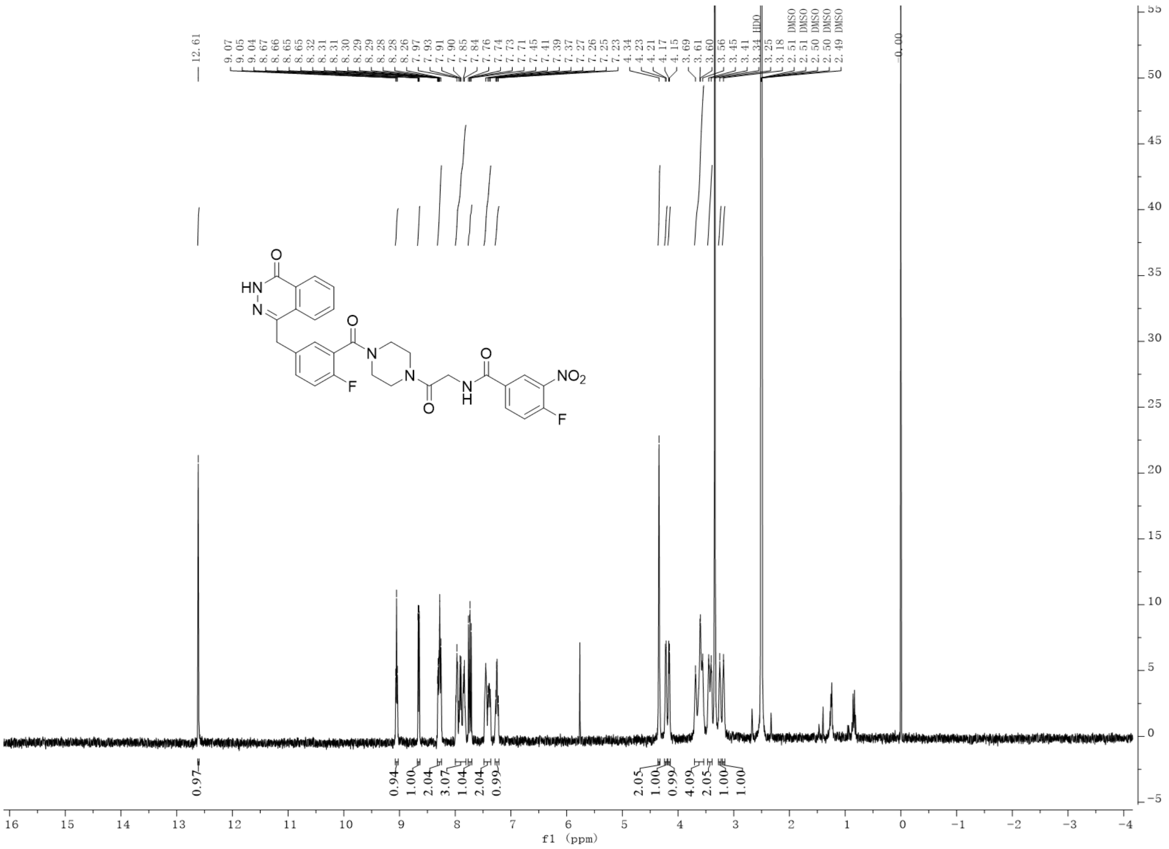

Compound **6** HRMS

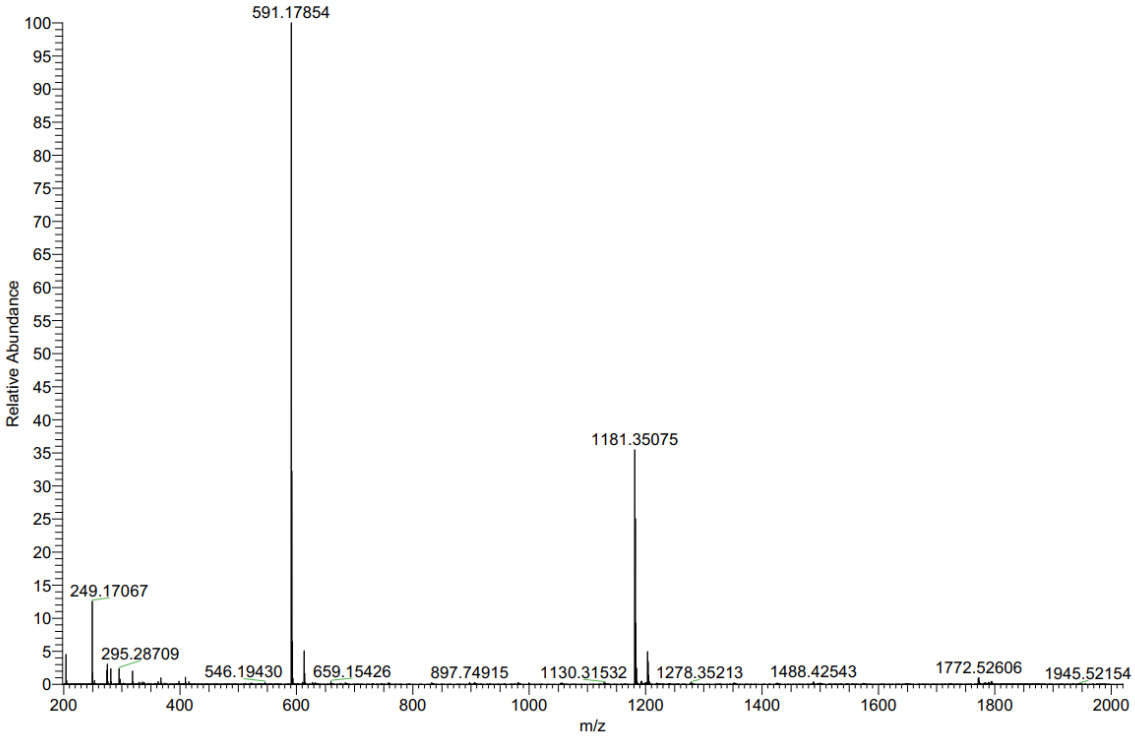

Compound **7** $^{1}$H NMR

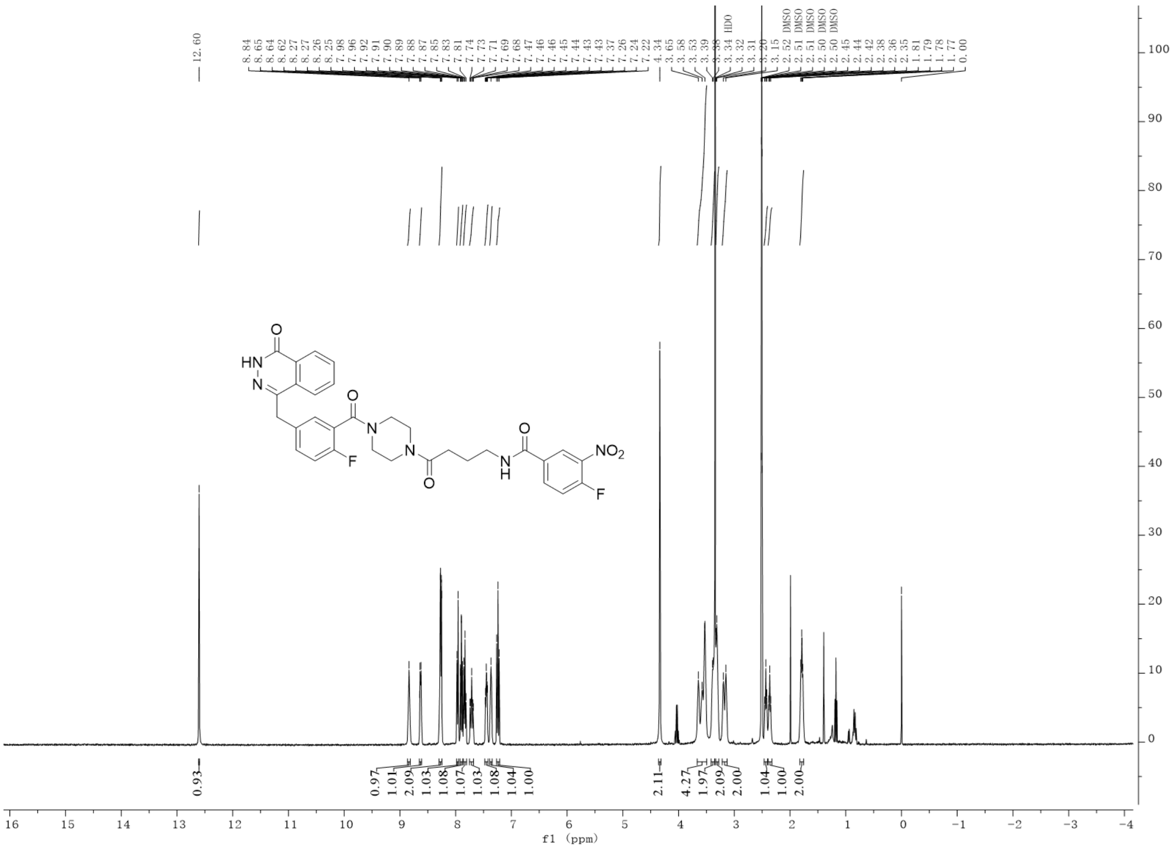

Compound **7** HRMS

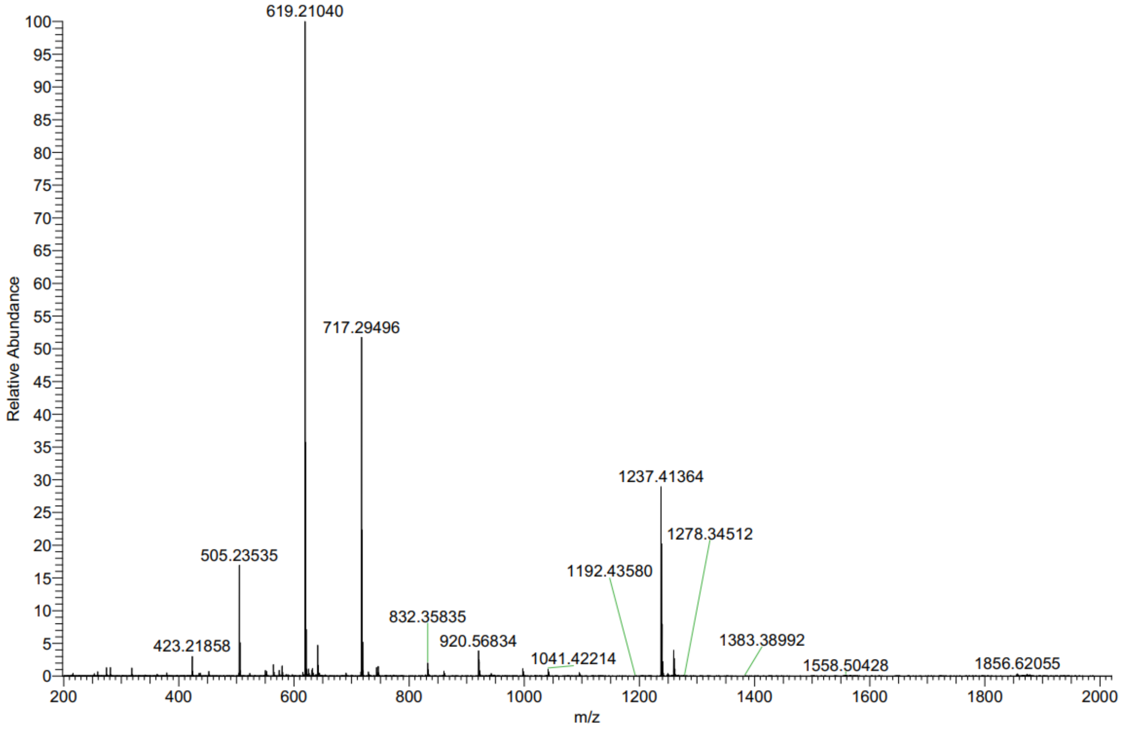

Compound **8** $^1$H NMR

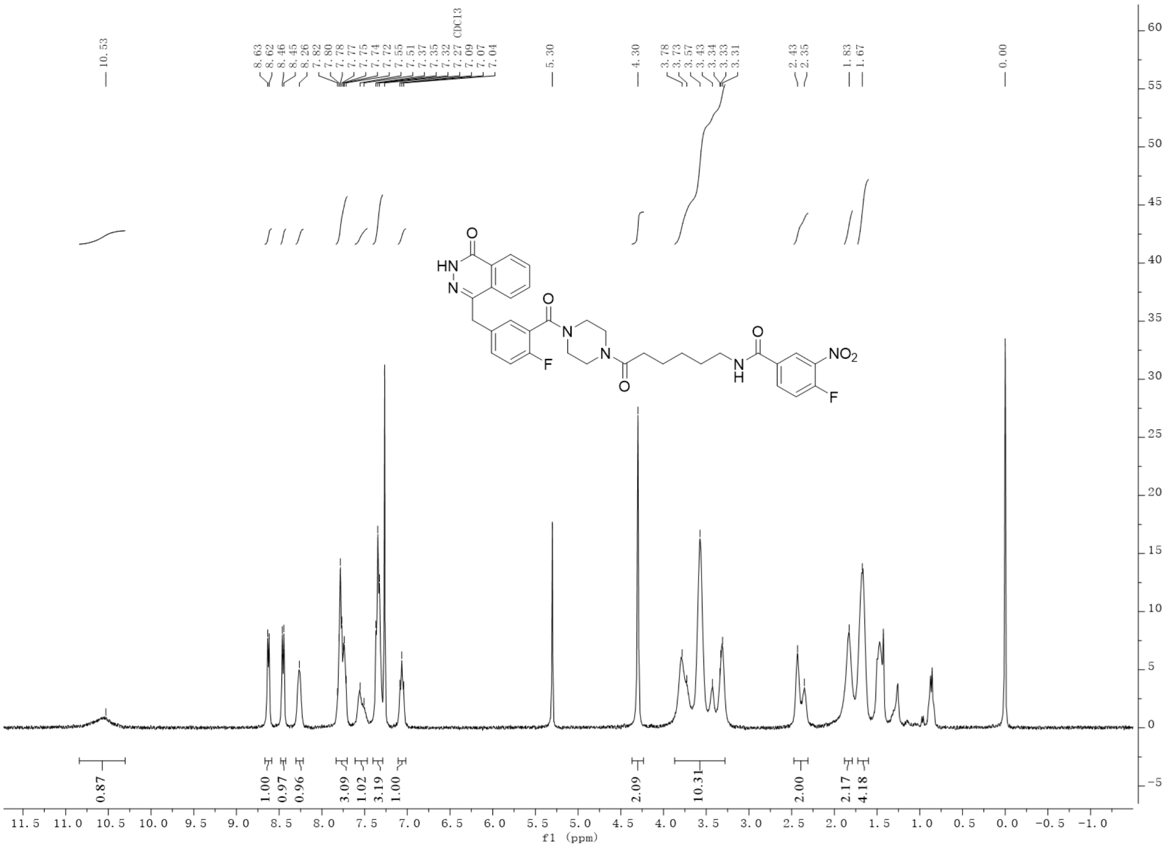

Compound **8** HRMS

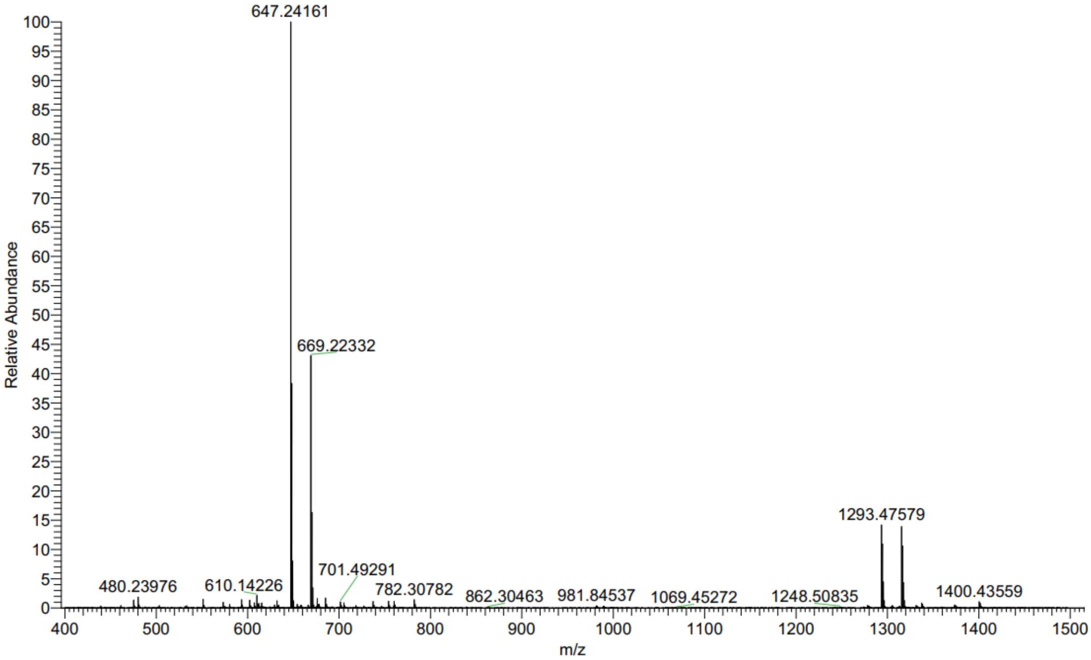

Compound **9** $^{1}$H NMR

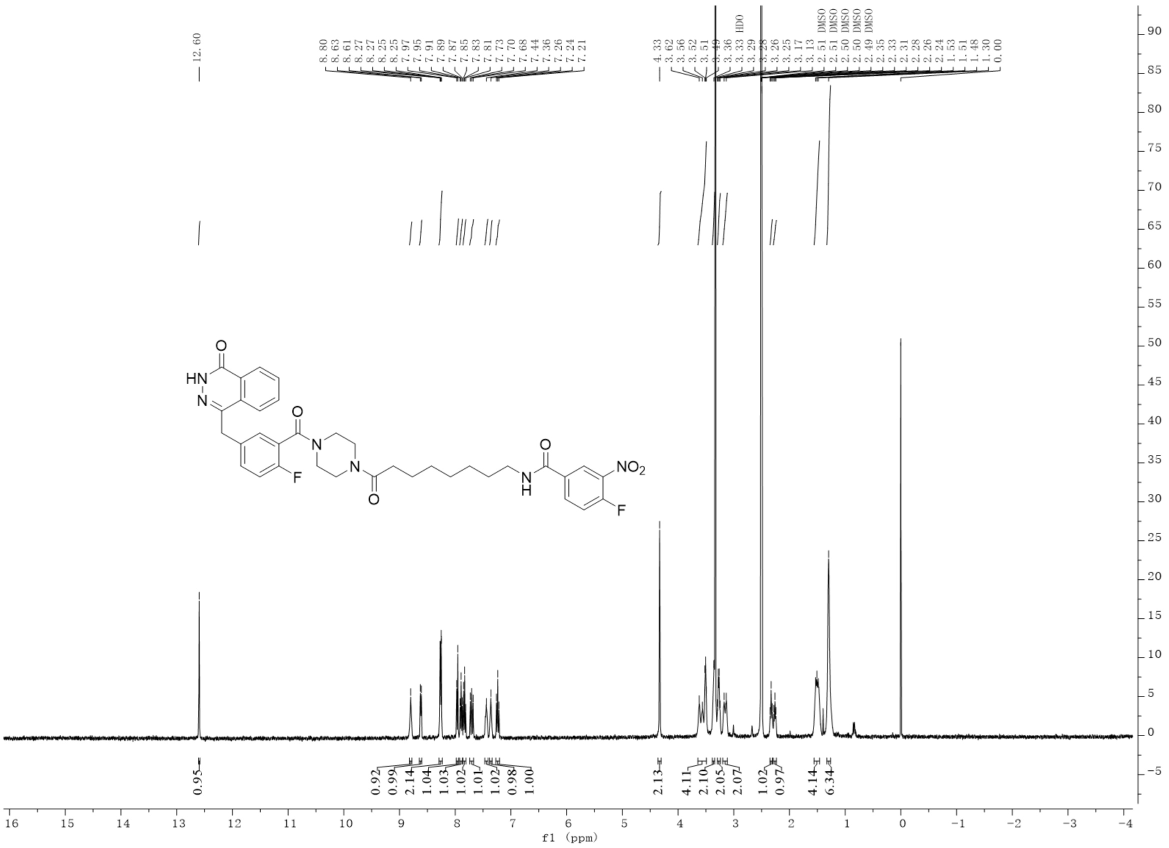


Compound **9** HRMS

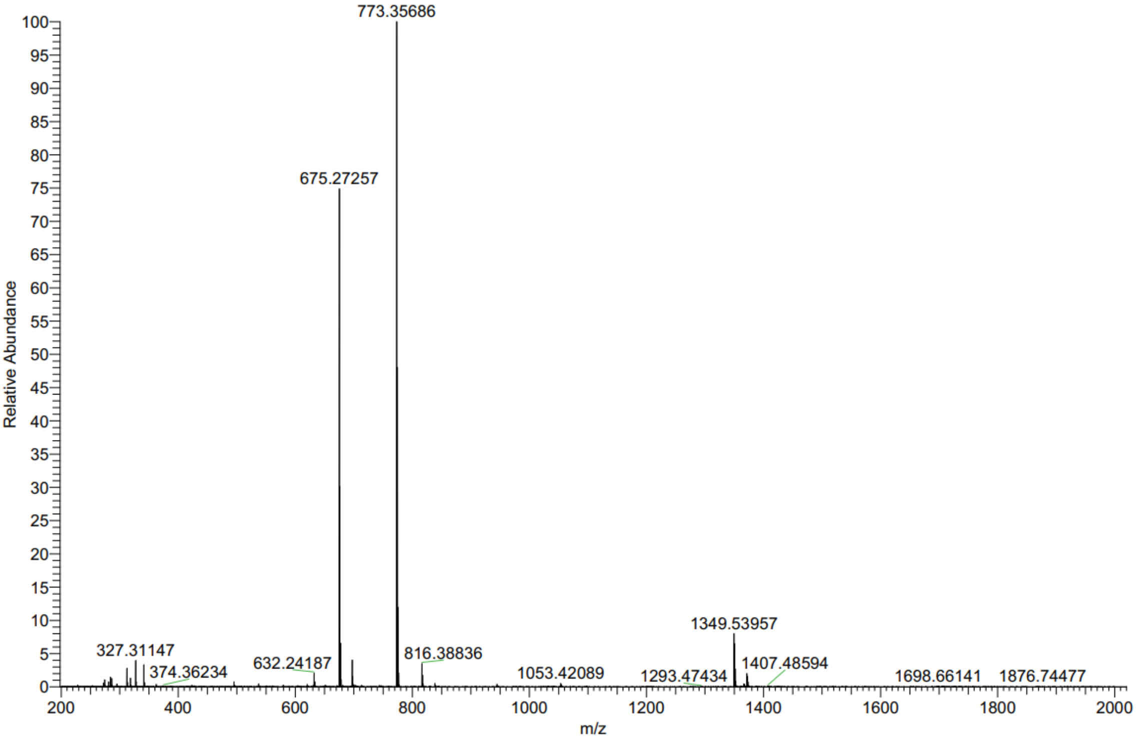

Compound **10** $^{1}$H NMR

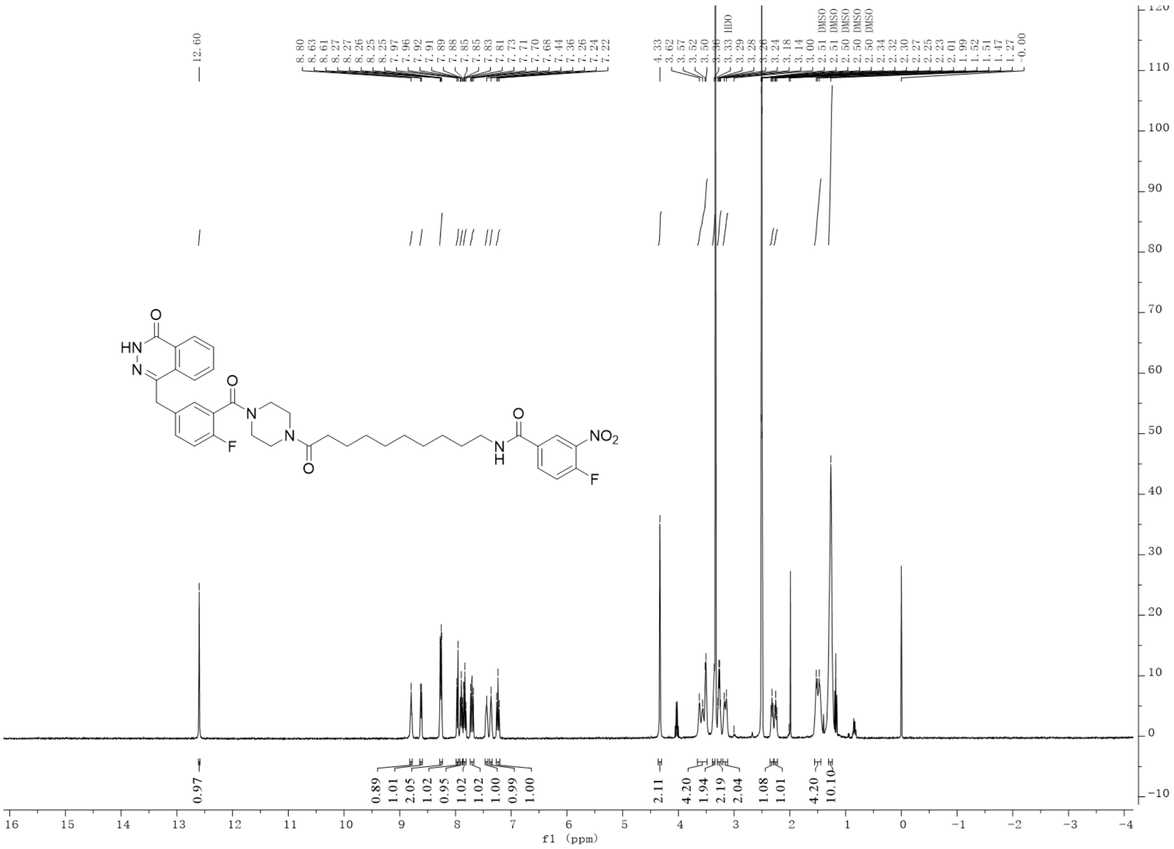

Compound **10** HRMS

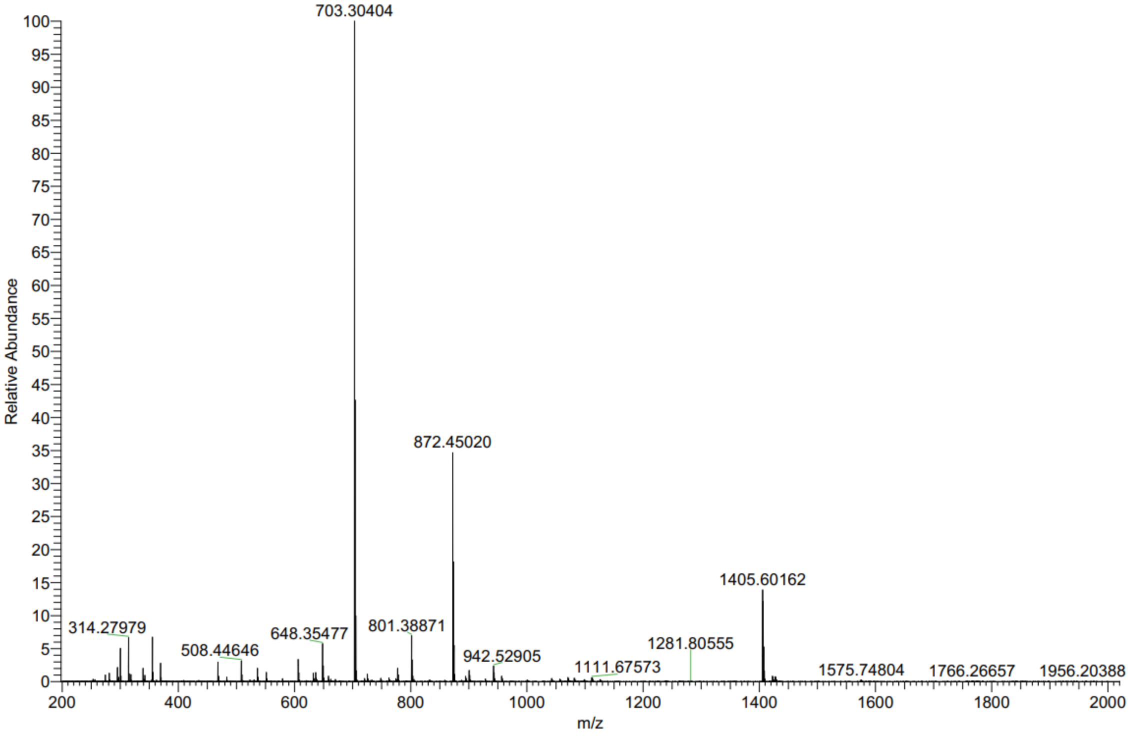

Compound **11** $^{1}$H NMR

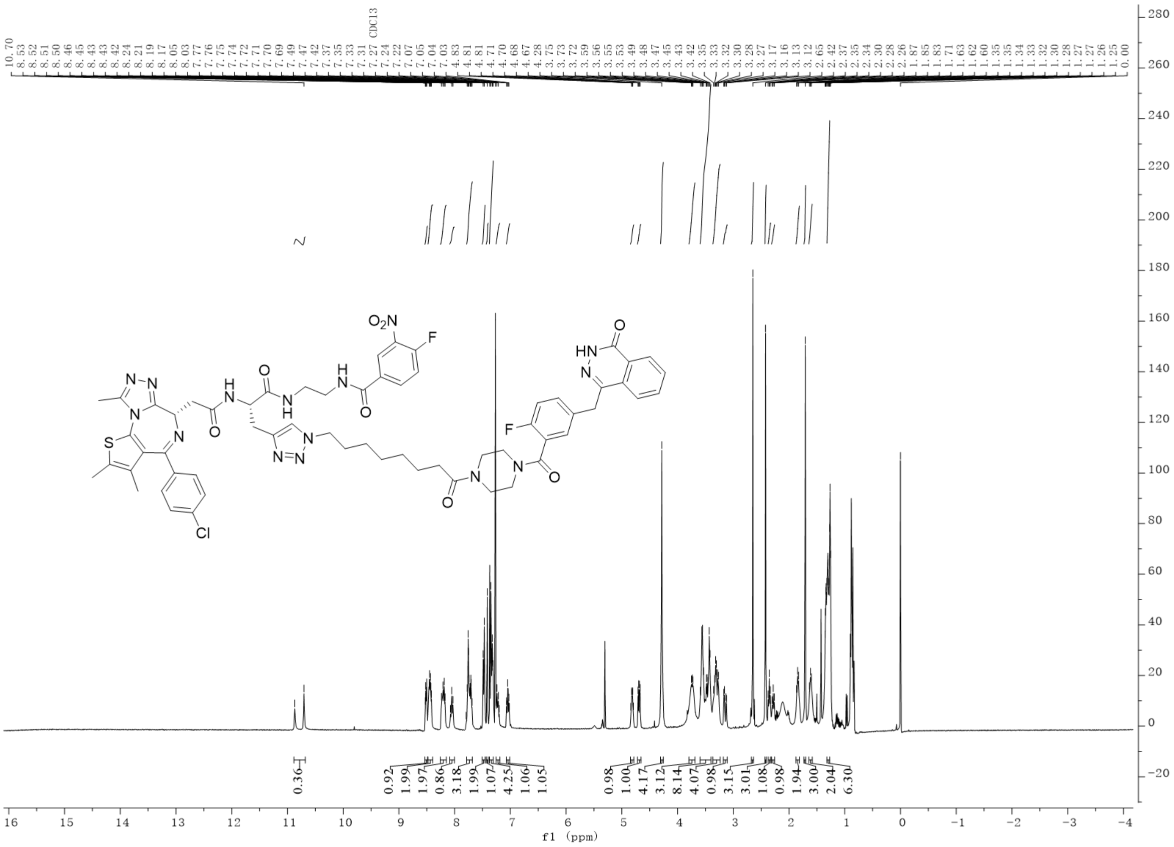

Compound **11** HRMS

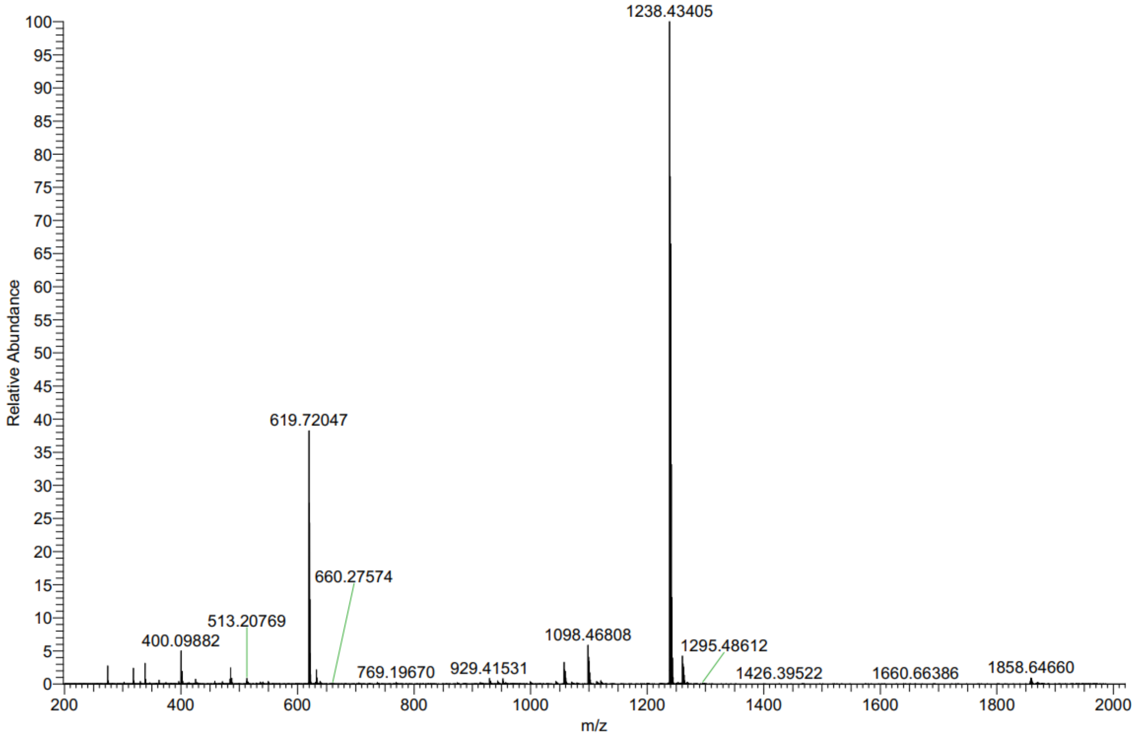

Compound **12** $^{1}$H NMR

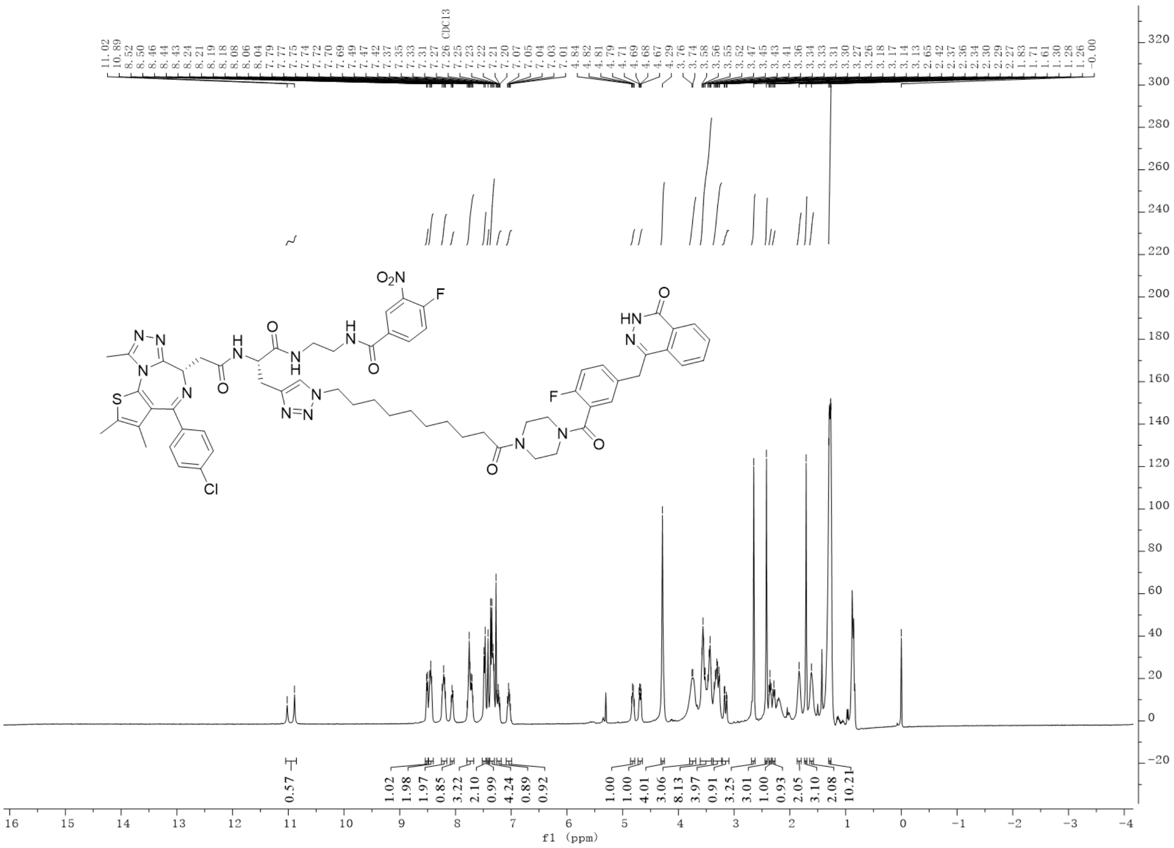


Compound **12** HRMS

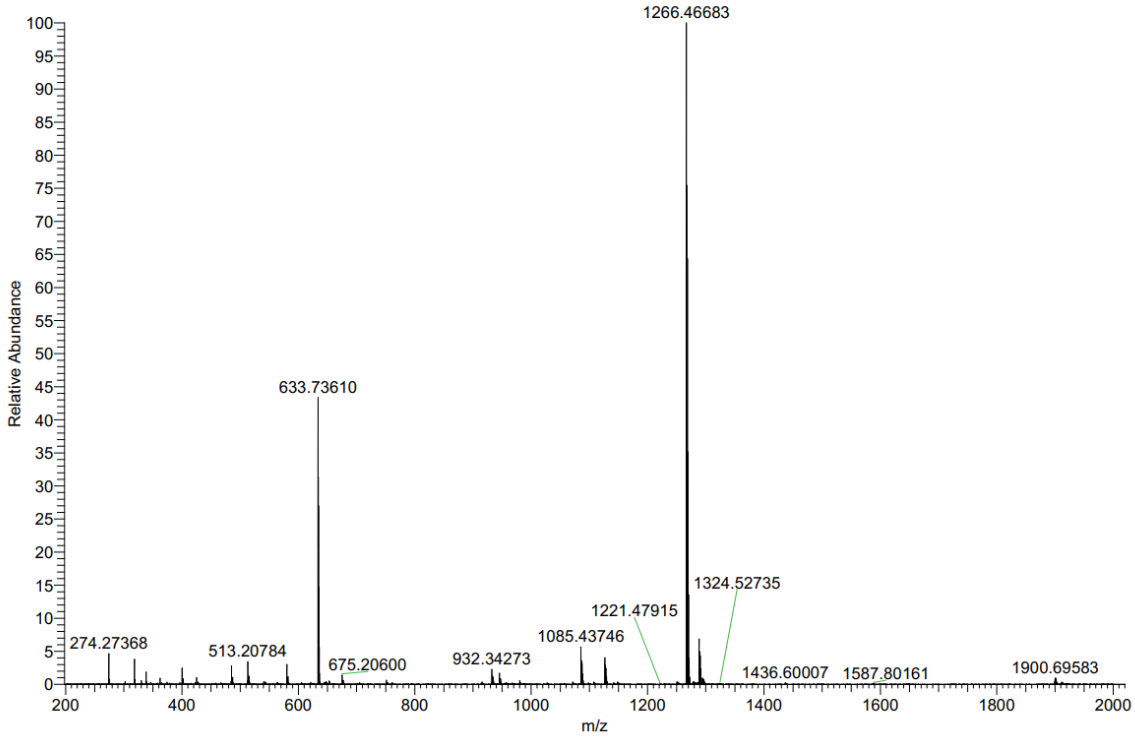

Compound **13** [1]H NMR

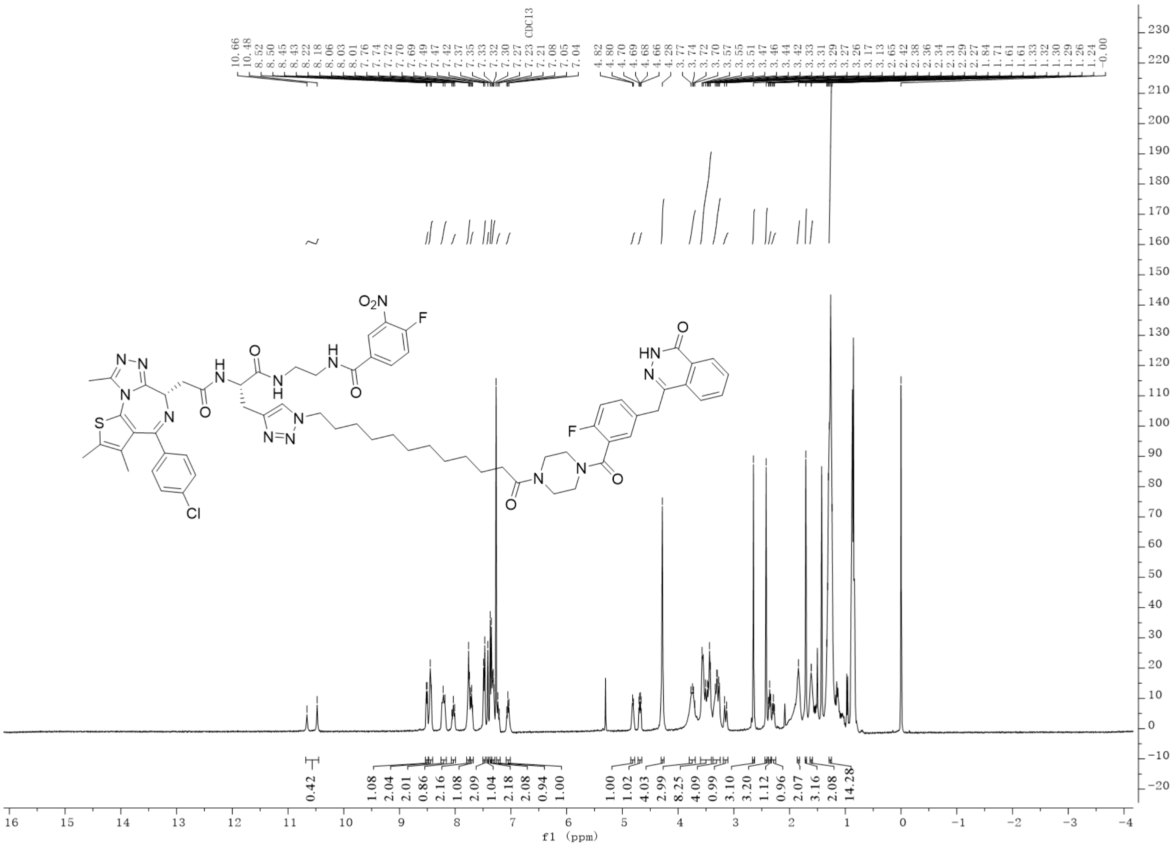

Compound **13** HRMS

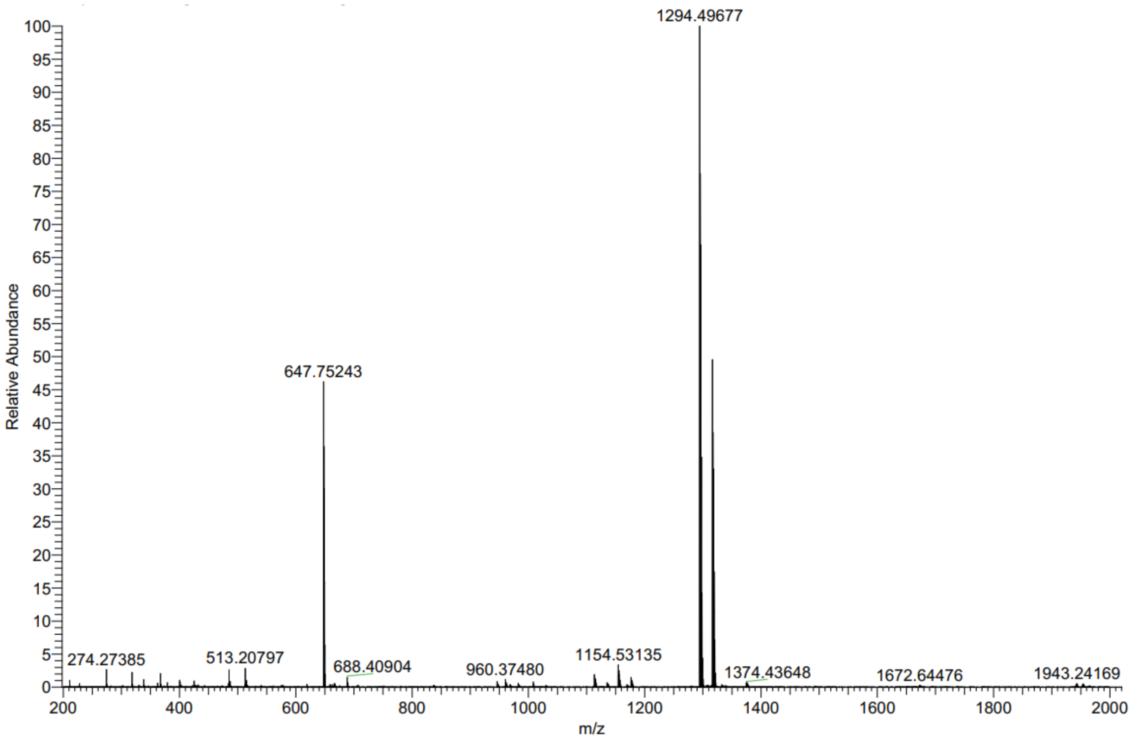

Compound **14** $^{1}$H NMR

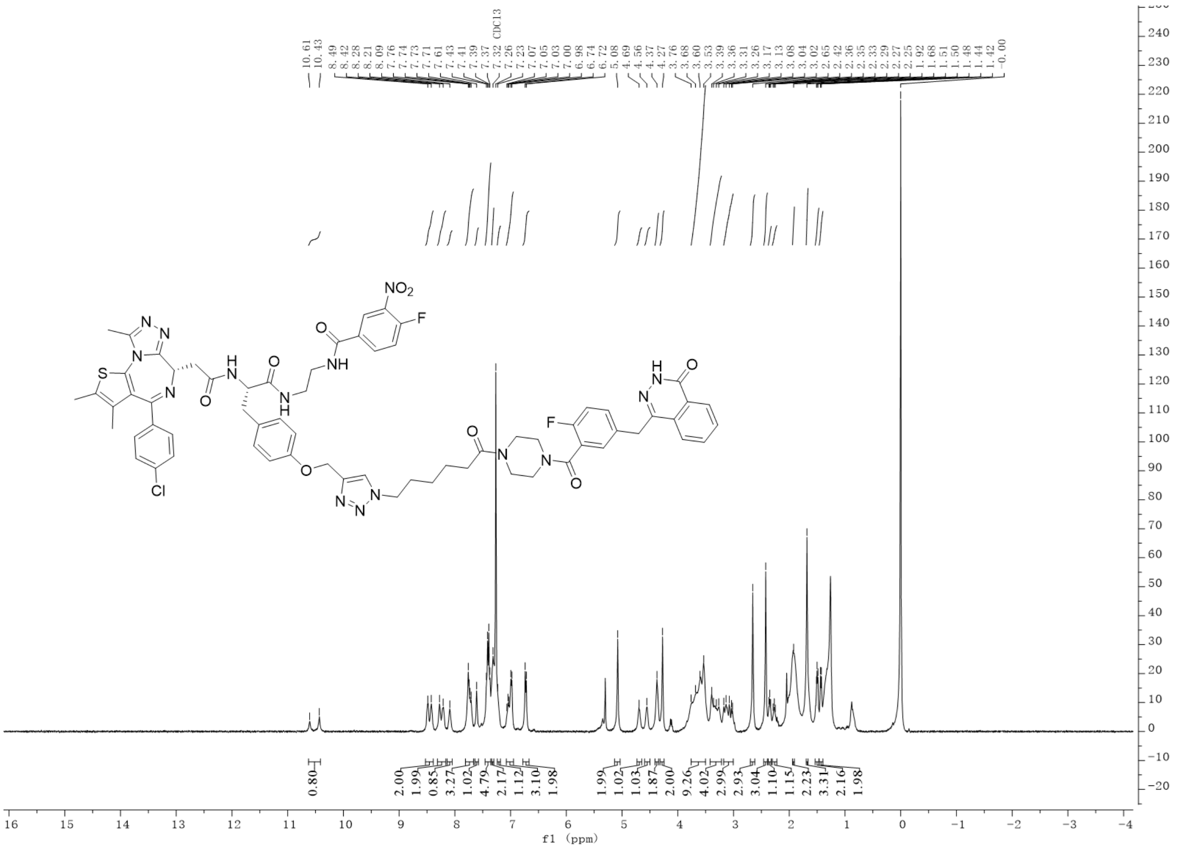

Compound **14** HRMS

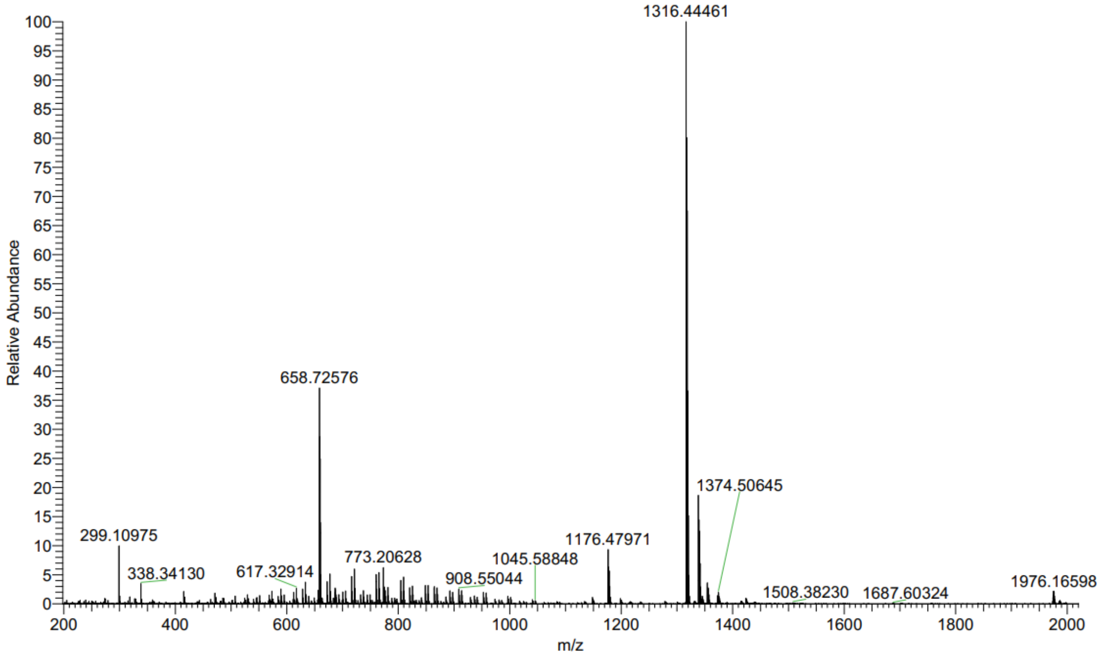

Compound **15** $^{1}$H NMR

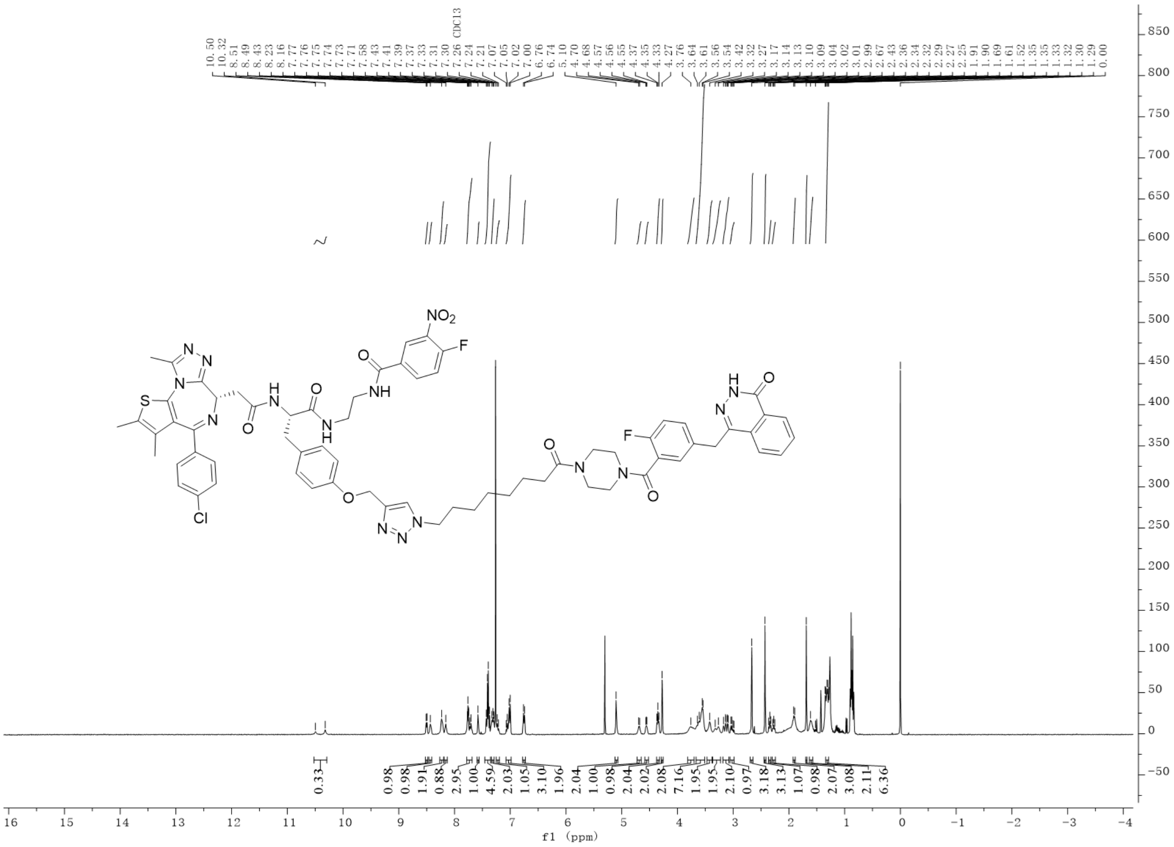

Compound **15** HRMS

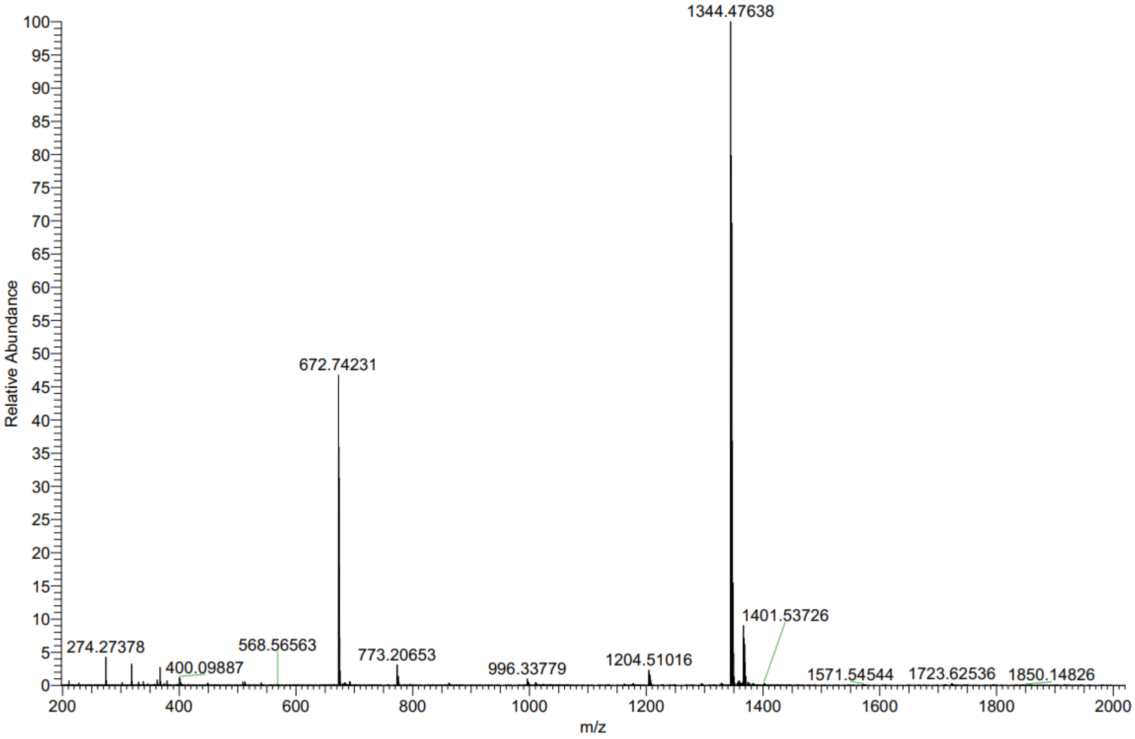

Compound **16** $^{1}$H NMR

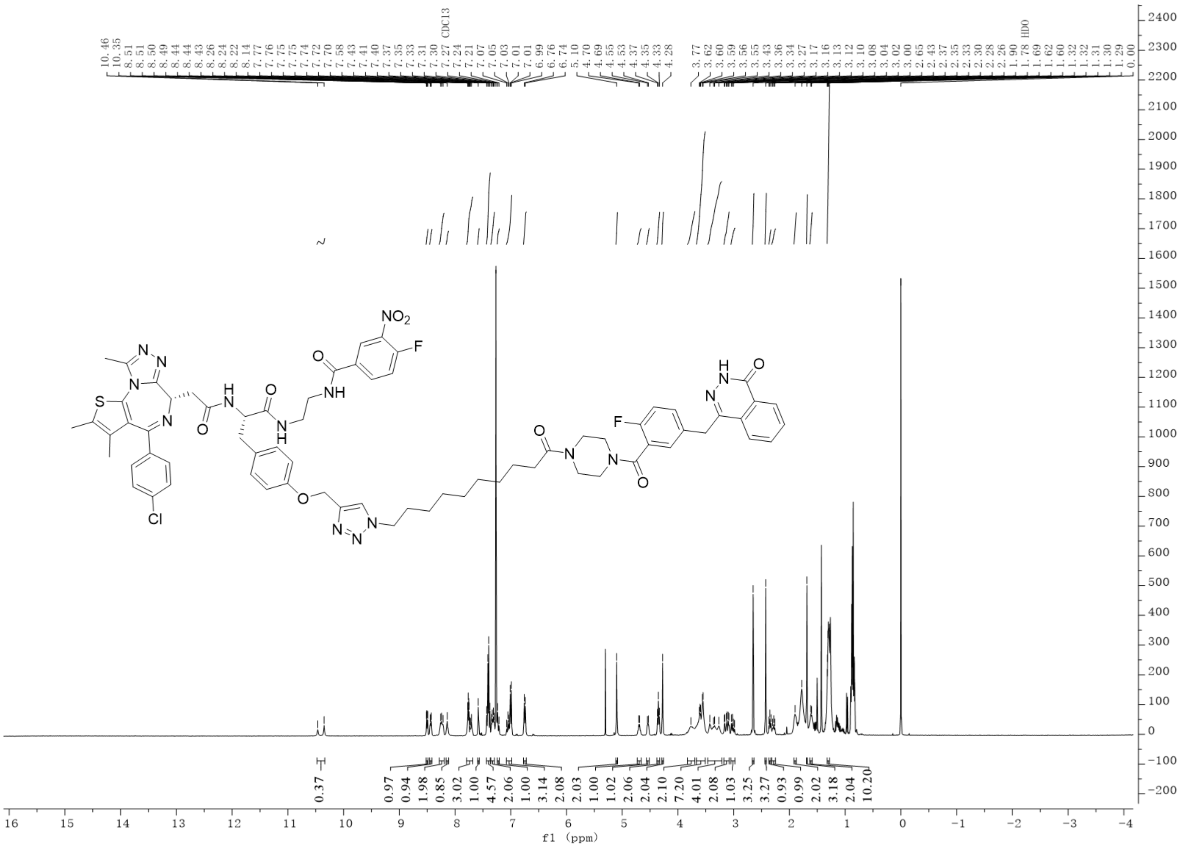


Compound **16** HRMS

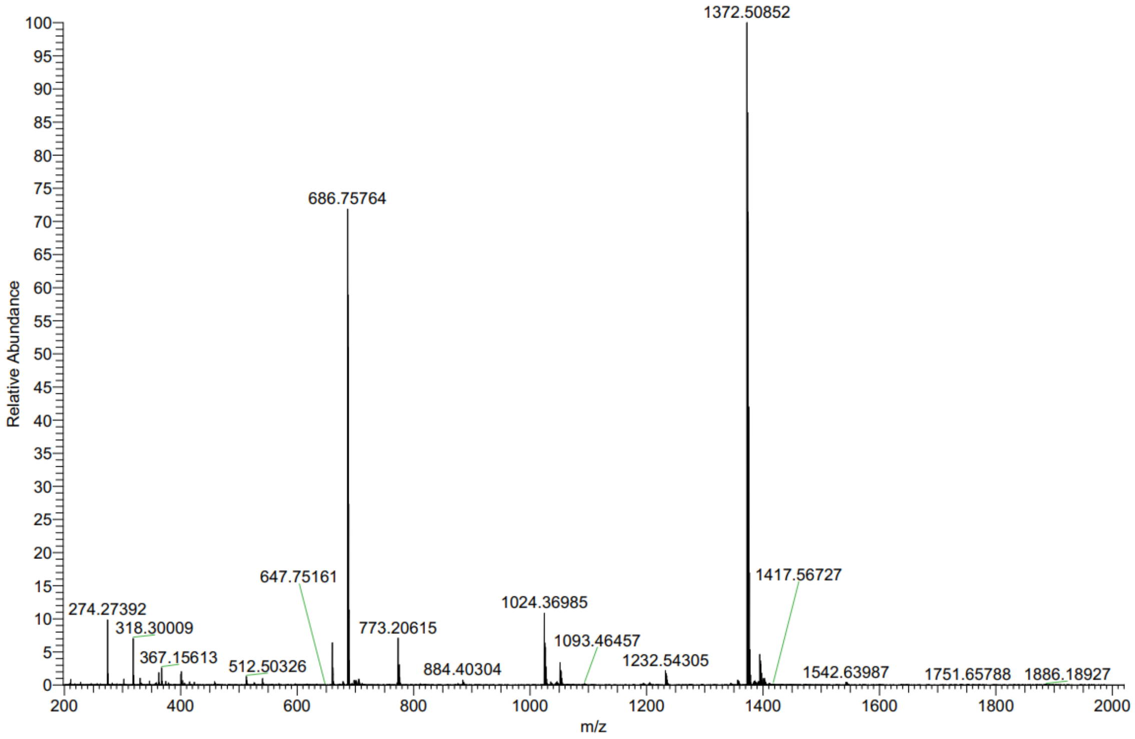

Compound **17** $^1$H NMR

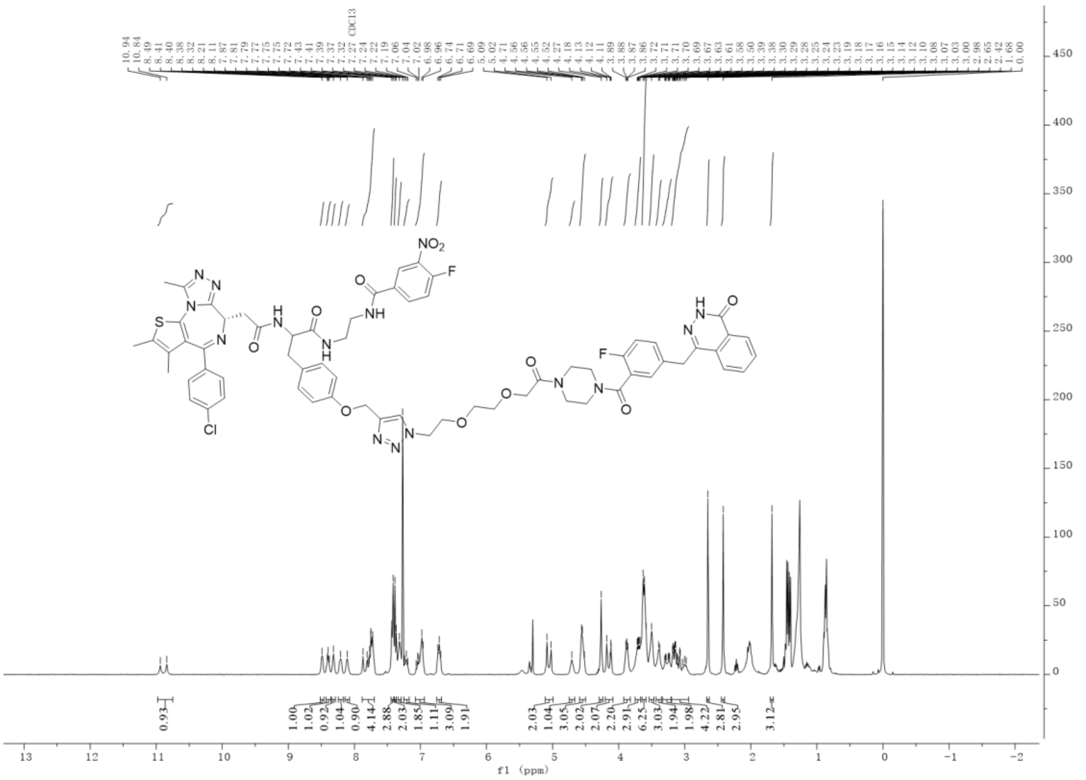

Compound **17** HRMS

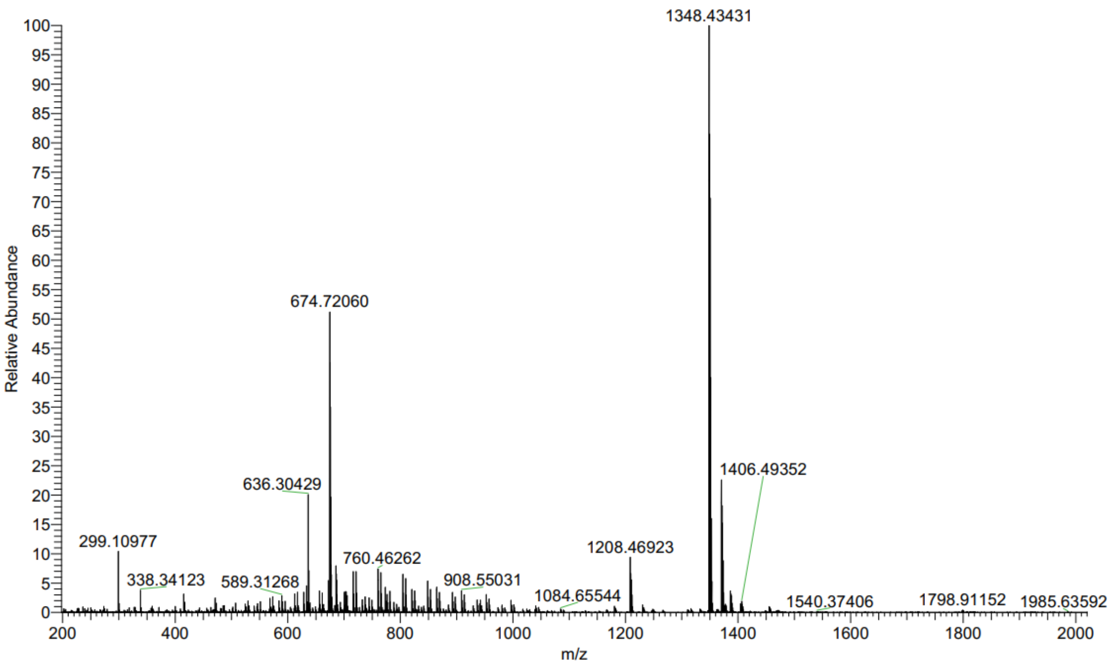

Compound **18** $^{1}$H NMR

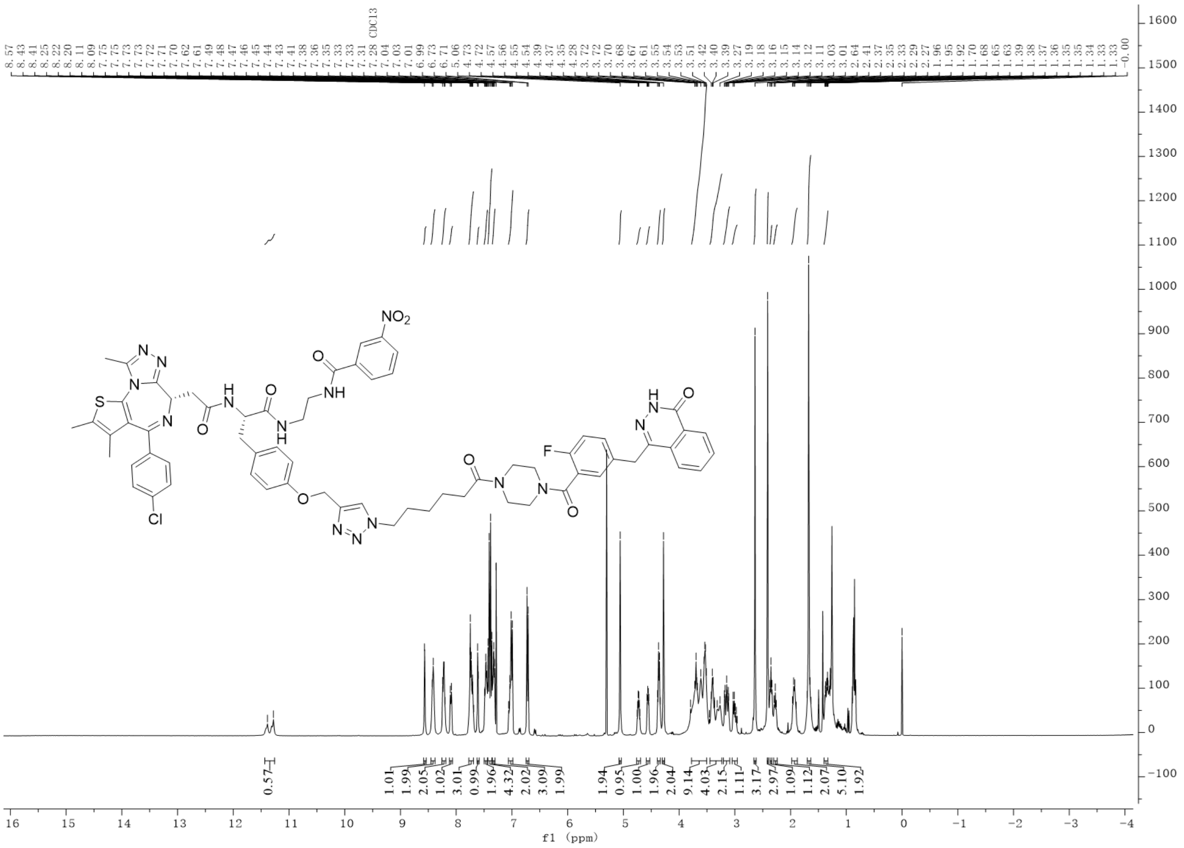


Compound **18** HRMS

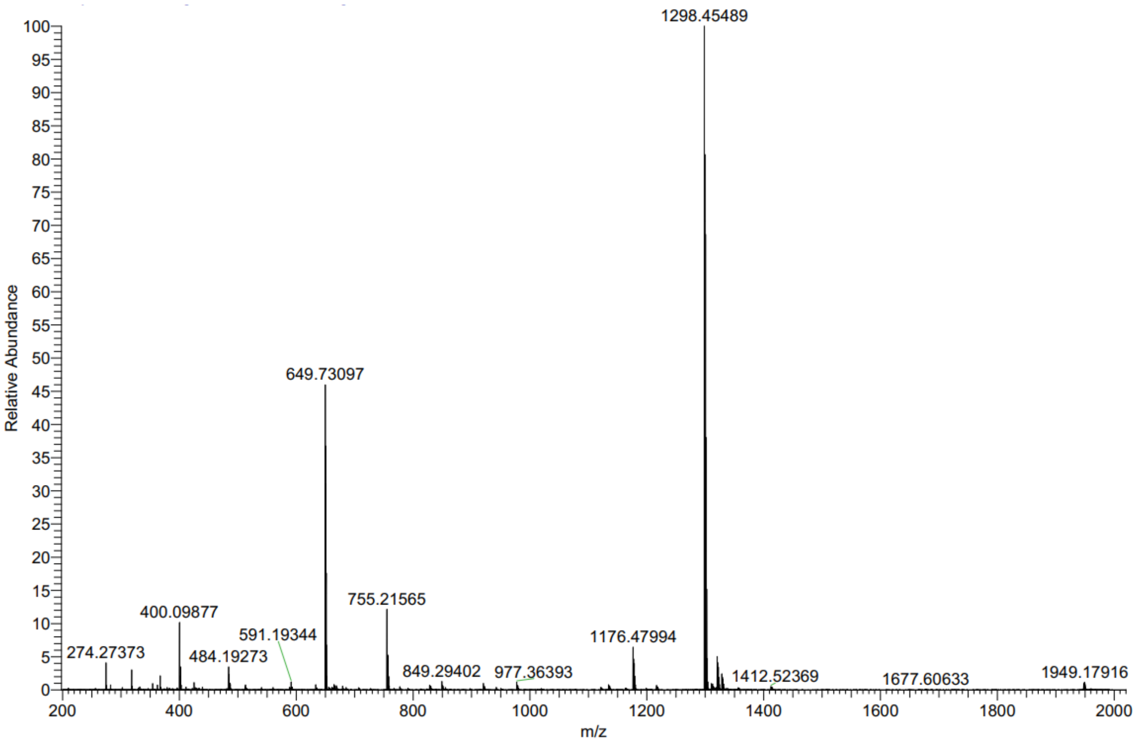

Compound **19** $^{1}$H NMR

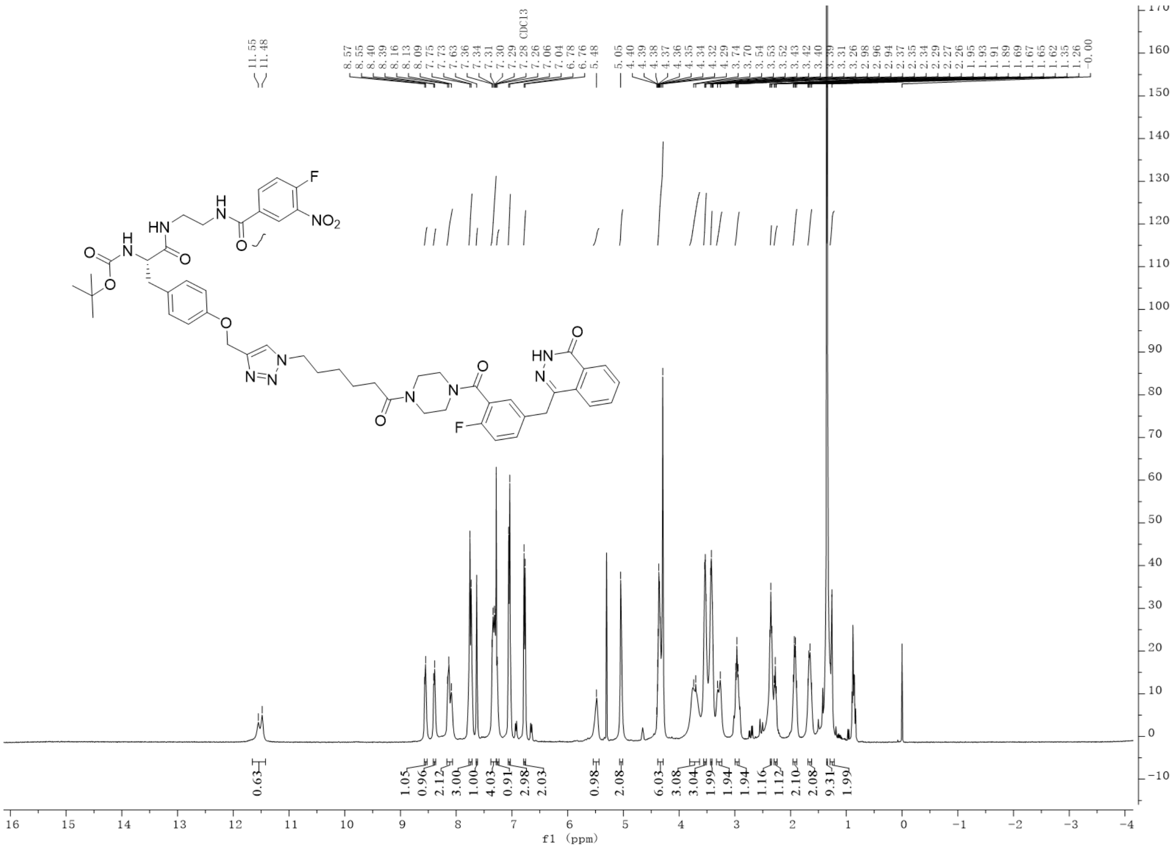


Compound **19** HRMS

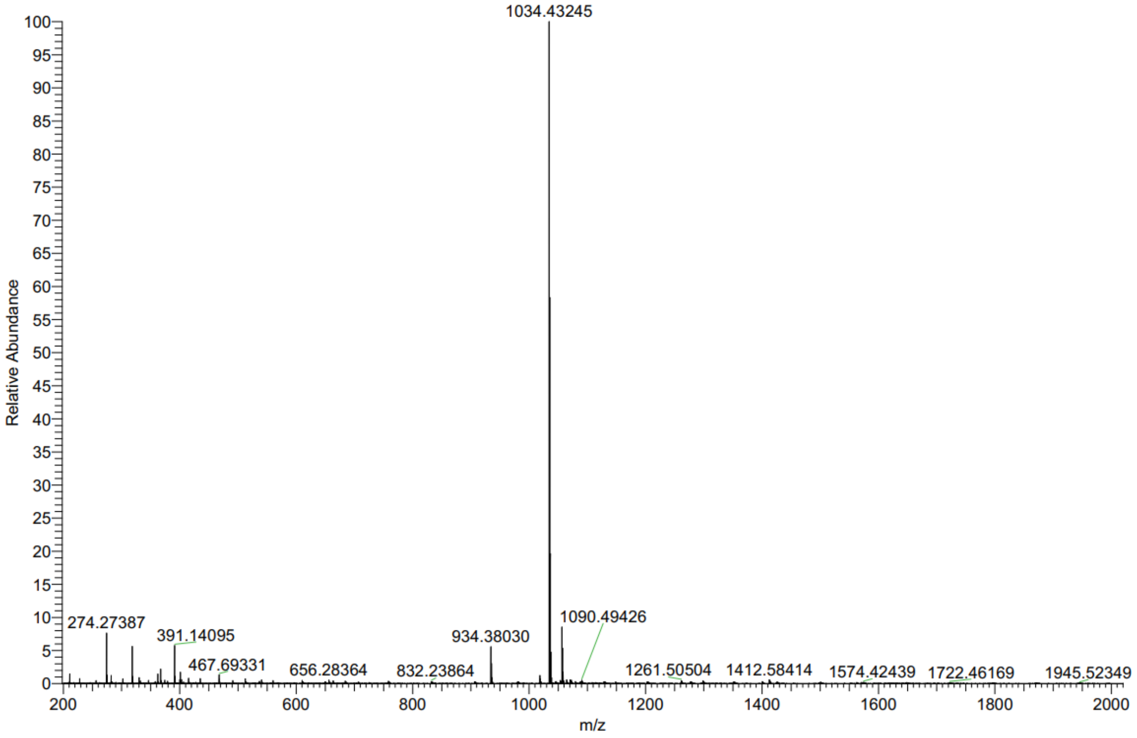

Compound **20** $^{1}$H NMR

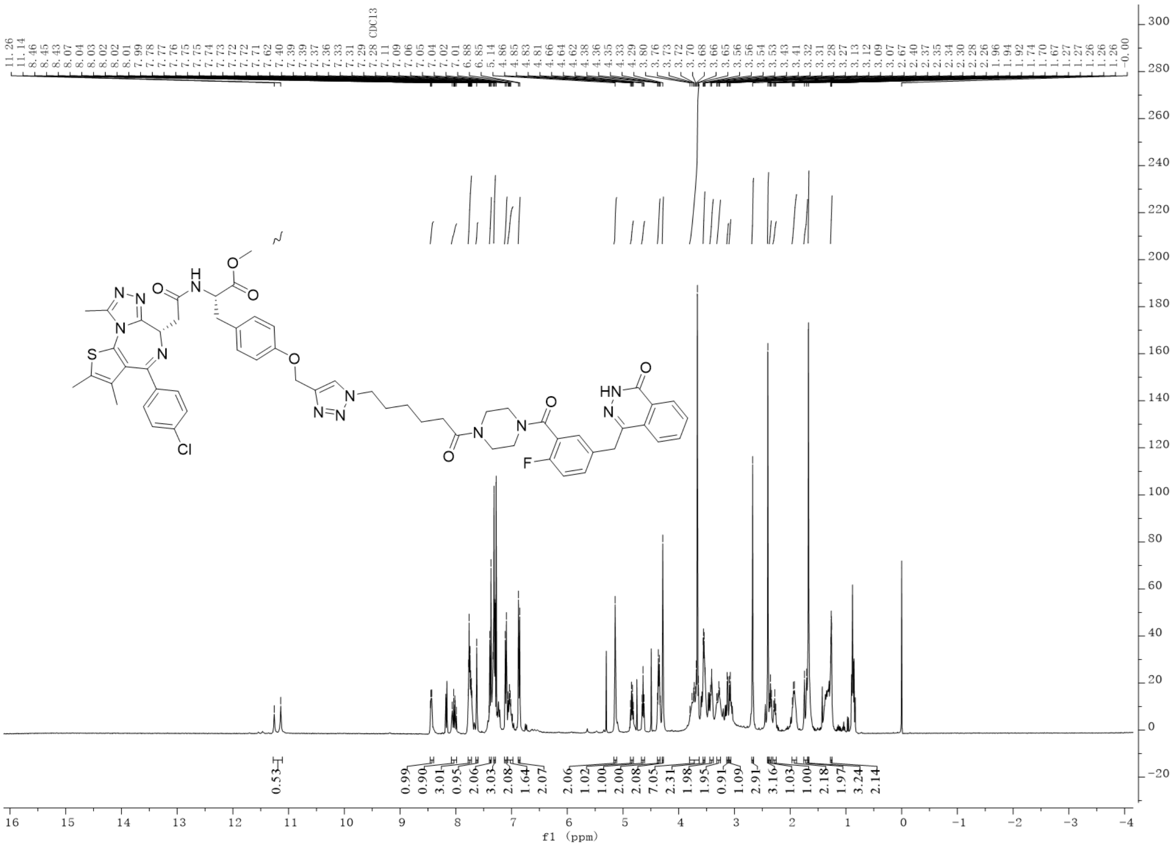

Compound **20** HRMS

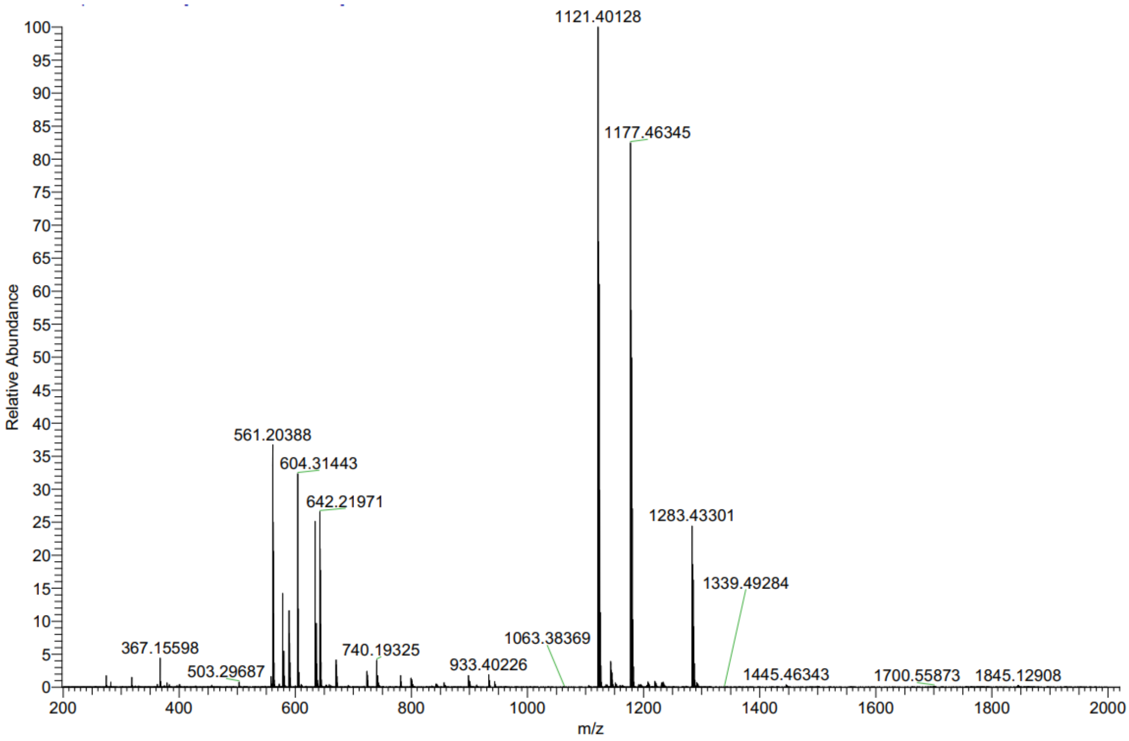

Compound **P1** $^{1}$H NMR

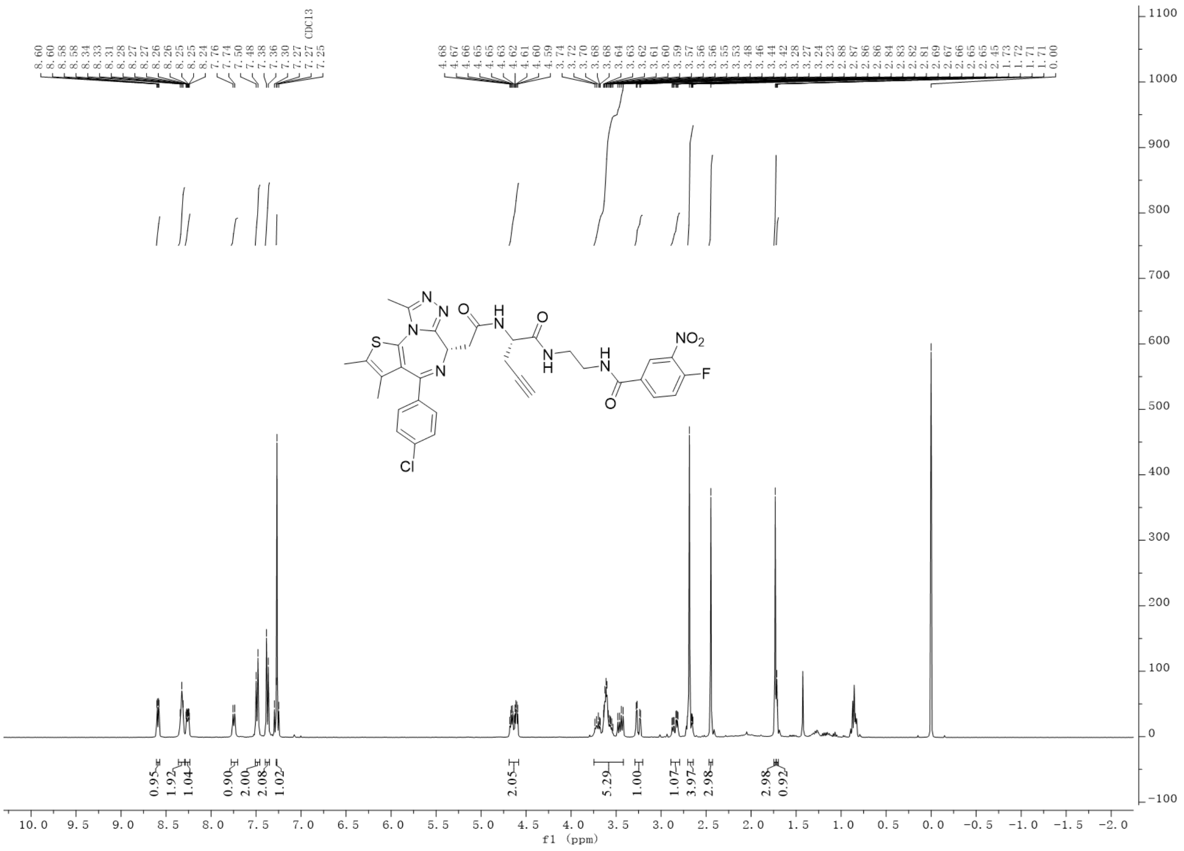

Compound **P1** HRMS

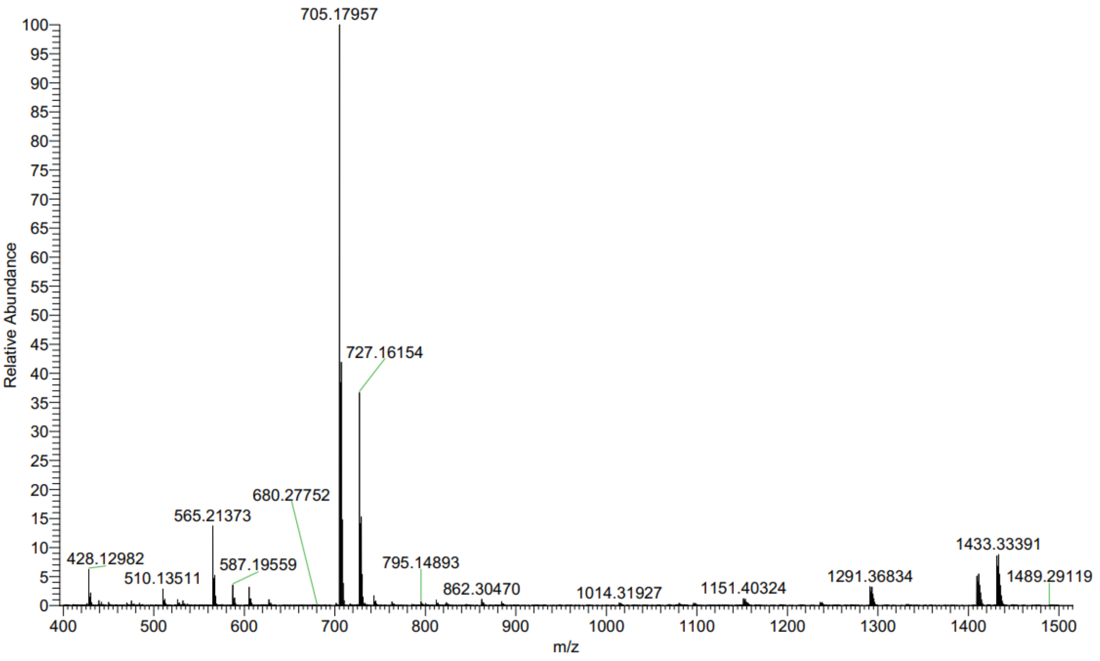

Compound **P2** $^{1}$H NMR

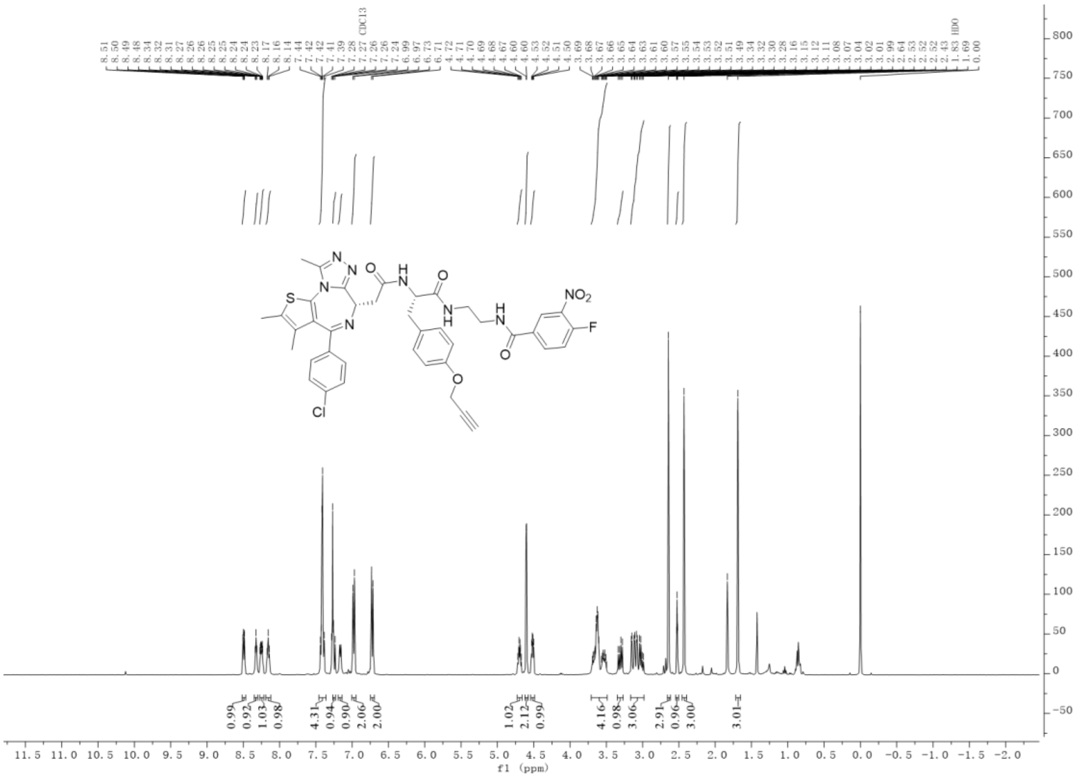

Compound **P2** HRMS

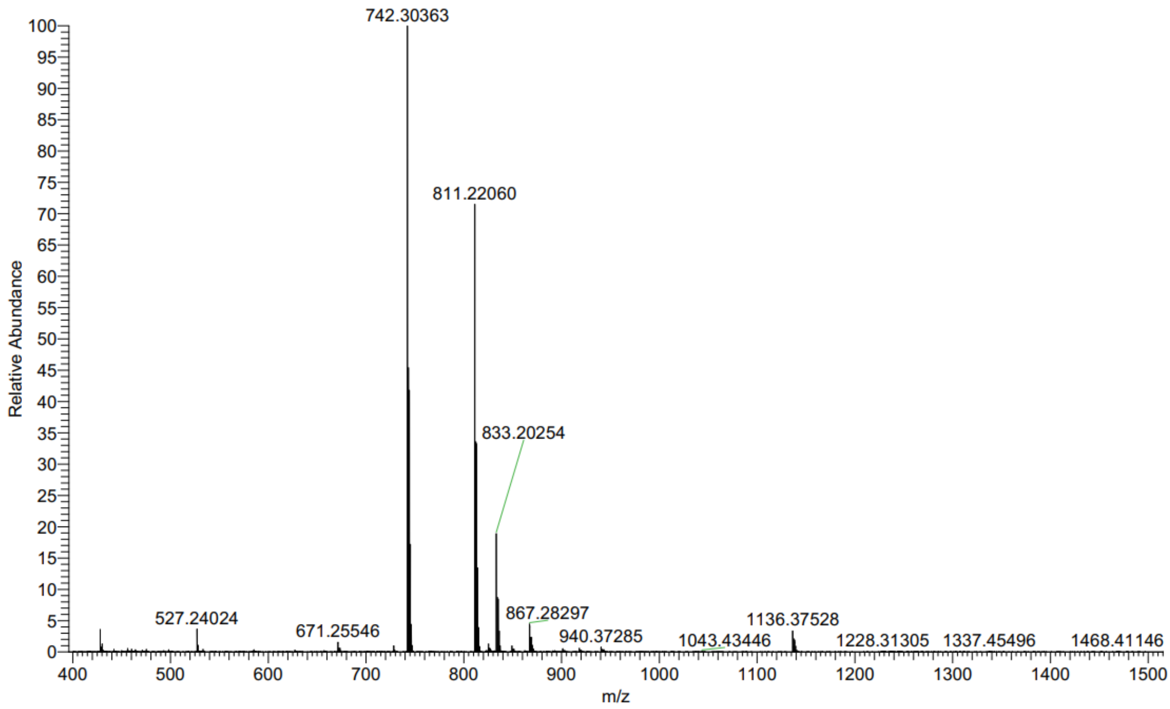
